# Assessing changes in global fire regimes

**DOI:** 10.1101/2023.02.07.527551

**Authors:** Sayedeh Sara Sayedi, Benjamin W Abbott, Boris Vannière, Bérangère Leys, Daniele Colombaroli, Graciela Gil Romera, Michał Słowiński, Julie C. Aleman, Olivier Blarquez, Angelica Feurdean, Kendrick Brown, Tuomas Aakala, Teija Alenius, Kathryn Allen, Maja Andric, Yves Bergeron, Siria Biagioni, Richard Bradshaw, Laurent Bremond, Elodie Brisset, Joseph Brooks, Sandra Bruegger, Thomas Brussel, Haidee Cadd, Eleonora Cagliero, Christopher Carcaillet, Vachel Carter, Filipe X. Catry, Antoine Champreux, Emeline Chaste, Raphaël Daniel Chavardès, Melissa Chipman, Marco Conedera, Simon Connor, Mark Constantine, Colin Courtney Mustaphi, Abraham N Dabengwa, William Daniels, Erik De Boer, Elisabeth Dietze, Joan Estrany, Paulo Fernandes, Walter Finsinger, Suzette Flantua, Paul Fox-Hughes, Dorian M Gaboriau, Eugenia M. Gayo, Martin.P Girardin, Jeffery Glenn, Ramesh Glückler, Catalina González-Arango, Mariangelica Groves, Rebecca Jenner Hamilton, Douglas Hamilton, Stijn Hantson, K. Anggi Hapsari, Mark Hardiman, Donna Hawthorne, Kira Hoffman, Virginia Iglesias, Jun Inoue, Allison T Karp, Patrik Krebs, Charuta Kulkarni, Niina Kuosmanen, Terri Lacourse, Marie-Pierre Ledru, Marion Lestienne, Colin Long, José Antonio López-Sáez, Nicholas Loughlin, Elizabeth Lynch, Mats Niklasson, Javier Madrigal, S. Yoshi Maezumi, Katarzyna Marcisz, Grant Meyer, Michela Mariani, David McWethy, Chiara Molinari, Encarni Montoya, Scott Mooney, Cesar Morales-Molino, Jesse Morris, Patrick Moss, Imma Oliveras, José Miguel Pereira, Gianni Boris Pezzatti, Nadine Pickarski, Roberta Pini, Vincent Robin, Emma Rehn, Cecile Remy, Damien Rius, Yanming Ruan, Natalia Rudaya, Jeremy Russell-Smith, Heikki Seppä, Lyudmila Shumilovskikh, William T. Sommers, Çağatay Tavşanoğlu, Charles Umbanhowar, Erickson Urquiaga, Dunia Urrego, Richard Vachula, Tuomo Wallenius, Chao You, Anne-Laure Daniau

## Abstract

Human activity has fundamentally altered wildfire on Earth, creating serious consequences for human health, global biodiversity, and climate change. However, it remains difficult to predict fire interactions with land use, management, and climate change, representing a serious knowledge gap and vulnerability. We used expert assessment to combine opinions about past and future fire regimes from 98 wildfire researchers. We asked for quantitative and qualitative assessments of the frequency, type, and implications of fire regime change from the beginning of the Holocene through the year 2300. Respondents indicated that direct human activity was already influencing wildfires locally since at least ^~^12,000 years BP, though natural climate variability remained the dominant driver of fire regime until around 5000 years BP. Responses showed a ten-fold increase in the rate of wildfire regime change during the last 250 years compared with the rest of the Holocene, corresponding first with the intensification and extensification of land use and later with anthropogenic climate change. Looking to the future, fire regimes were predicted to intensify, with increases in fire frequency, severity, and/or size in all biomes except grassland ecosystems. Fire regime showed quite different climate sensitivities across biomes, but the likelihood of fire regime change increased with higher greenhouse gas emission scenarios for all biomes. Biodiversity, carbon storage, and other ecosystem services were predicted to decrease for most biomes under higher emission scenarios. We present recommendations for adaptation and mitigation under emerging fire regimes, concluding that management options are seriously constrained under higher emission scenarios.

## Introduction

Human alteration of land cover and climate is reshaping wildfire on Earth (Bowman et al., 2020; Davis, 2021; T. M. Ellis et al., 2022; Pereira et al., 2022). Most terrestrial ecosystems have coevolved with fire over millions of years and many require periodic disturbance to maintain ecosystem structure and function (Bond et al., 2005; Harris et al., 2016). Yet, when fires exceed their historical patterns of intensity, extent, severity, seasonality, and frequency (hereafter *fire regime*; Figure.1a), they can deleteriously influence biodiversity (Feng et al., 2021; Kelly et al., 2020), climate (IPCC, 2021), and societies (Doerr & Santín, 2016; Johnston et al., 2021; Jones, 2017). In some regions, recent state changes in fire regime have reduced ecosystem services, including air quality, water availability, habitat, and ecosystem carbon storage (Collins et al., 2021; Crandall et al., 2021; McClure & Jaffe, 2018; Pausas & Keeley, 2019; Xie et al., 2022). These changes in disturbances can cause loss of life and property, degradation of health, acute risk to fire managers, emergency evacuations, and other socioeconomic impacts (Balch et al., 2020; Raymond et al., 2020).

**Figure 1.**
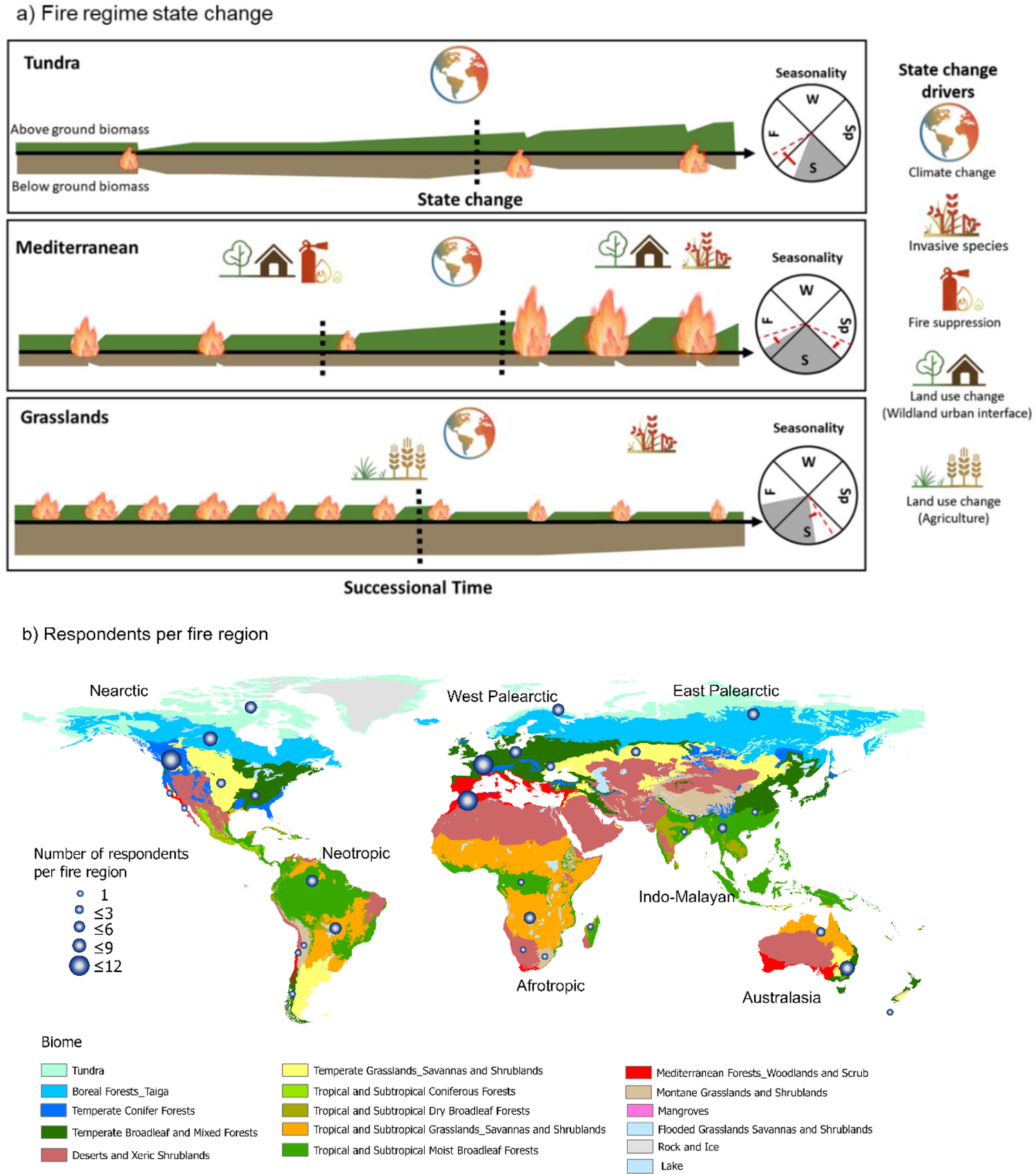
**a) Conceptual diagram of fire regime characteristics and state changes for three example biomes**. Fire regime is defined in terms of spatial (e.g., extent, type, patchiness), temporal (e.g., frequency, interval, seasonality), and physical (e.g., intensity, severity) fire characteristics. The size of flame in the Figure represents fire extent, and the vertical placement of the flame represents fire type (e.g., surface vs. crown). The green and brown bands represent above- and below-ground biomass, respectively. The vertical black dashed lines represent fire regime state change. The gray wedges represent fire seasonality before fire regime change: W: winter, Sp: Spring, S: summer and F: fall/autumn. Red dashed lines inside wedges represent new fire regime seasonality after state change **b) The location of the fire regions used in this study** (Olson 2001) with the number of respondents per fire region.

In the past and across large spatial scales, the dominant driver of fire regime has been the interaction between climate and vegetation (Abbott et al., 2016; Girardin et al., 2013; Harris et al., 2016; McDowell et al., 2020; Molinari et al., 2020). All aspects of climate, but especially patterns of precipitation and temperature influence vegetation composition and its moisture content. Climate and weather also control ignition sources, with lightning being the most common natural source of wildfire. Consequently, climate lays the foundation for fire regimes through fuel availability, flammability, and ignition likelihood (Bowman et al., 2009; Chen et al., 2021; Scholten et al., 2021). As humans modified global patterns of vegetation, ignition, and climate over the past several millennia (Abbott et al., 2019; E. C. Ellis et al., 2021; McDowell et al., 2020; Watson et al., 2018), fire disturbance became progressively more anthropogenically influenced across local to global scales (Figure. 1a). For example, humans have directly modified vegetation type and density for 77% of the Earth’s terrestrial surface, primarily through agriculture, with myriad consequences for fuel characteristics and ignition sources (Balch et al., 2017a; Bowman et al., 2011; Marlon et al., 2008; Słowiński et al., 2022; Watson et al., 2018). Understanding the characteristics and sensitivity of fire regime change is crucial for sustainable land management as well as climate change mitigation, adaptation, and planning (Cochrane & Bowman, 2021). However, knowledge of thresholds and tipping points in the relationships linking climate, land use, and fire regimes is lacking.

In this context, we conducted an assessment of scientific opinion about the drivers and consequences of fire regime change in the Holocene and near future. Combining assessments from multiple sources allows an integrative evaluation of the range of possible futures complementary to numerical model projections (Morgan, 2014; Sayedi et al., 2020; Schuur et al., 2013). These assessments can address the current needs of decision-makers and ecosystem managers to better understand and apply the consensus view from the research community. Using the collected informed opinion from experts, we evaluated centennial to millennial-scale state changes (Figure.1a) in past, present, and future fire regimes at both regional and biome levels through four questions: 1. How have fire regimes varied during the Holocene (the last ~11,700 years); 2. How likely are fire regime state changes under different future climate change scenarios; 3. What component of ecosystems will be affected by potential future fire regimes; and 4. What types of human activities could be the most effective for mitigation and adaptation under future fire regimes?

We asked experts from around the world (included here as co-authors) to respond to these questions for terrestrial ecosystems in seven biogeographic realms and 14 biomes (Figure1.b) that reflect global bioclimatic, socioeconomic, and fire regime characteristics (Olson et al., 2001). Each expert responded for a biome-scale fire regime within a biogeographic realm (we define this unit “a *fire region”*) of their expertise. We received responses for 70% of flammable land area worldwide (total land surface excluding rock, ice, and lakes;Figure1.b, S1, Table S2).

## Methods and materials

We used expert assessment to evaluate the risk of fire regime change and its consequences in the future. After a literature review of both scientific and policy related documents about wildfires and fire regime change, we designed a questionnaire to gather scientific opinion on changes in fire regimes and their effects on ecosystems, climate, and societies (see Supplementary Information). The focus was specifically on centennial-to-millennial changes in past, present and future fire regimes by biogeographic realms and biome to consider long-term processes beyond observational and instrumental records. After two pretesting rounds, the finalized questionnaire was distributed to 430 scientists with fire related expertise. The list of experts was developed based on publications and referees from respondents. We received 123 filled surveys from 98 respondents (46% female, 45% male, 9% unspecified) (Table S2) from 23 countries (Figure.S24). The primary research disciplines as self-identified by the respondents were 55% paleoecology, 17% ecology, and 28% other fields such as geography or geosciences.

The questionnaire included background information on the concept of fire regime and state change to delineate a standardized definition for this study for experts to use and a description of RCP scenarios and predicted temperature and precipitation (see Supplementary Information). The question section focused on four key topics: 1) past fire regimes, 2) current fire regime states, 3) future fire projections, and 4) interventions and management. For each section, a self-reported expertise, confidence level, sources used to generate estimates (e.g., published or unpublished empirical data, professional opinion) (Table.S3), along with a list of sources of uncertainty were provided (Table.1). The questionnaire was a combination of quantitative and qualitative questions (full questionnaire included in the Supplementary Information). The qualitative questions included both open-ended questions and numerical responses. The open-ended questions were analyzed using thematic analysis method by coding the responses into categories after reviewing the responses.

For some of the quantitative questions, three quantiles (5% lower, 50% central and 95% upper) were provided to build a credible range of 90% for each question. We primarily used median central estimates for the main manuscript; though full ranges are shown in the Supplementary Information. We used nonparametric boxplots to evaluate the range of responses for each question, with descriptive statistics calculated in R version 3.6.1. We used ArcGIS Pro 3.0 for spatial analyses and visualizations.

## Results and discussion

In the following subsections, we present estimates and suggestions based on experts’ responses and we compare these opinions of estimates and projections with relevant literature. We focus on general patterns and trends among biomes and biogeographic realms (the comprehensive expert responses by fire region are available in the Supplementary Information). For the purposes of our study, we defined state change broadly as a large and sustained departure from a set of specific system behaviors (details in Supplementary Information-questionnaire). Fire regime state changes can be triggered by pulse or press disturbances, including internal and external drivers (e.g. climate change, vegetation shifts, human behavior), and can be reversible or permanent. For example, a state change in fire regime may be expressed as a shift in the central tendency (a decrease in mean annual area burned), overall variance (an increase in inter-annual variability of area burned), or frequency of events that exceed an ecological threshold (a change in the return interval of crown fires)(Scheffer et al., 2009; Seddon et al., 2014).

### Past to present drivers of fire regime

The median estimated number of fire regime state changes during the Holocene ranged from seven (*Temperate grasslands, savannas, and shrublands*) to two (*Tundra*) across biomes (Figure. S2). Regarding the timing of the three largest fire regime state changes, 16% of respondents identified the early Holocene (ca. 11,700–8,200 BP), 27% the mid Holocene (ca. 8,200–4,200 BP), and 57% the late Holocene (ca. 4,200–0 BP). A temporal acceleration of state change occurred after the Industrial Revolution (1760 A.D.), which included 20% of the fire regime changes (Figure. S3). This represents a ^~^10-fold increase in the rate of fire regime changes over the last 250 years. The *Nearctic* and *Australasia* showed even stronger recent changes in fire regime, with 30% and 36% of the identified fire regime changes occurring in the past 250 years, respectively.

Climate was identified as the main driver of fire regime changes during the Holocene (47% of responses), especially in the early and mid-Holocene. Direct human activity was the second most identified driver (32%). However, the onset of strong human influence on fire regime occurred at different times in different regions (Figure. 2), with the greatest influence during the late Holocene. However, for the post-industrial period (1950 A.D.-present), climate change and direct human activity were mentioned equally often (40% and 46%, respectively). Vegetation and fuel were the least mentioned drivers for each time interval (Figure. S4), likely because these factors respond to climate on centennial timescales, illustrating the importance of temporal scale when considering drivers.

**Figure 2.**
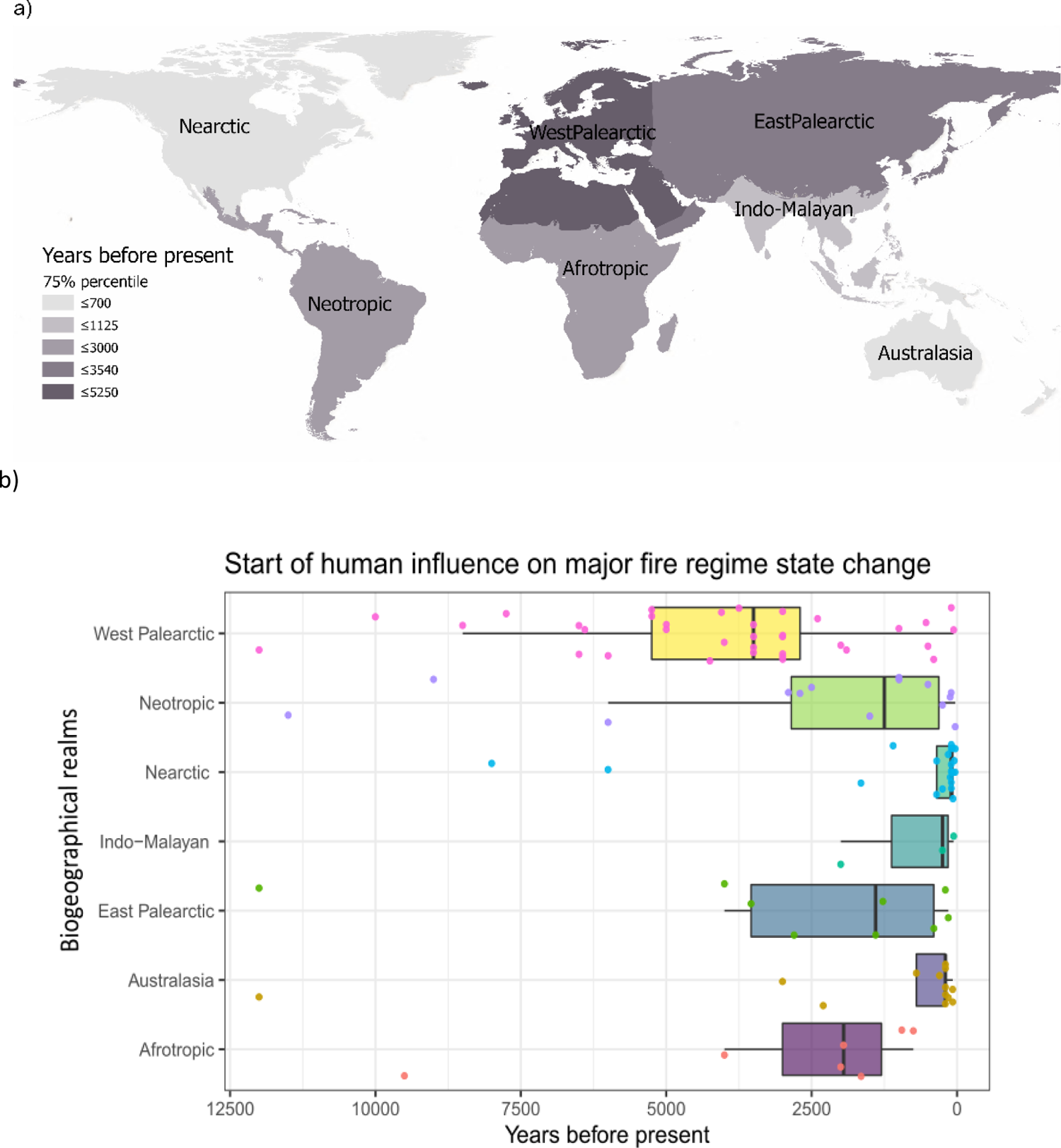
Estimates of when the earliest human-driven fire regime state changes occurred during the Holocene as estimated by respondents. a) 75^th^ percentile values of the earliest time humans were identified as a major driver of fire regime change by respondents. b) Points represent individual responses, while box plots represent the median, quartiles, most extreme points within 1.5-times the interquartile range (IQR), and points beyond 1.5times the IQR.

Respondents identified several dimensions of fire regime altered over the past 250 years, including fire frequency, extent, and severity (Figure. S5). A wide range of human-wildfire interactions specific to fire regions were identified. For example, in *Indo-Malayan Tropical forests*, deforestation due to economic development has changed the fuel structure and ignition sources, increasing fire activity in an ecosystem where it was historically rare. Other fire management strategies such as increased fire suppression and exclusion of Indigenous or traditional prescribed burning practices were recognized as drivers of increased fire severity, especially when coupled with recent temperature increases. In seven biomes it was suggested that a change in fire regime occurred since the industrial revolution (Figure. S3), with the median estimate of current fire regime duration less than 200 years (Figure.S6). For *Tundra, Mediterranean forests, woodlands, and scrub; Tropical and subtropical moist broadleaf forests, and Tropical and subtropical grasslands, savannas, and shrubs*, this duration was less than 70 years (Figure. S6).

### Timing and type of future fire regime change

Respondents provided estimates of fire regime change for their fire region in 2050, 2100, and 2300 based on the IPCC representative concentration pathways (RCP) 2.6, 4.5, and 8.5, representing increasingly severe climate change scenarios. Most respondents projected that the likelihood of fire regime change increased with time and climate change severity (Figure. S9-10). Under RCP8.5, nine biomes were projected to have more than 50% chance of experiencing a fire regime change by 2050, compared to one biome for RCP2.6. By 2100 and 2300 even under RCP2.6, five biomes were predicted to have a ≥50% likelihood of fire regime change (Figures. 3a, S9-10). The climate sensitivity (amount of increase in the likelihood of a fire regime change across RCP scenarios) varied substantially among biomes. For example, for *Tundra*, RCP2.6 was enough to initiate a fire regime change, while the projected likelihood of fire regime change was much lower under RCP2.6 than RCP8.5 for *Boreal forests* (Figure.3b, S11-12). Analysis for all scales, years, and scenarios are presented in the Supplementary Inforamtion.

**Figure 3.**
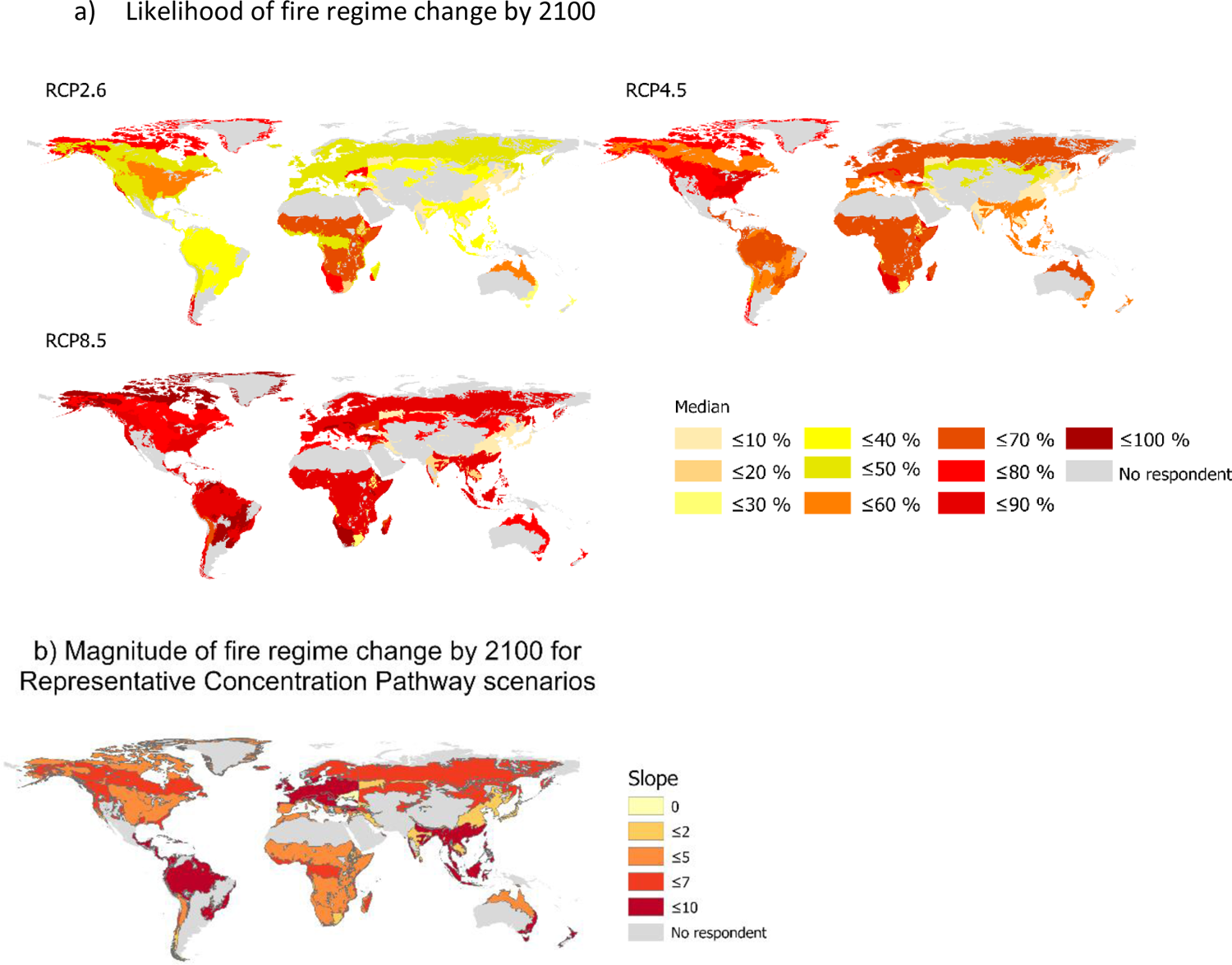
Likelihood of fire regime change under different RCP scenarios as estimated by respondents. **a)** Median of central values of the likelihood of fire regime change for year 2100 under three RCP scenarios. **b)** Magnitude of fire regime change for each biome was based on the slope between the estimated likelihood of a fire regime change (%) and the amount of radiative forcing for the three RCP scenarios (W.m^-2^). The higher values represent a higher magnitude of fire regime likelihood change moving from RCP2.6 to RCP8.5.

The climate sensitivity estimates from this study were in agreement with many modeling studies projecting future changes in fire activity (Bowman et al., 2020). Climate drivers such as *fire weather* and *fire danger days* are predicted to increase in many areas of the globe (IPCC, 2021), particularly in fire-prone regions such as the Mediterranean basin, southwestern USA, and subtropical regions of the Southern Hemisphere (Bowman et al., 2017; Cook et al., 2022). An increase in extreme *fire behavior* is also predicted in many regions such as the Amazon, western USA, Mediterranean and southern Australia (Bowman et al., 2020). Substantial intensification of fire behavior is projected for higher latitudes through the end of the 21^st^ century (Abbott et al., 2021; Flannigan et al., 2013; Talucci et al., 2022), though local fire patterns are expected to be heterogeneous (McCarty et al., 2021).

Respondents projected an intensification of fire regime across biomes, with burned area, frequency, and severity increasing for all but a few biome-time-step combinations (Figure. 4). The magnitude of change generally increased with time and with higher emission scenarios (Figure. S13-16). These predictions are consistent with other studies, suggesting a substantial intensification of fire regimes with greater warming. For example, panarctic wildfire emissions have been predicted to increase by 250% by 2100 under RCP8.5 (Abbott et al., 2016). Similarly, fire emissions in Finnish Boreal forests have been predicted to experience a 190% increase, even under RCP4.5 (McCarty et al., 2021). In Europe, burned area is predicted to increase between 180% to 360% until the end of the century under RCP8.5, but less than 60% under RCP2.6 (Wu et al., 2015). In another study in southern Europe, burned area is projected to increase 5–50% per decade under high emission scenarios (Dupuy et al., 2020). Another study projected a 40–100% increase in burned area with a 1.5° to 3°C warming for Mediterranean Europe (Bowman et al., 2020). An increase in burned area is likewise predicted for the Amazon and western USA (Abatzoglou et al., 2021; Bowman et al., 2020). In the grasslands of central Asia, the potential burned area is expected to increase 13% by 2080 (Zong et al., 2020).

**Figure 4.**
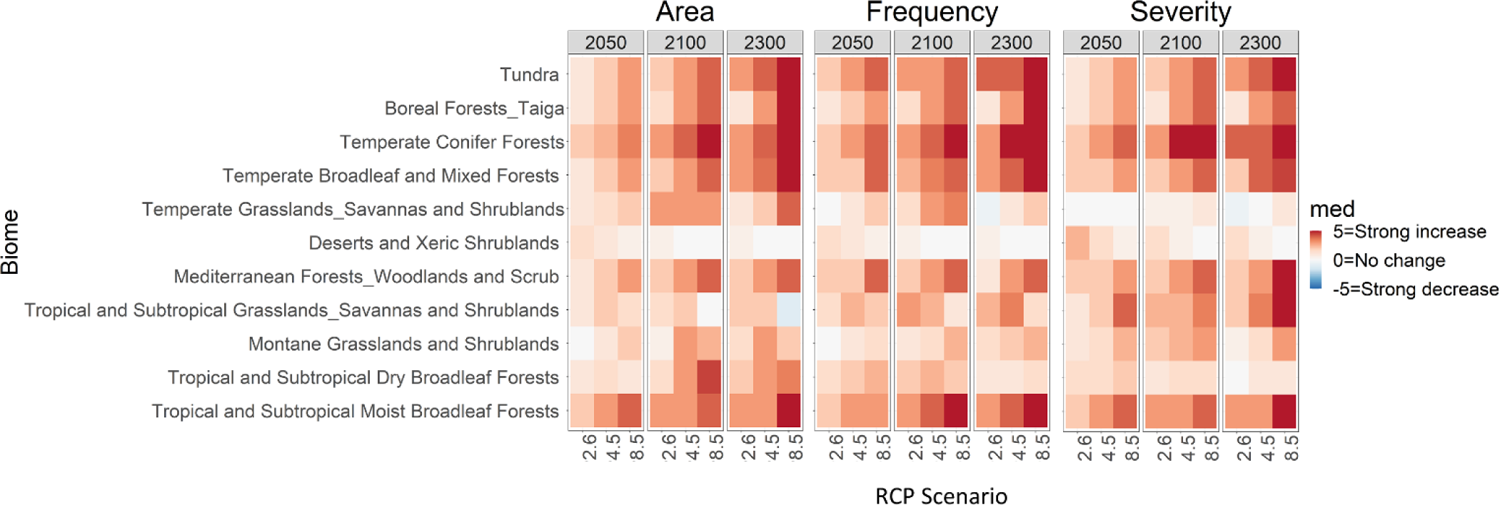
Direction and magnitude of change of fire regime characteristics as estimated by respondents. Median estimates of changes in fire extent (area), frequency, and severity for global warming scenarios RCP2.6, RCP4.5 and RCP8.5 in 2050, 2100, and 2300.

Contrary to most regions, less burning was predicted by experts in some parts of Africa under warmer scenarios, consistent with observations (Andela & Van Der Werf, 2014) that reveal a more intense fire regime under cooler and wetter climates that favor fuel build up in these dry regions (Daniau et al., 2013). More generally, fire frequency and severity are expected to decrease in fuel-limited ecosystems under drier conditions, while they should increase in ecosystems with ample humidity (Rogers et al., 2020).

When interpreting these survey results, it is important to recognize that within a single fire region, different ecological communities may experience divergent future fire trajectories (Moritz et al., 2012). For example, the risk of fire-climate interactions can vary in different type of conifer forests of western North America (moist-dry-subalpine) based on their elevation (Halofsky et al., 2020). This community-dependence was mentioned by respondents for most fire regions in this study. For example, substantial differences exist between eastern and western *Boreal forests* of the *Nearctic*, with the latter experiencing increasing trends in burn rates(Chavardès et al., 2022a). Although with projected climate change, convergence towards increasing burn rates is possible for eastern and western *Boreal forests of the Nearactic* during the mid-to late-21^st^ C.(Chavardès et al., 2022b).

### Consequences of fire regime change

Biodiversity, carbon stocks, and ecosystem services were anticipated to decrease with future fire regime change. Respondents estimated that the magnitude of change will increase for more extreme warming scenarios and longer timeframes in most biomes (Figure. 5, S17-20). However, the respondents suggested a general increase in albedo in the time frame 2050-2100, which could exert a transitory stabilizing effect on climate, and a further change in direction for some biomes through 2300 (Figure. 5, S21). Analysis for all scales, scenarios, and years are presented in the Supplementary Inforamtion.

**Figure 5.**
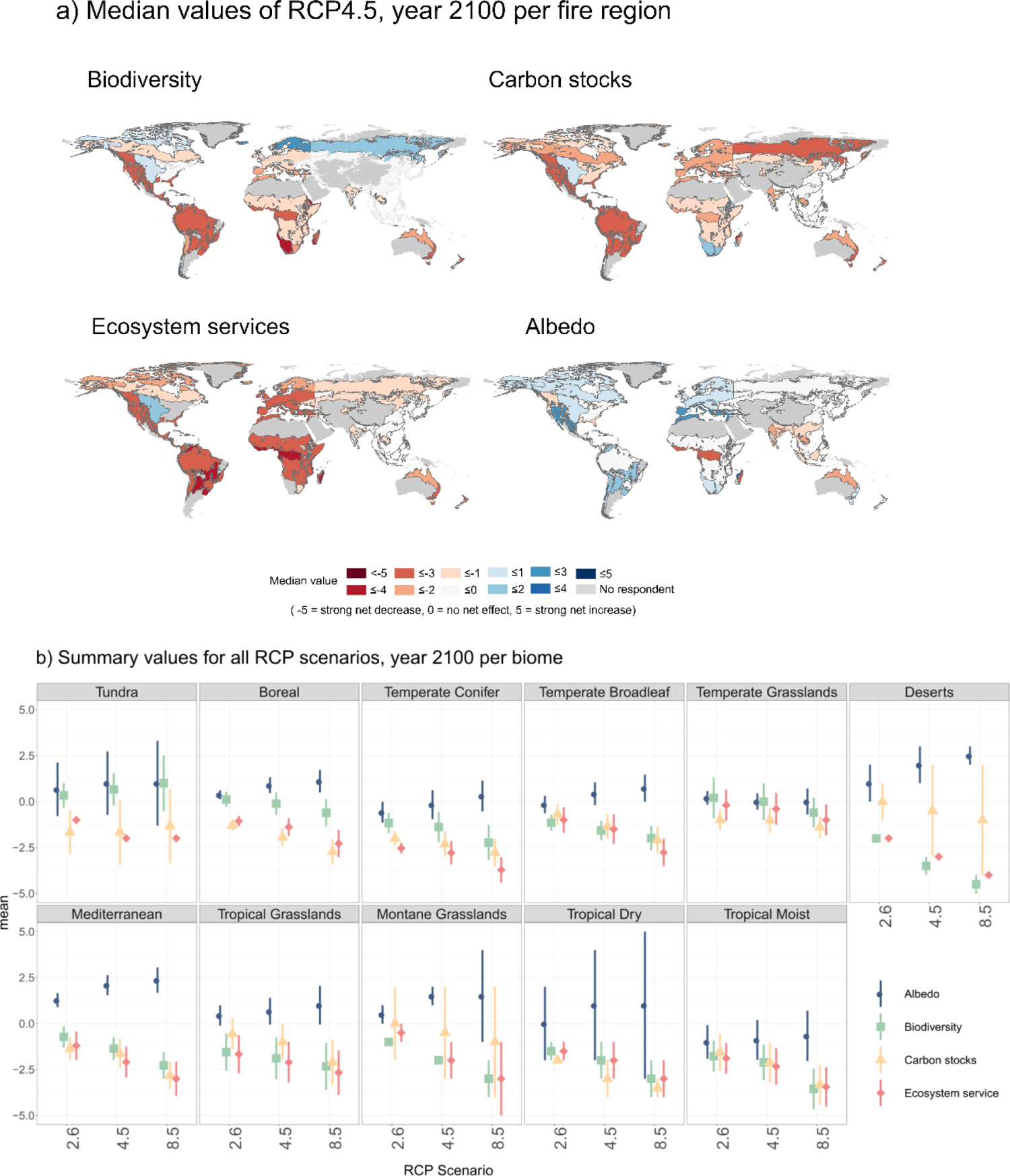
The net effect of predicted fire regime changes on ecosystem values in the future as estimated by respondents. a) The maps show the median value of experts estimate under RCP4.5, year 2100 (see Figures S18-21 for changes in RCP2.6 and RCP8.5). b) Average values and standard error for year 2100 under three RCP scenarios. The full names of the biomes can be seen in Figure1-b. Experts responded on a −5 to 5 scale for how strongly the future fire regime of the three RCP scenarios would affect the indicated parameters in the year 2100 (−5 = strong net decrease, 0 = no net effect, 5= strong net increase).

While it is difficult to assess agreement between respondents and the broader wildfire literature for such diverse fire regions, the overall results were generally aligned with the literature. It is anticipated that progressive increases in fire activity will impact biodiversity and ecosystem services in most regions, notably because ecosystem response to change and disturbance takes centuries to millenniums to reach equilibrium (Carcaillet et al., 2010, 2020). Local ecosystem services can be substantially altered by novel fire regimes. For example, in the Indian tropical dry forests, an increase in fire activity may negatively alter forest potential for water regulation by changing soil characteristics (Schmerbeck & Fiener, 2015), and atmospheric moisture recycling (Abbott et al., 2019). Similarly, changes in fire regime may also affect forest ecosystems and species that have historically been less affected by fire, as for beech forests (*Fagus sylvatica L.*) of Central Europe (Maringer et al., 2012).

In tropical biomes, an increase in extreme wildfires where fires are rare could affect tree mortality leading to habitat loss (Deb et al., 2018; Silveira et al., 2016). In the Mediterranean, an increase in fire events can create favorable conditions for some understory vegetation by temporarily reducing tree cover (Connor et al., 2019; Fournier et al., 2020). In savannas, longer fire intervals and elevated CO2 may create favorable conditions for woody vegetation (Sage, 2020), which can have a negative impact on biodiversity (Veldman et al., 2015). New fire regimes can also be favorable to some introduced and invasive taxa, including cheatgrass in some desert environments of the USA, tussock grass and pampas grass in Spain, and *Robinia, Ailanthus* in Portugal (Maringer et al., 2012). These species exploit novel disturbance niches to outcompete native vegetation during post-fire recovery. Consequently, both direct and indirect effects of fire regime change can alter plant community structure and composition with amplifying feedbacks on different aspects of fire regimes including frequency and extent (Bishop et al., 2020; Wan et al., 2014).

The consequences of fire regime change for ecosystem carbon balance are diverse. Novel climatic conditions in peatlands can slow their recovery from disturbances, decreasing carbon stocks (Loisel et al., 2021). More severe and frequent fires can threaten legacy carbon in Boreal forests (Walker et al., 2019), though changes to successional trajectories may offset or negate these losses in some cases (Mack et al., 2021). Respondents in this study projected a net decrease in *Boreal* carbon stocks under warmer scenarios. Another indirect effect of wildfire on carbon balance is local air pollution. For example, ozone produced during combustion can damage plant tissues, potentially doubling carbon losses by reducing photosynthesis post fire (Lasslop et al., 2019). Because human land use and fire regime are so closely linked, human actions such as deforestation coupled with cropland development can decrease carbon stocks at the same time as they modify the fire regime (Bowman et al., 2011; Cochrane & Bowman, 2021).

In other studies, it has been suggested that increased vegetation cover in higher latitudes will lead to decreased of albedo, which can have a more pronounced warming effect than greenhouse gases (Field et al., 2007). Not only can fires directly change the albedo of the region by altering land characteristics, but they also affect albedo by the pollutants they produce. For example, enhanced black carbon and soot deposition associated with increased fire disturbance will contribute to accelerated ice melting and decreased albedo (Aubry-Wake et al., 2022; McCarty et al., 2021). Conversely, increased tropical peatland fire can increase albedo (Ohkubo et al., 2021). Albedo may be reduced in the immediate aftermath of fire in sub-Saharan Africa, but it returns to pre-fire conditions within a few years (Gatebe et al., 2014). Perhaps because of the spatiotemporal complexity of the wildfire-albedo interaction, most respondents predicted little change in albedo.

### Our capacity to prevent, control, or adapt to future fire regimes

Respondents identified different fire regime drivers depending on the warming scenario and fire region. Under higher emissions (i.e., RCP4.5 and RCP8.5), most experts suggested that climatic factors would be the dominant driver of fire regime change. Conversely, under RCP2.6, only about half of the responses indicated that climatic factors would be the most important driver. Within *Australasia* and *Nearctic* fire regimes, climatic factors were identified as the most important driver for all scenarios. On the other hand, in the *Neotropic, Afrotropic*, and *Indo-Malayan* biogeographic realms, human activities were identified as most important. While vegetation and fuel were also frequently mentioned, these factors were never suggested as the most important driver of future fire regime changes (Figure. S22), highlighting the complex interplay amongst climate, fuel, and fire, especially on centennial timescales.

Survey results suggest that human actions for the next 20-50 years will be highly influential in determining how different ecosystem values (e.g., biodiversity, carbon stocks) are likely to change. Only 14% of the respondents indicated human actions have no effect, primarily in the case of albedo (Table. S1). Only 10% of responses recommended non-intervention, which instead was rated as a negative or unhelpful approach, though there were mixed opinions across and within fire regions.

About half of the respondents considered direct land management as an important approach for mitigating impacts of changing fire regime. In particular, fuel treatment, vegetation management, urban/suburban landscaping, and agriculture were identified as potentially useful mitigation approaches. There was high agreement that prescribed burning would help biodiversity and ecosystem services, but there were mixed opinions about its effect on carbon stocks, potentially because the area subject to prescribed burning is relatively small compared to the total burned area each year. There was less agreement about other fuel management techniques such as forest clearing or thinning, potentially because of the variety of vegetation types under consideration, and the lack of consensus in the literature on mechanical treatments. Even though activities such as clear-cutting reduce fuel, fire activity may increase due to the effects on microclimate and residual biomass, therefore changing fuel structure and composition (Bergeron et al., 2010; Cyr et al., 2009; Lindenmayer et al., 2009, 2020; Maxwell et al., 2019; Stephens et al., 2020). Conversely, traditional or Indigenous practices, such as cultural burning, were suggested as beneficial in reinforcing fire regime resilience and reducing damage by preventing extreme wildfire events (Christianson, 2015; Fernandes, 2020; Fletcher et al., 2021). There was a high level of agreement among experts who mentioned restoring vegetation (i.e., native habitat conservation and restoration) as a positive impact on all ecosystem values. There were mixed opinions about introducing fire resilient plants, but agreement on the positive effect of reducing flammable invasive plants. The natural or artificial selection of nonflammable species was mostly considered to have a negative effect on biodiversity and ecosystem services but variable effects on carbon and albedo.

Several landscape management strategies had general support, including increasing landscape heterogeneity, diversification, and reduction of landscape flammability by targeted land use, as well as creating buffer zones around primary and old-growth forests (Barredo et al., 2021). Attention to the human-wildfire interface was a common recommendation, as certain levels of population (housing) density, wildland-urban interface, and landscape connectivity can dramatically affect the characteristics and societal consequences of fire (Archibald et al., 2012; Kelley et al., 2019; Moritz et al., 2014; Syphard et al., 2007).

Direct fire management was recommended in 17% of expert responses. In the case of fire suppression as a direct fire management strategy, there was less agreement about the direction of effects on different factors. In some studies, it has been shown that fire suppression had a negative impact on fire-dependent ecosystems. Such policies have led to fuel accumulation and increased flammability that has contributed to today’s extreme wildfire events (Marlon et al., 2012; Parisien et al., 2020; Schoennagel et al., 2017; Valese et al., 2014). A greater proportion of respondents (23%) indicated the importance of social or political awareness and action. These included climate mitigation, education of the public regarding fire danger and ignitions, direct or indirect conservation policies, and incorporation of Indigenous or traditional knowledge (Table. S1). Although we have summarized and combined all the management suggestions from experts of different fire regions, we remind the reader that suitable management applications can significantly vary between and within biomes based on various factors (detailed responses for each fire region can be found in the Supplementary Information).

There was strong agreement among experts, except for the *Afrotropic* and the *Indo-Malayan* biogeographic realms, that under RCP8.5, humans will have a decreasing capacity to control wildfires. This was most obvious for the *Neotropic* and *Nearctic*, and *Australasia* for the later years (2100 and 2300). Experts on *Neotropic* and *Nearctic* fire regimes indicated that this same decreased capacity was likely to apply under RCP4.5. 70% of the respondents indicate that under RCP2.6, humans will maintain some ability of managing fire-impacts (Figure. S23).

### Charting a course in a world of uncertainty

For each question, respondents identified the main sources of uncertainty (Table 1), which included limited observational data, inadequate modeling frameworks, and system complexity, particularly social dimensions. Respondents emphasized that the impact of different human activities is not completely understood for the past or present and that untangling different fire drivers can be difficult due to multiple interactions and feedbacks, which are often not represented in coupled models. Respondents also mentioned that the unprecedented rate of climate change and diversity of human activity made estimations of fire return intervals and other dimensions of fire regime uncertain. Respondents were uncertain about emergent economic and policy direction, beliefs, and technologies as tools to combat changing fire regime. Additional sources of uncertainty included the exclusion of small fires below the detection of satellite census and a lack of information about fire severity impacting ecological succession, albedo, and carbon-climate feedbacks.

**Table 1.** Sources of uncertainty identified by respondents

| **Past** |  | **Current** |  | **Future** |  | **Management** |  |
| --- | --- | --- | --- | --- | --- | --- | --- |
| Sources of uncertainty | % | Sources of uncertainty | % | Sources of uncertainty | % | Sources of uncertainty | % |
| Limited proxy-paleo data | 23 | Spatial variability | 31 | Vegetation shift | 12 | Political and socio-economic | 22 |
| Spatial variability | 17 | Limited data | 17 | Ecosystem interactions and feedbacks<sup>2</sup> | 12 | Capacity and effectiveness of management and intervention | 13 |
| Proxy resolution | 15 | Combustion/severity | 8 | Climate change | 10 | Climate change | 11 |
| Model limitation | 11 | Small fire detection/remote sensing | 7 | Political and socio-economic | 8 | Ecosystem interactions and feedbacks | 10 |
| Human influence<sup>1</sup> | 11 | Human activity | 8 | Model limitation | 7 | Technology developments | 6 |
| Proxy reliability | 8 | Recent changes | 4 | Fire management and intervention | 7 | Science and management connection | 4 |
| Temporal variability | 6 |  |  | Land use change | 6 | Vegetation state | 4 |
| Chronological | 5 |  |  | Precipitation | 6 |  |  |

<sup>1</sup>E.g., Fire management history, land use history, etc.
<sup>2</sup>E.g., Climate-vegetation dynamics, albedo and new vegetation, vegetation-fire interaction, fuel-ignition relationships, climate-human intervention, climate-vegetation feedbacks.

While our study may have limited application in many specific management contexts, there were some general patterns that could be informative for policymakers, managers, and researchers. First, there is a high level of agreement across regions that the risk of damaging fire characteristics will be greater under higher emissions. While the specific consequences will vary by fire region and habitat type, the overall message is clear: rapid reduction of greenhouse gas emissions is needed to restore Holocene-like climate conditions. Otherwise, the emergence of novel climates and fire regimes outside of the range of Holocene variability, will complicate our ability to conserve habitats, ensure healthy communities, and preserve terrestrial carbon uptake and storage. Without a reduction in emissions, changes in fire regime could eclipse climate mitigation policies such as negative emissions through reforestation and afforestation (Anderegg et al., 2020; Veldman et al., 2019). Any carbon uptake from recovered or cultivated forests could be negated by the increased fire frequency or intensity projected for many regions (Hammond et al., 2022; Smith et al., 2020).

A second overall lesson is that knowledge of past fire regimes provides perspective on how climate, vegetation, and human actions interacted to shape fire in the Earth system (Marlon et al., 2008; Molinari et al., 2018; Pechony & Shindell, 2010). For example, paleo-ecological knowledge about vegetation community and historical amplitude of fire regime change in a given biome can provide estimates of historical thresholds and optimal vegetation structure for management purposes (Hennebelle et al., 2018). Likewise, fire histories show human-vegetation-climate linkages, such as decreasing tree cover creating microclimates favorable to the encroachment of flammable vegetation in the understory (Feurdean et al., 2020). An overarching lesson learned from looking at Holocene fire histories is that projected conditions are generally unprecedented, meaning that human-fire interactions could have extreme and unexpected consequences (Bova et al., 2021). We should not assume that historical management practices will suffice (Crandall et al., 2021; E. C. Ellis et al., 2021; Pyne, 2007) given accelerated rates of vegetation change since c. 4000 BP (Mottl et al., 2021), the emergence of novel biotic and abiotic conditions (Burke et al., 2019; Finsinger et al., 2017; Ordonez et al., 2016), and increasing populations. For example, the expansion of human development in fire prone areas in the western US is increasing both wildfire incidence and cost of suppression (Balch et al., 2017a, 2017b).

Third, despite the substantial uncertainties associated with fire regimes, mitigation efforts such as allowing some fires to burn to reduce fuel loads, prescribed burning, and fuel treatments will help limit fire impacts and cost (Harris et al., 2016; Mietkiewicz et al., 2020; Moritz et al., 2014; Radeloff et al., 2018). Likewise, the conservation of large, contiguous ecosystems allows the use of more effective wildfire management tools such as prescribed burning and increases resilience when unexpected wildfire behavior emerges (Bentley & Penman, 2017; Driscoll et al., 2016; Miller, 2020).

While this study brought together a diverse group of paleo and present fire researchers, we point out that our group is not geographically balanced. Despite invitations to several hundred researchers, we only received a few responses for some fire regions (Figure. 1b), including the African subtropical and tropical grassland region, which accounts for a large portion of global area burned (Ramo et al., 2021). This reflects the broader geographical and cultural bias in ecological research generally and wildfire research specifically (Bradstock et al., 2002; Hantson et al., 2016; Metcalfe et al., 2018), highlighting the need for more diverse research networks.

We also point out a potential bias in climate scenarios. Low emissions scenarios such as RCP2.6 or SSP1 are sometimes omitted from model inter-comparisons because they are often deemed unfeasible (McGuire et al., 2018). In reality, it is the more extreme warming scenarios that are becoming less likely (Hausfather & Peters, 2020; Lovins et al., 2019). We found substantial differences in fire regimes between RCP2.6 and higher emission scenarios. Given the current rate of the global transition to renewable energies (Breyer et al., 2022; Hausfather & Peters, 2020; O’Neill et al., 2020), it is important to understand both the “low” as well as the “high range” of climate change futures.

### Expert assessment application in fire management

Decision-making in landscape and fire management requires a great understanding of complex human and natural systems and their interactions with fire. Currently, policymakers and managers working on fire issues are operating in a complex environment with sometimes conflicting traditional, scientific, and political information and priorities. As the physical, biological, and human factors controlling wildfire behavior change rapidly in the Anthropocene, improving science-policy-management integration would be highly beneficial for human wellbeing and ecosystem function. The primary scientific knowledge that is used by decision-makers regarding the future is based on models or single expert advice. As several respondents mentioned in this study, quantitative models cannot capture all the factors influencing the evolution of fire behavior. Various types of expert elicitations can complement quantitative models to generate more robust and reliable guidance that allows adaptive management. This could range from informal interpretation of model outputs by expert panels to iterative combinations, such as expert input on supervised and unsupervised machine learning models. These approaches should be (and already are in some cases) used in various aspects of fire management, from detecting fires to planning and policy, by providing a benchmark or improving the initial parameters and weights (Jain et al., 2020).

At some level, we are not proposing a new role for experts in policymaking and management, but we are suggesting that the integration of local expert knowledge be done in a more rigorous and robust way. Policymakers and managers are making decisions based on available information that is often filtered through informal information networks, especially trusted relationships and professional networks. By involving more independent experts to inform policy, it is possible to limit the “one-expert syndrome”, where the opinions or a single knowledge holder dominate the decision-making process, whether that view represents the broader consensus.

Our study shows that experts have different views about different aspects of wildfires (i.e., there is not strong consensus on several dimensions of current and future fire behavior). These biases come from various sources, including scientific background and study methodology (Oppenheimer et al., 2019). Therefore, developing methods to transfer group expertise to decision-makers is critical. Even though our study has been done on a global scale to identify large patterns, this method can be beneficial for local management by bringing local expertise (that in many regions is not represented in scientific publications that are mostly produced in developed countries), such as indigenous knowledge, into the decision-making process (Christianson, 2015).

## Supporting information

SI_Figures_Questionnaire

SI_DataAnalysis

## Acknowledgements

This study emerged during the PAGES-supported Global Paleofire Working Group 2 workshop “Fire history baselines by biome” held in September 2016 at Château de la Tour, Beguey (Bordeaux, France) led by A.-L. D. and Tim Brücher. PAGES, Past Global Changes, is funded by the Swiss Academy of Sciences and the Chinese Academy of Sciences and supported in kind by the University of Bern, Switzerland. Financial support was provided by the U.S. National Science Foundation award numbers 1916565, EAR-2011439, and EAR-2012123. Additional support was provided by the Utah Department of Natural Resources Watershed Restoration Initiative. SSS was supported by Brigham Young University Graduate Studies. MS was supported by National Science Centre, Poland (grant no. 2018/31/B/ST10/02498). JCA was supported by the European Union’s Horizon 2020 research and innovation programme under the Marie Skłodowska-Curie grant agreement No 101026211. A.-L.D acknowledge PAGES, PICS CNRS 06484 project, CNRS-INSU, Région Nouvelle-Aquitaine, University of Bordeaux DRI and INQUA for workshop support. We dedicate this manuscript to our late colleague Dr. Daniele Colombaroli.

## Notes

### Competing Interest Statement

The authors have declared no competing interest.

### Summary of Updates

I added all the authors' individual names instead of the group name.

