## Supplementary material for "Assessing changes in global fire regimes": SI_Figures_Questionnaire

*Affiliations are placed at the end of the document

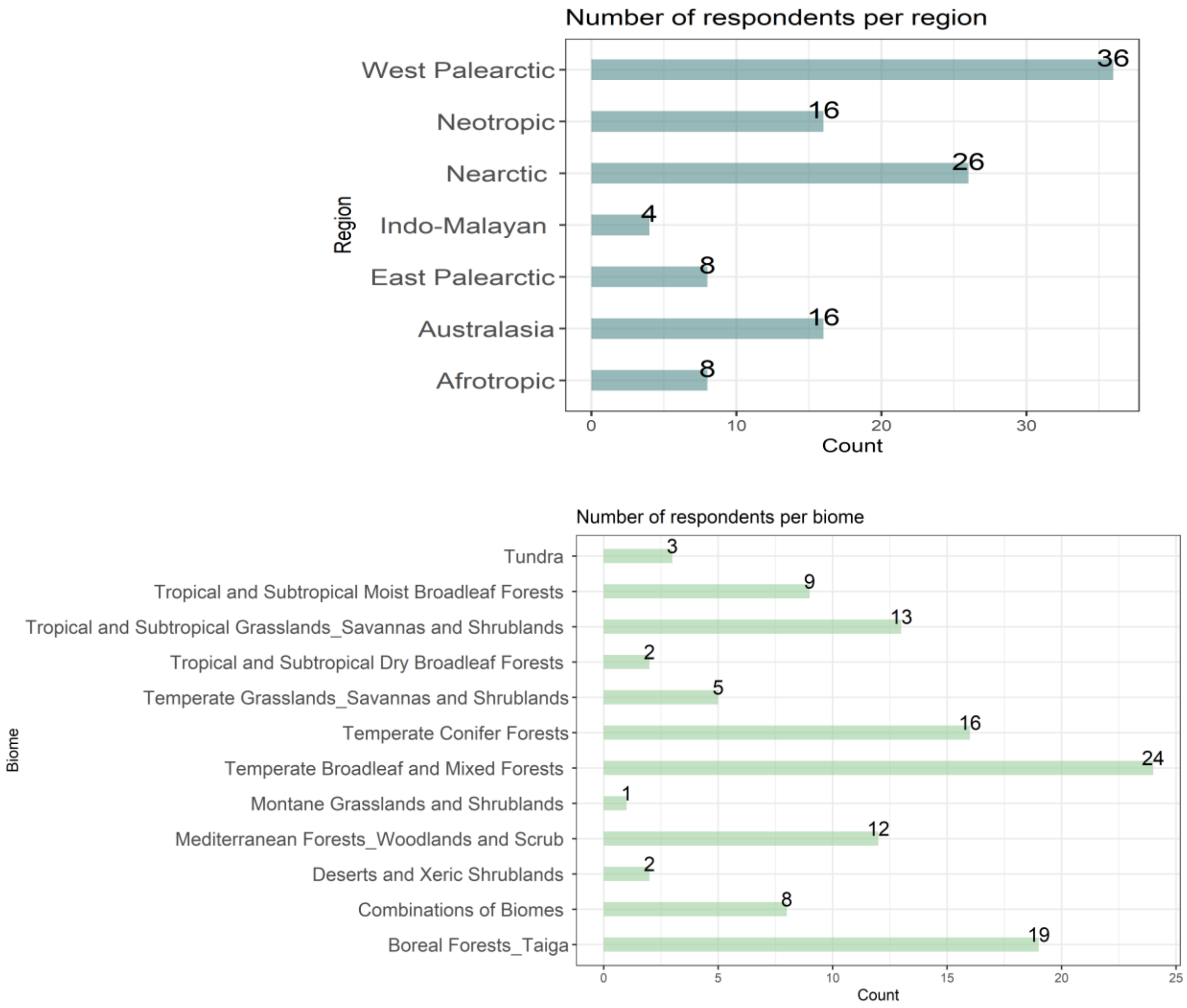

**Figure S1.** Number of respondents per region and biome

**Paleo:**

There has been ca. six intervals of rapid climate change during the Holocene, mainly due to natural reasons(1). However, not all regions across the globe have experienced climatic changes to the same extent(2). For example, the tropics did not have a temperature peak in the early-mid Holocene as seen in most regions(3). It should also be noted that maybe the seventh rapid climate change which is the most intense and fast is the transition of the Anthropocene (1950-present) with the emission of anthropogenic greenhouse gases. Vegetation change rate has also been different in different regions of the globe during the Holocene. Anthropogenic activities have been known as the primary reason of global vegetation change in the late Holocene. This is mainly because the rate of vegetation change in this time has exceeded the rate of vegetation change during the last deglaciation which the temperature rise was significantly higher (~6°C) than late Holocene (~ 1°C)(4). It has been challenging to detangle the role of humans versus climate for fire regime changes. Even though there is evidence that humans have impacted and shaped most of the terrestrial land from 12ka BP(5), there is low certainty about the human role in fire regimes until 6ka BP(6–8).

Number of fire regime state changes during the Holocene

a) Region b) Biome

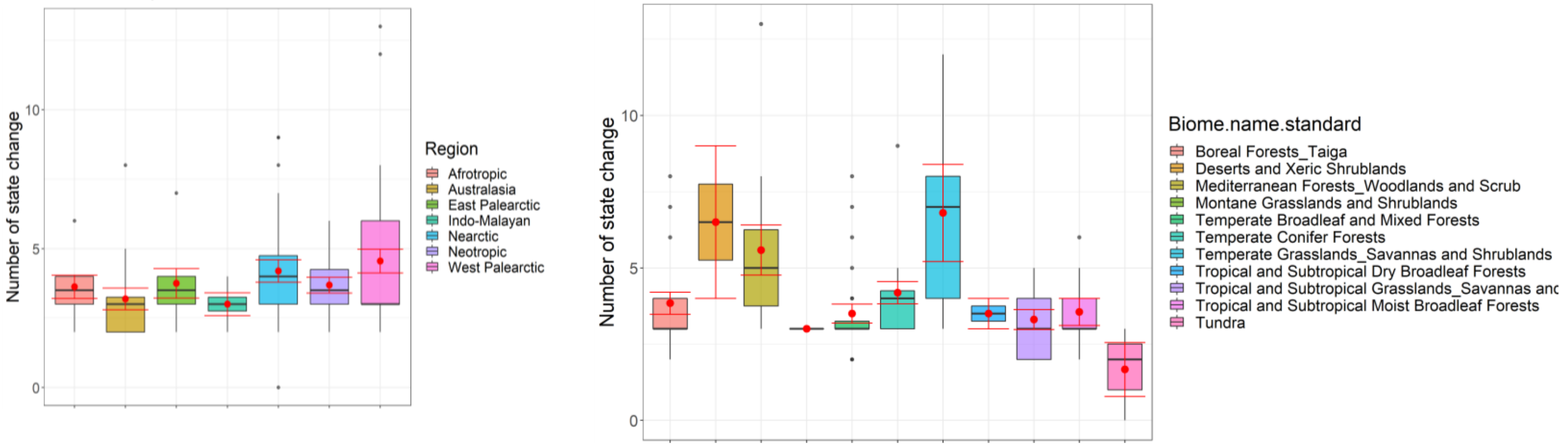

**Figure S2.** Number of fire regime state changes during the Holocene. The boxplots represent the median (black line) and average values (red dots). Red lines represent standard error.

Occurrence of the three largest fire regime state changes during the Holocene

a) Global

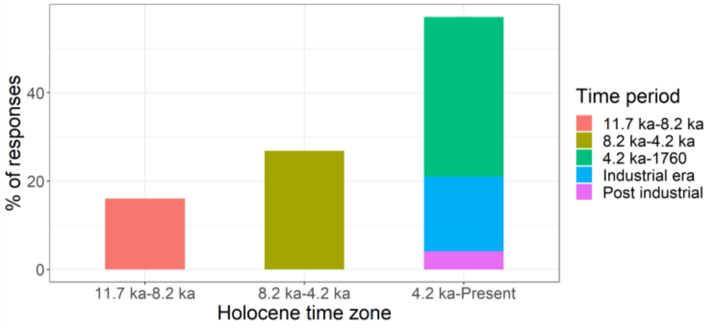

b) Region

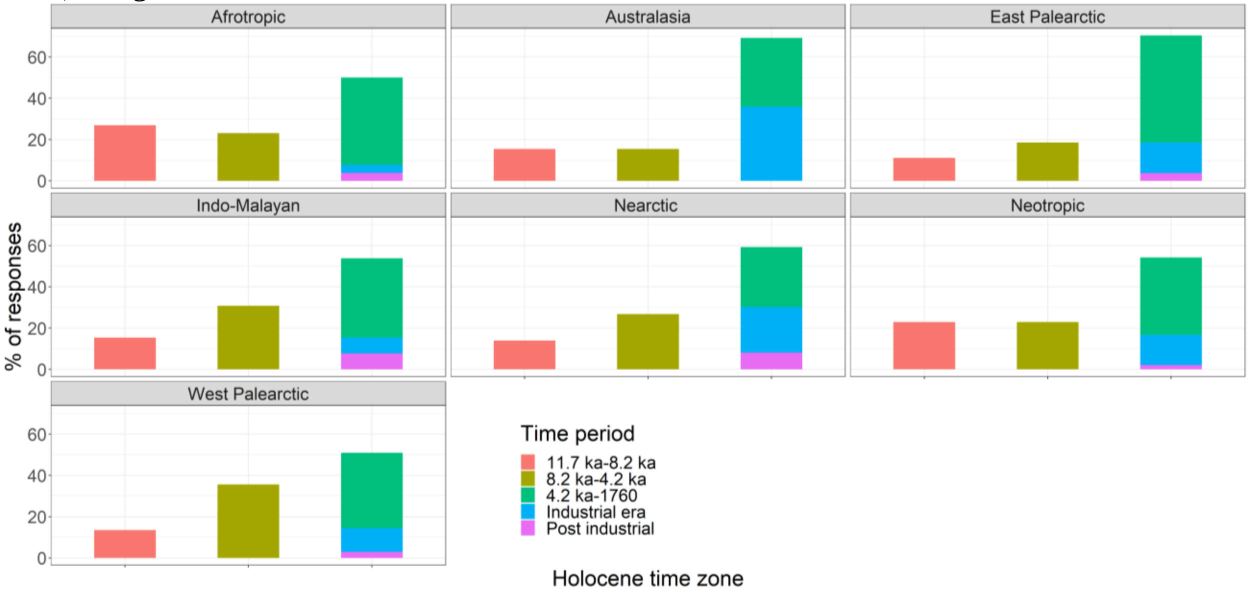

c) Biome

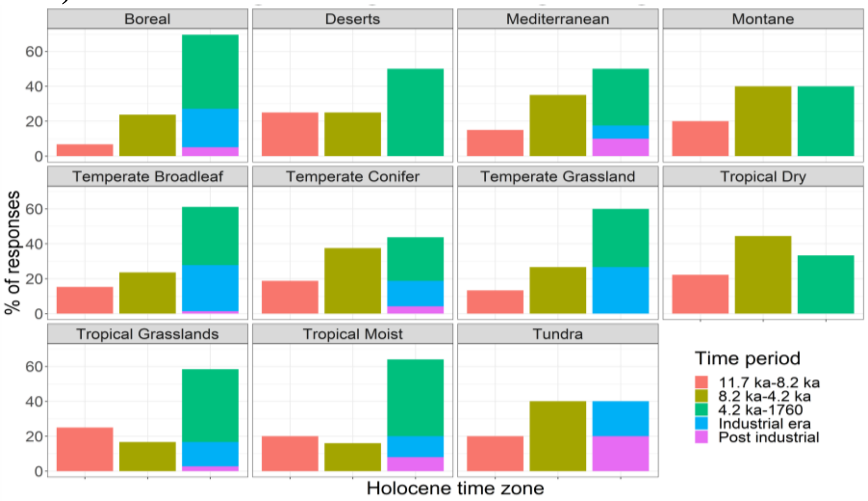

**Figure S3.** Time periods for the three largest fire regime state changes during the Holocene (the order of time periods for all plots are similar to the global plot). For the purpose of this study the following time periods were defined: 11700 8200BP: Early Holocene; 8200-4200BP: Mid Holocene, 4200 -0: Early Holocene; 262-70BP: Industrial era,70-0BP: Postindustrial.

Drivers of major fire regime state changes during the Holocene

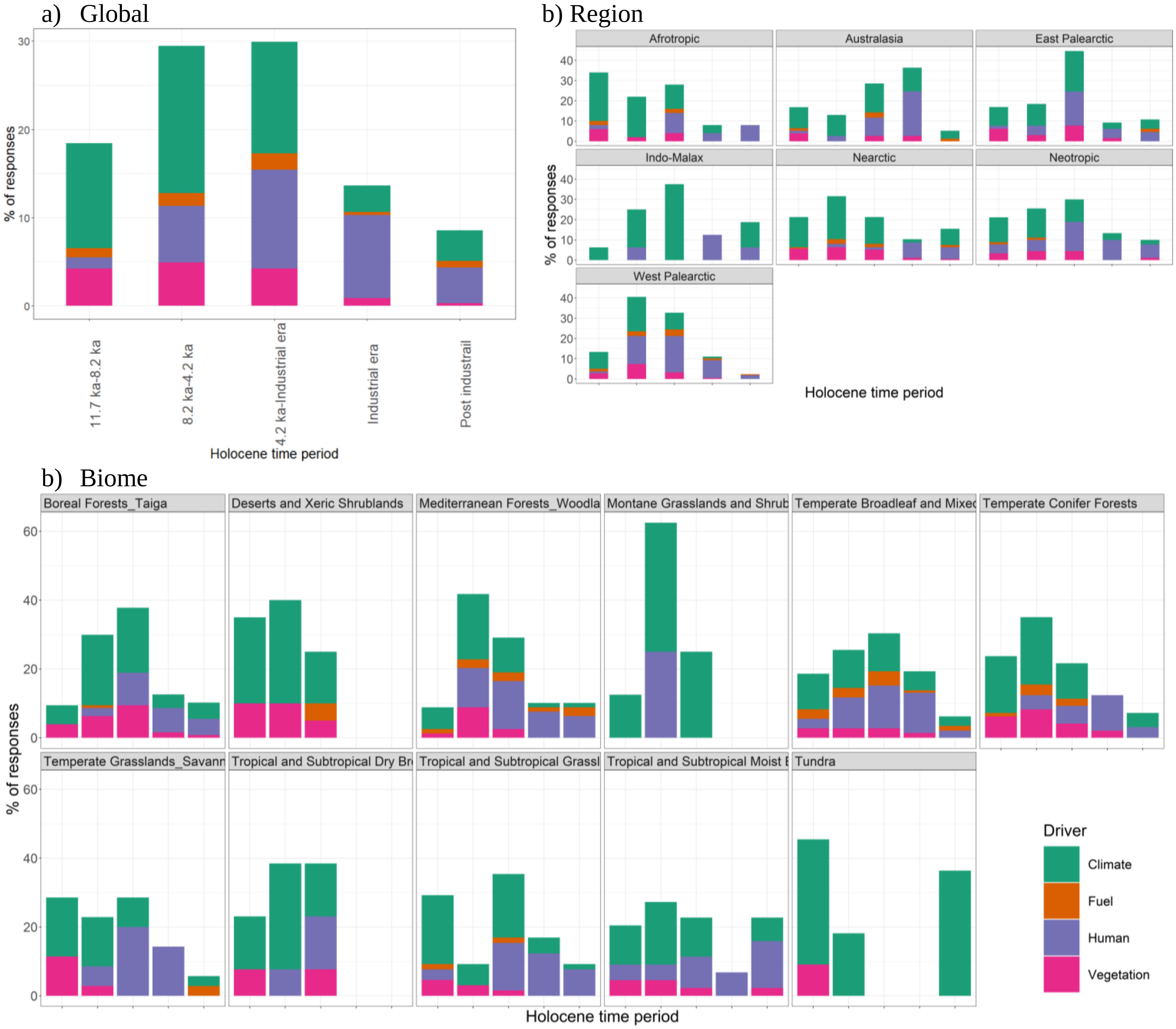

**Figure S4.** Drivers of the three largest fire regime state change during Holocene (the order of time periods for all plots are similar to the global plot).

a) Region b) Biome

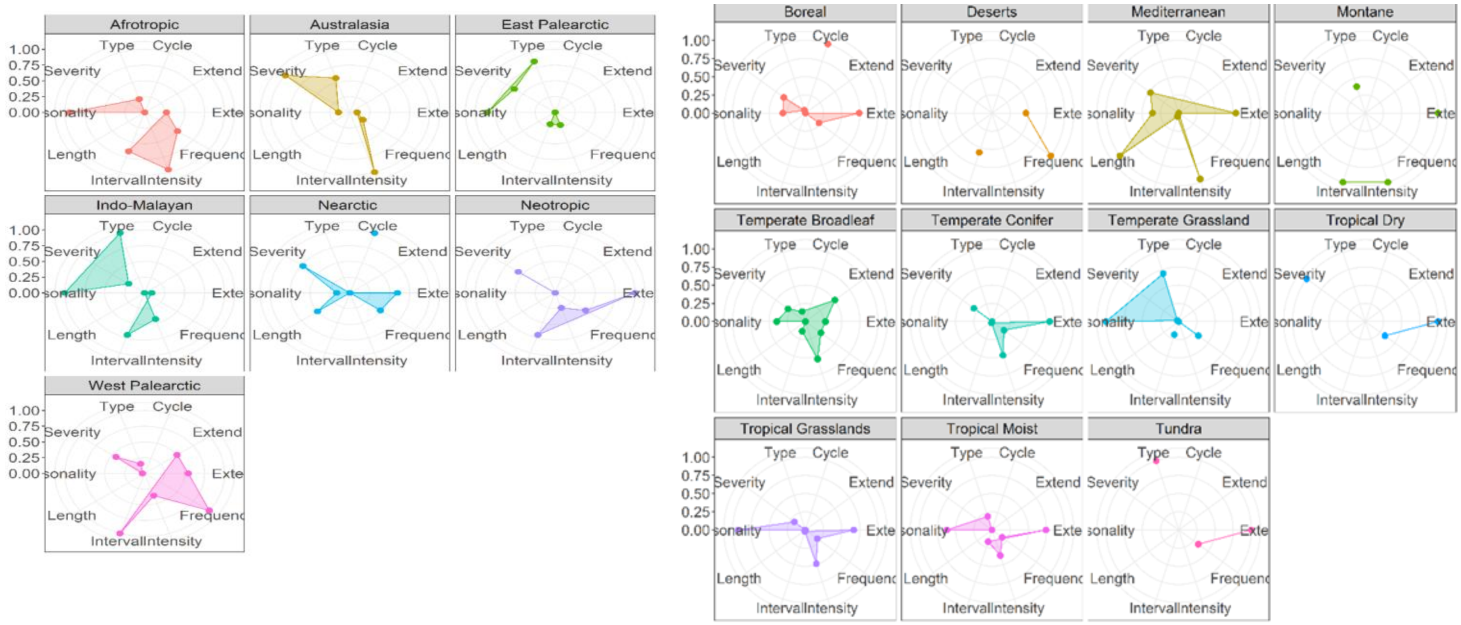

**Figure S5.** Fire regime aspects that post-industrial society influence most strongly. The y axis represents percentage of responses. Including: Spatial (Extent, severity, type and intensity) and Temporal (Frequency, seasonality, cycle, interval).

#### Current:

The median estimation of the duration of the current fire regimes was less than 200 years for 10 out of 11 biomes (low agreement for boreal forests, temperate conifer forests and tundra, based on standard error). For five biomes, this duration was less than 75 years. Responses for the tropical and subtropical dry broadleaf forest suggested indicated substantially more persistent fire regimes than other regions of >1000 years (low agreement; Fig.S6). Experts estimated the percentage of annual area burnt to be less than 5% globally for 7 out of 11 biomes. Tundra and boreal forests had the least estimated area burned <1%. Some biomes, such as tropical and subtropical grasslands and savannas and tropical and Subtropical Dry Broadleaf Forests, were estimated to have a significantly higher area burned by ~35% (Fig.S7). Experts estimated the current fire interval to be between 2 years (tropical and subtropical dry broadleaf forests) to 1900 yrs. (tundra). Overall, there was low agreement within experts of different biomes for the fire return interval (Fig.S8).

Persisted time of the current fire regime

a) Region b) Biome

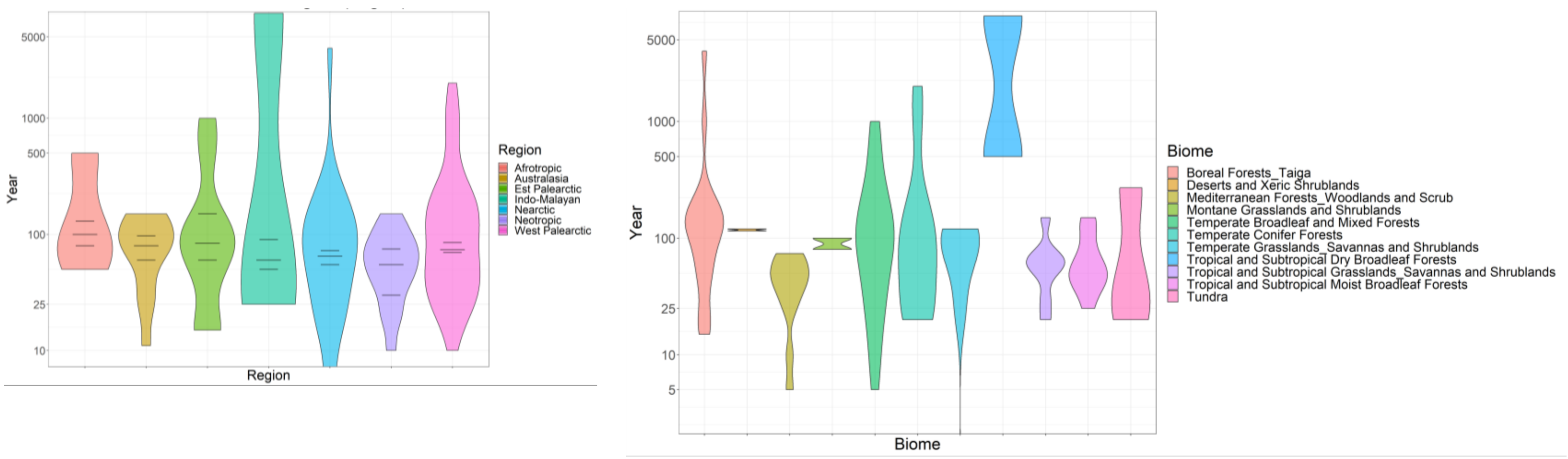

**Figure S6.** Violin plots showing persisted time of current fire regime. The violin plots show the among-expert distribution of the central estimates (width indicates number of estimates in that range).
The horizontal black lines indicate the among-expert medians of lower, central, and upper estimates.

Burned area

a) Region

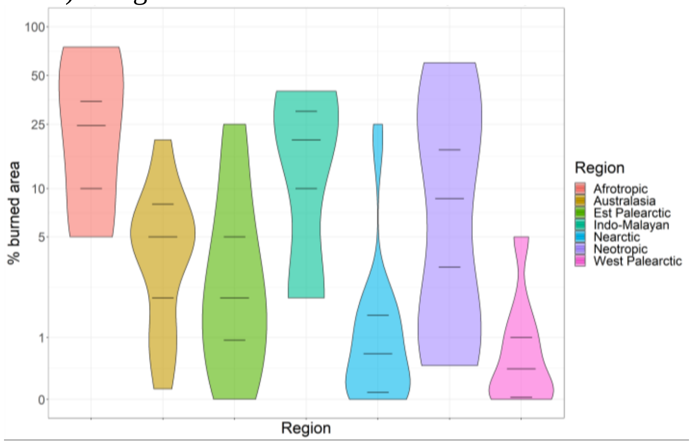

b) Biome

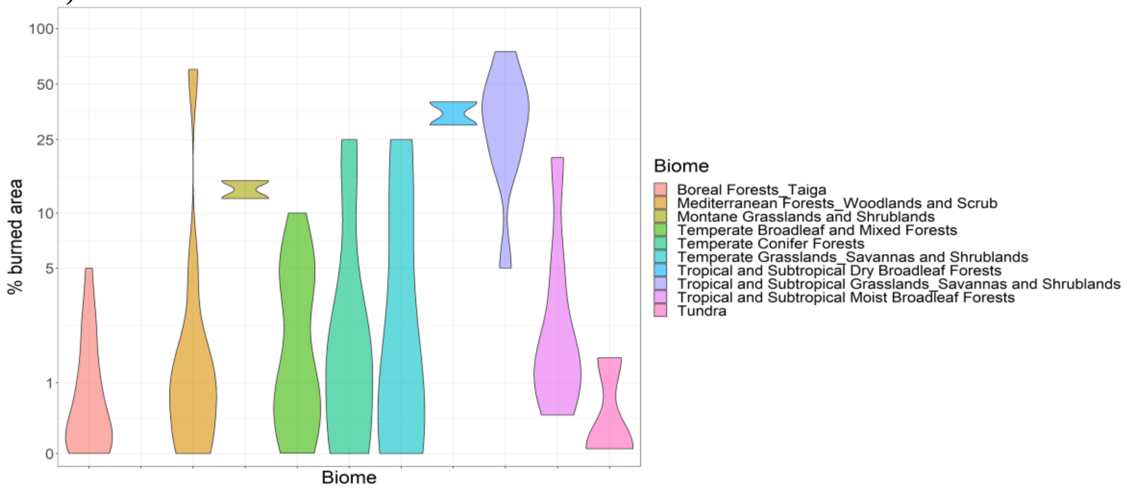

**Figure S7.** Violin plots showing burned area of current fire regimes.

Fire return Interval

a) Region

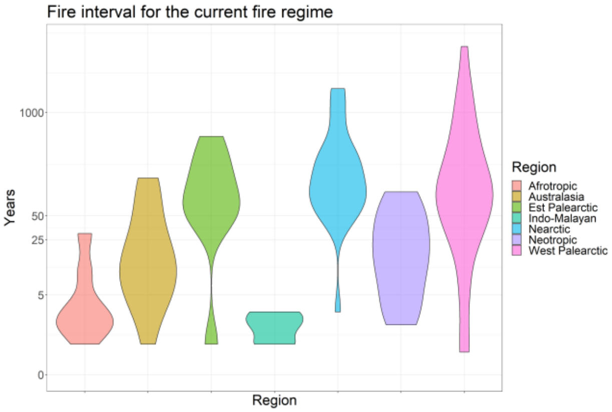

b) Biome

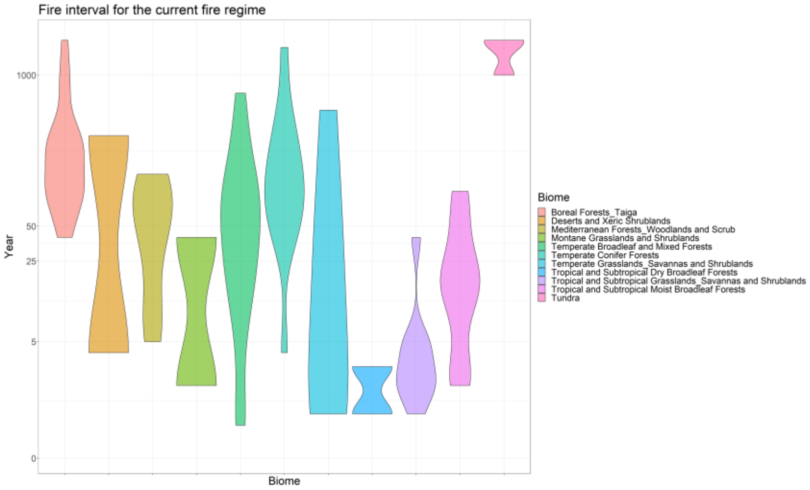

**Figure S8**. Violin plots showing fire return interval for current fire regimes.

Q9 does not have any figures.

#### Future

Likelihood of fire regime state change in the future

a) Global

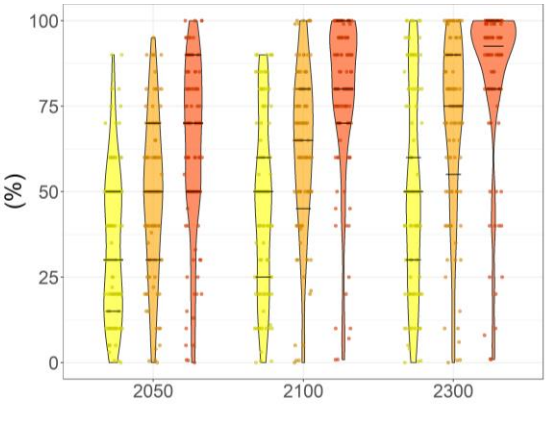

b) Region

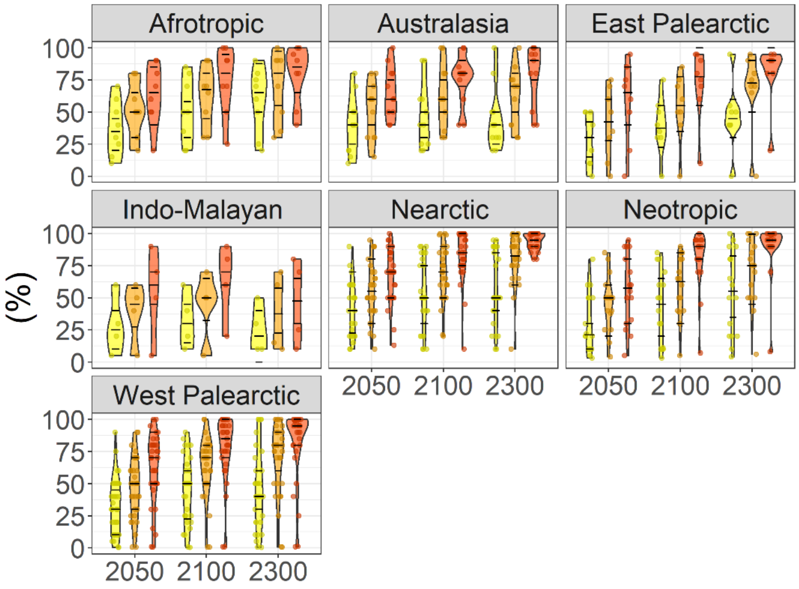

c) Biome

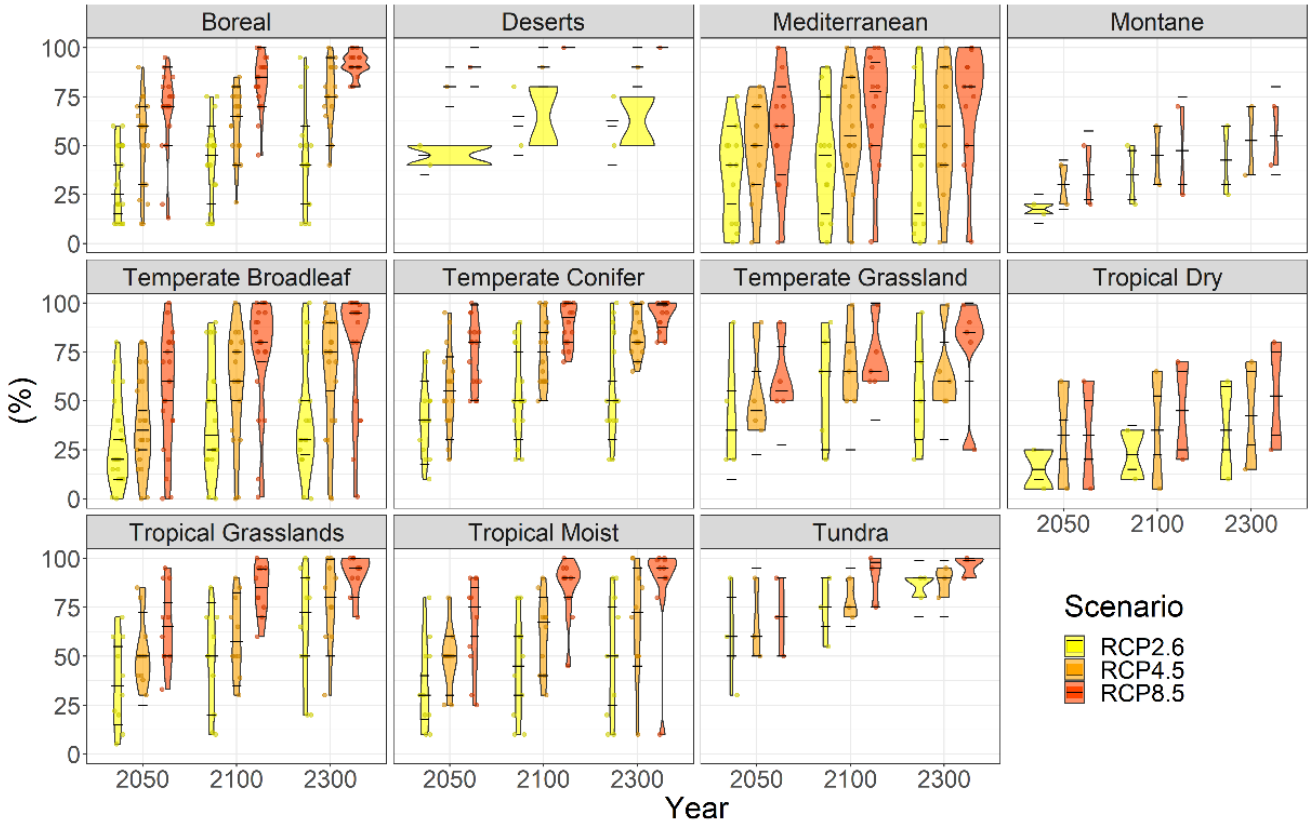

**Figure S9.** Estimated likelihood of fire regime change for years 2050, 2100 and 2300 under three RCP scenarios (RCP2.6, RCP4.5, and RCP8.5). The horizontal lines show the median "lower", "central" and "upper" estimates with a 90% confidence interval. The points and ranges show the estimates for each expert individually.

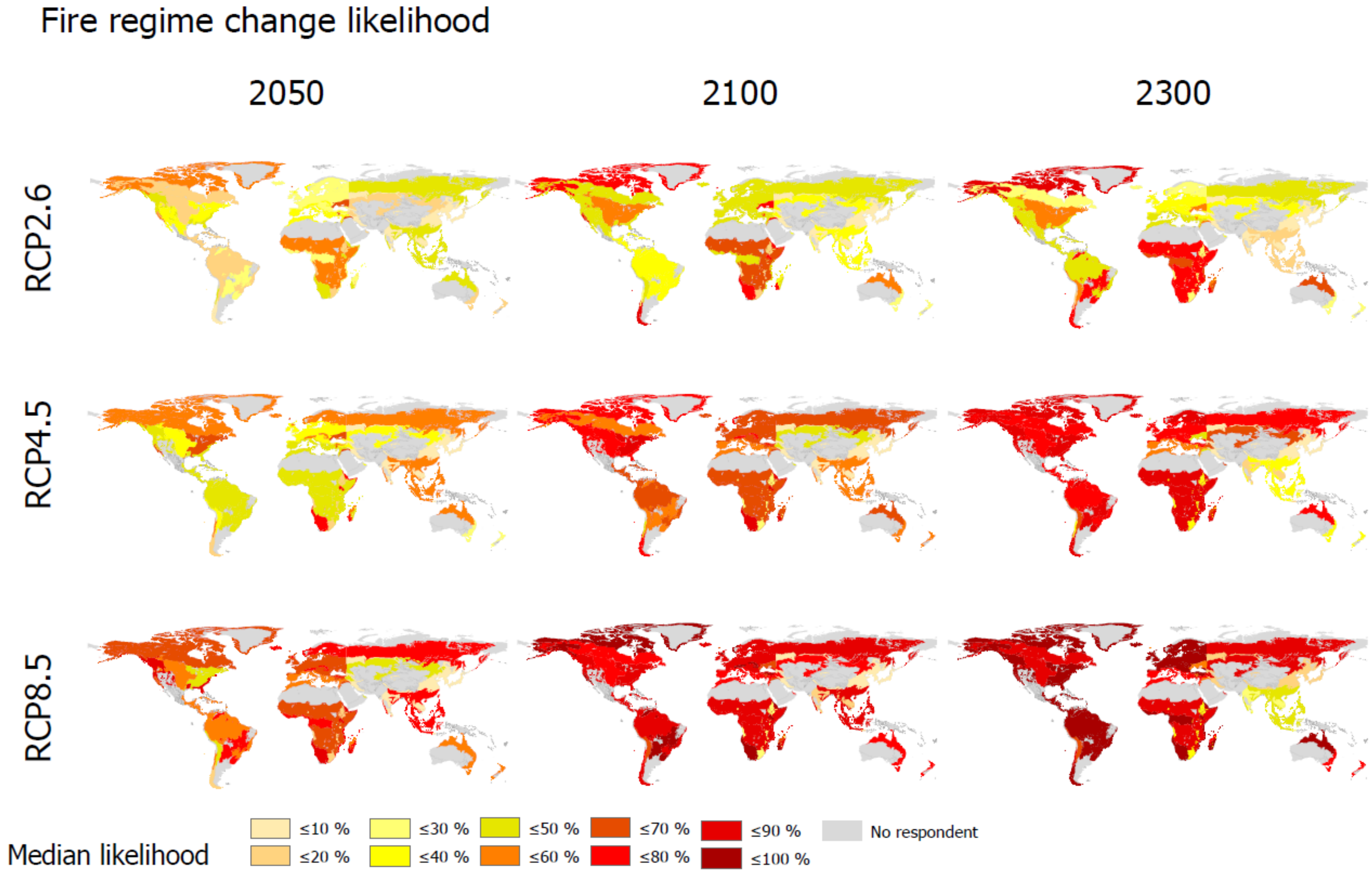

**Figure S10.** Median likelihood of fire regime changes for years 2050-2300 under three RCP scenarios

Magnitute of fire regime change likelihood between differnet RCPs

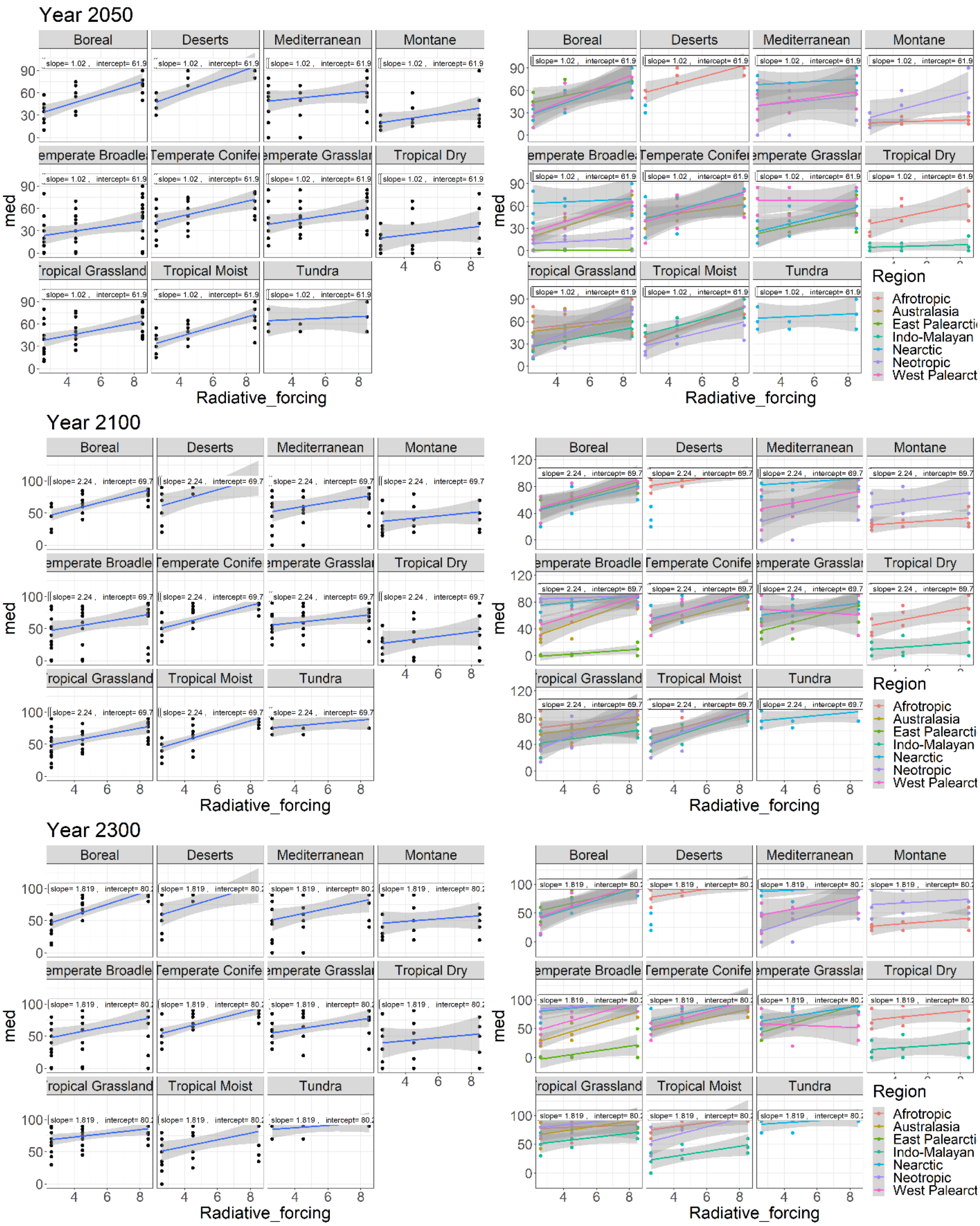

**Figure S11**. Magnitude of fire regime change calculated based on based on the slope between the median likelihood of a fire regime change (%) from all quantiles and the amount of radiative forcing for the three RCP scenarios (W m-2). The dots are individual’s estimates.

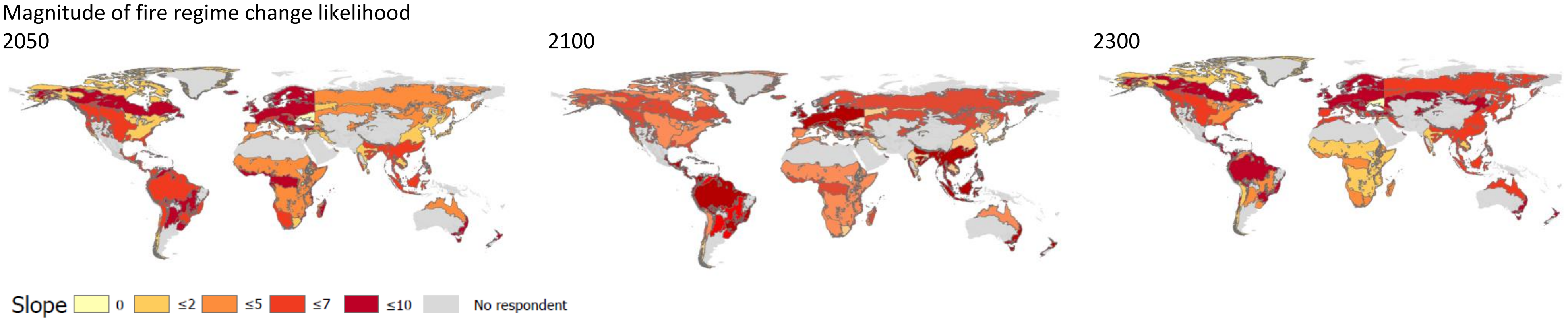

**Figure S12**. Magnitude of fire regime change likelihood between RCP2.6-4.5-8.5. The value of the slope is based on the median likelihood of a fire regime change (%) and the amount of radiative forcing

Experts of *Temperate broadleaf and mixed forests of Australasia* (high agreement), *Temperate conifer froests of Nearctic* (high agreement), and *Tropical and subtropical moist boreadleaf forests of Neotropic* (high agreement) predicted the most increase in burn area under all scenarios and timelines (Fig.S13). Unlike the *Deserts and xeric shrublands of Nearctic* for the *Deserts and xeric shrublands of Afrotropic* (one respondent), *Tropical and subtropical grasslands_savannas and shrublands of Afrotropic* (low agreement), and *Temperate broadleaf forest of East Paleoarctic* (one respondent) a decrease of burned area was predicted.

Direction of change of fire regime charecteristics

a) Global

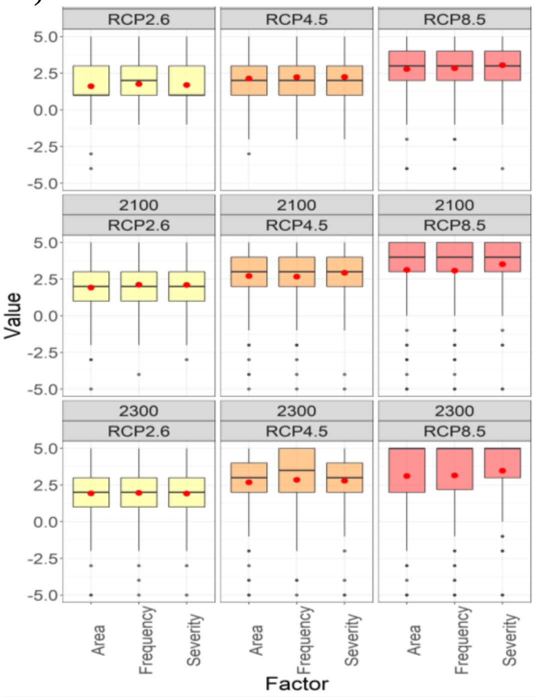

b) Region -2050

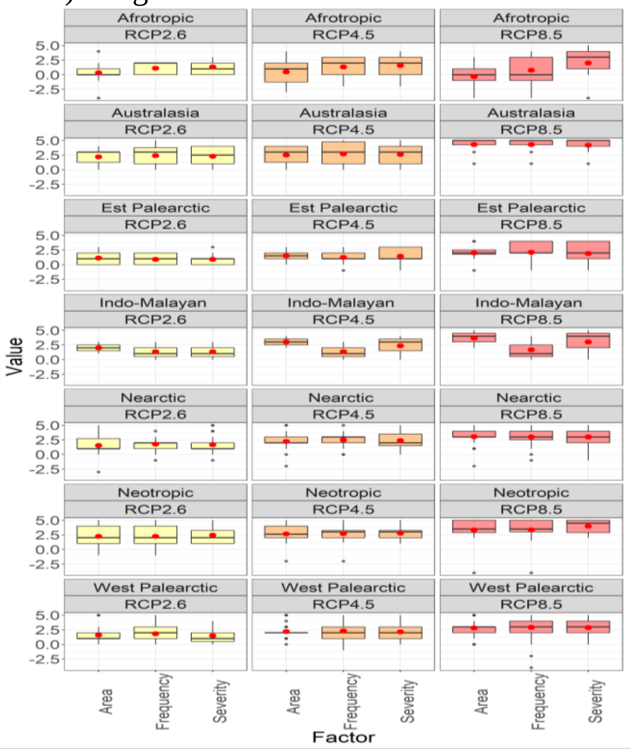

Region-2100

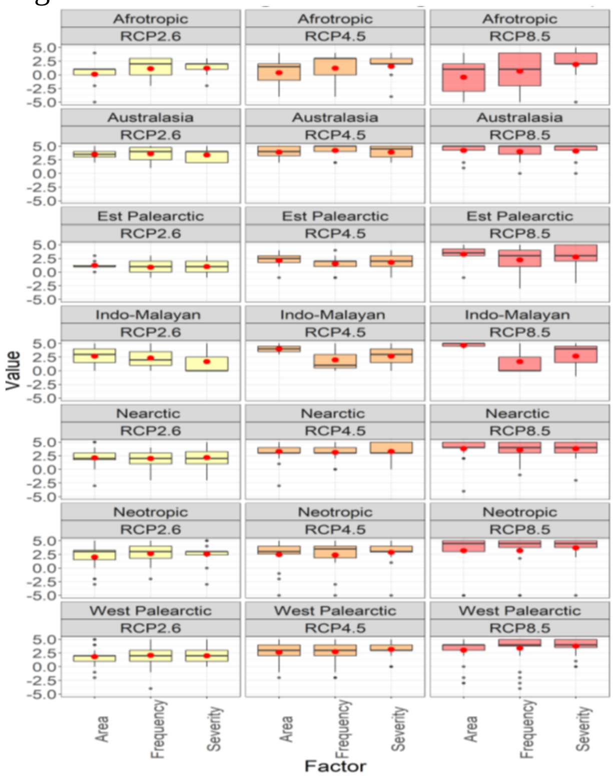

Region-2300

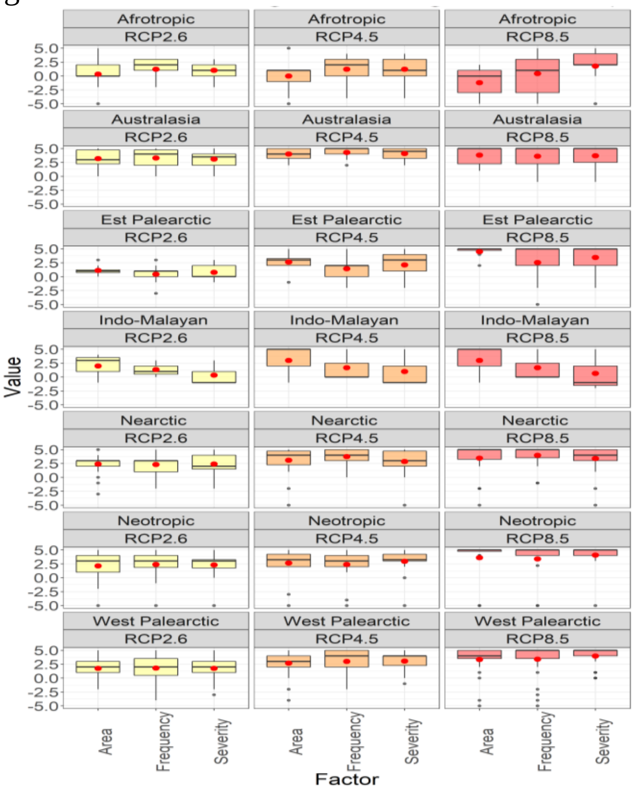

c) Biome-2050

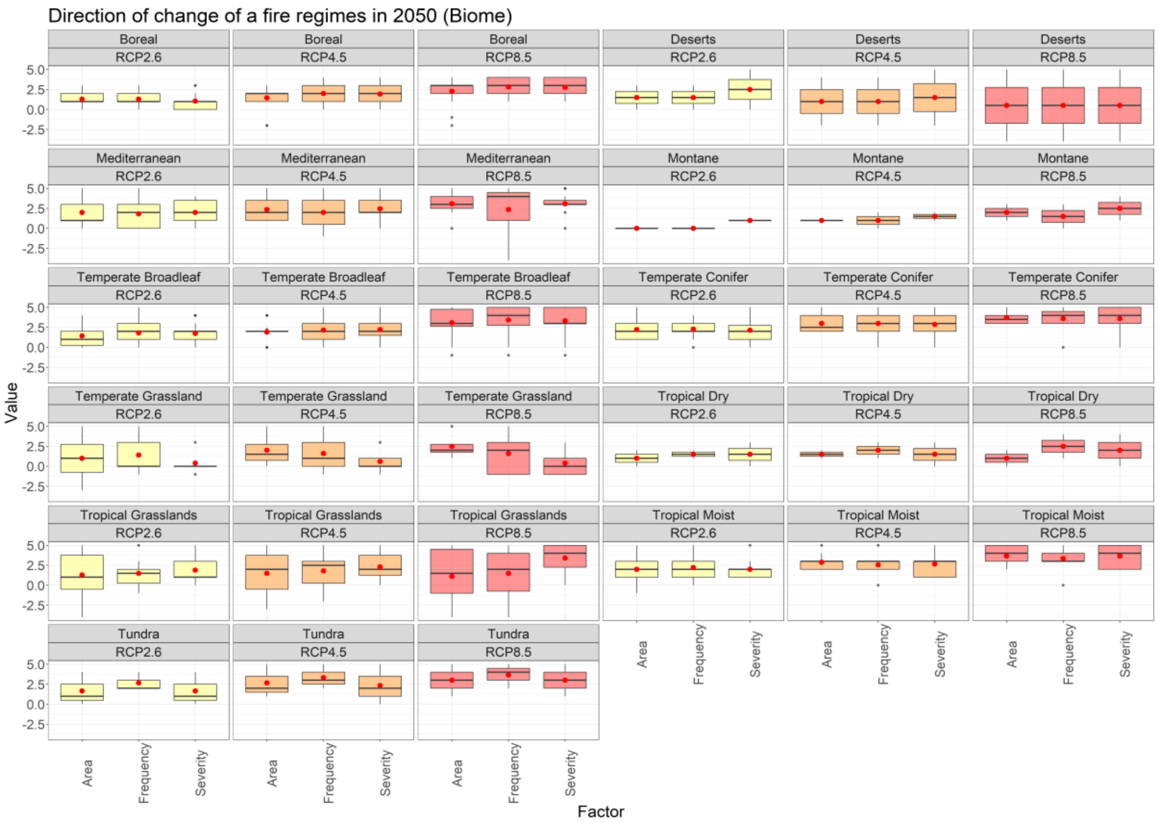

Biome-2100

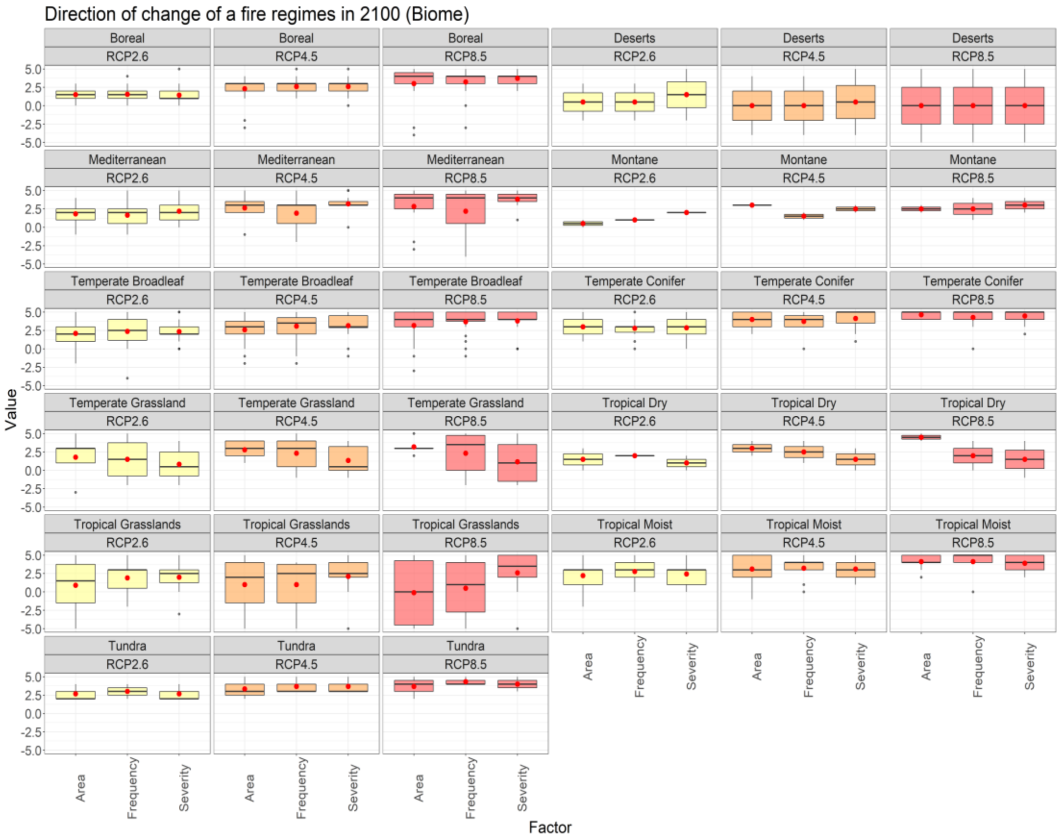

Biome- 2300

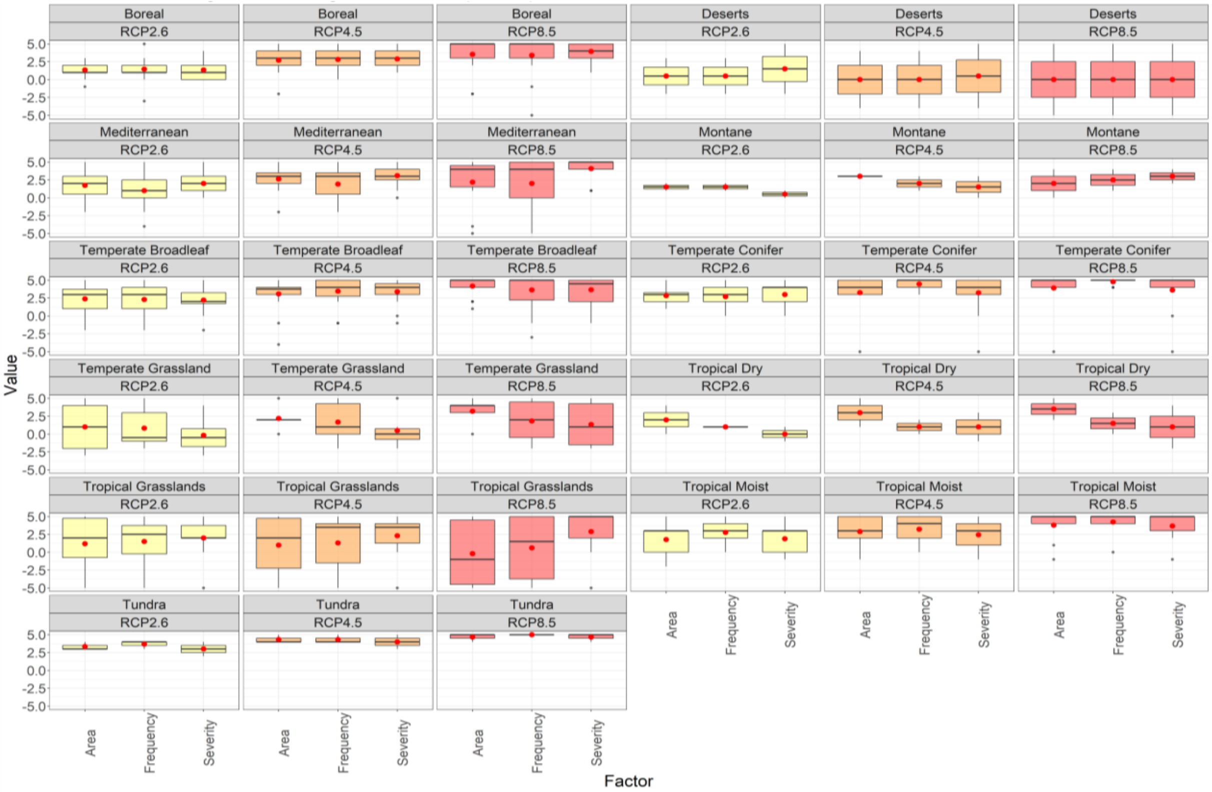

**Figure S13.** Fire regime characteristics direction change in the future. The boxplots represent the median (black line) and average values (red dots) of experts estimate of net change of different ecosystem values under three different RCP scenario for the year 2100, (-5 = strong net decrease, 0 = no net effect, 5= strong net increase).

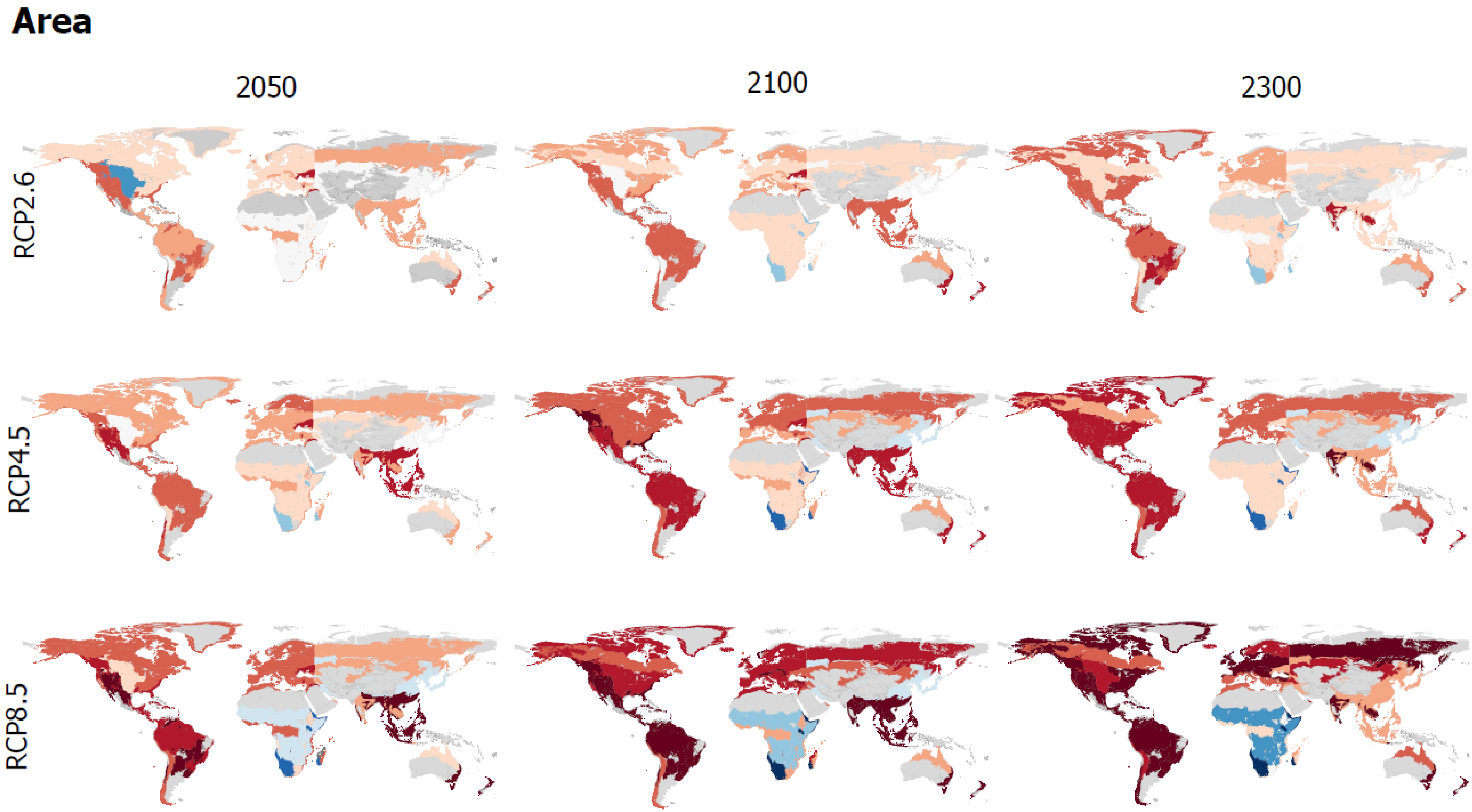

**Figure S14**. Median values of burned area change until 2050-2300 and three RCP scenario

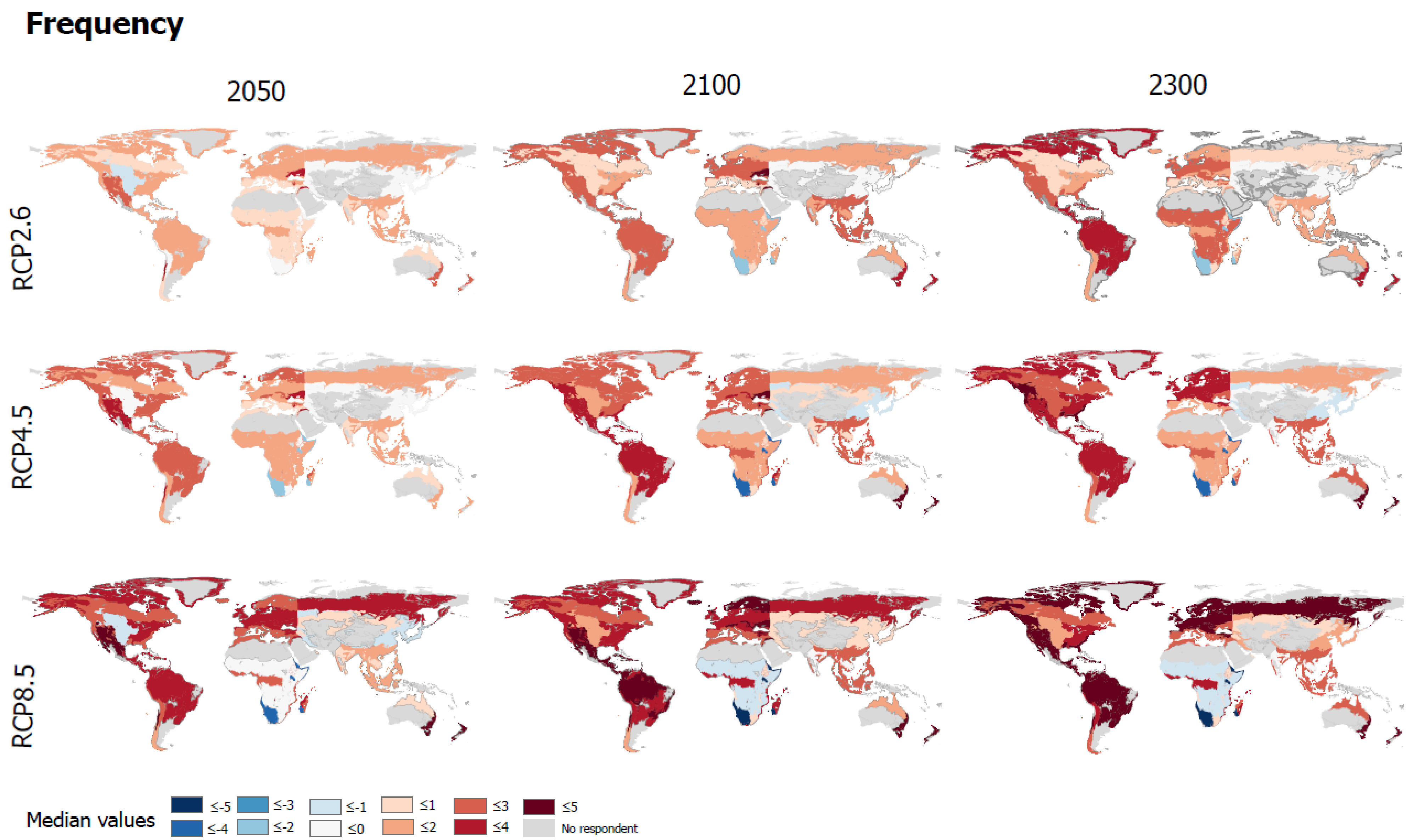

**Figure S15.**Median values of fire frequency change until 2050-2300 and three RCP scenarios

### Severity

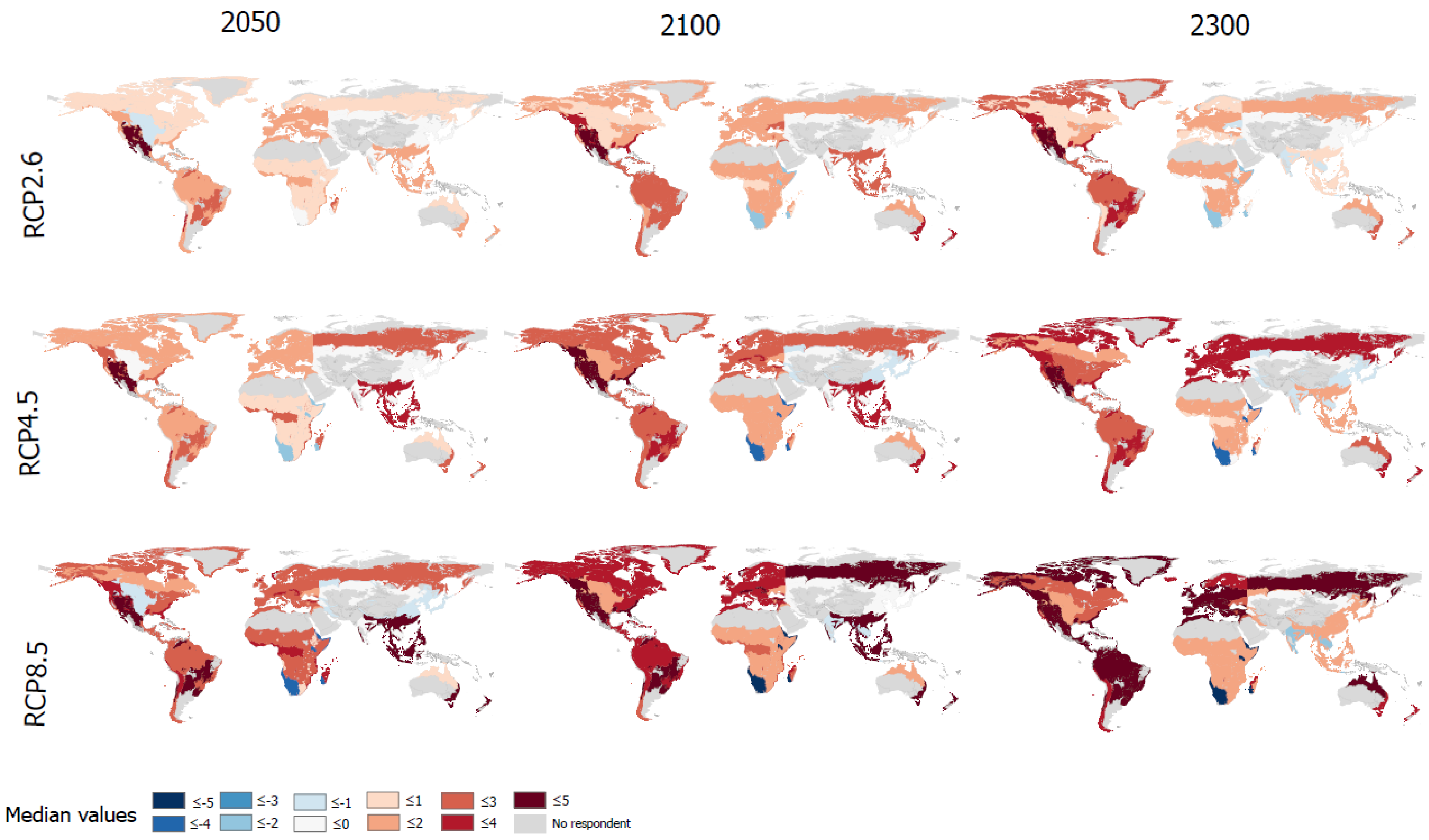

**Figure S16**. Median values of fire severity change until 2050-2300 and three RCP scenario

Ecosystem values change in the future as a consequence of fire regime change

d) Global

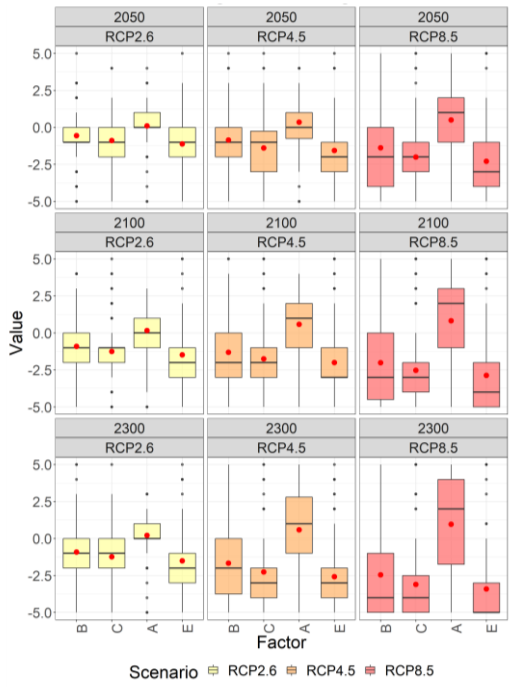

e) Region -2050

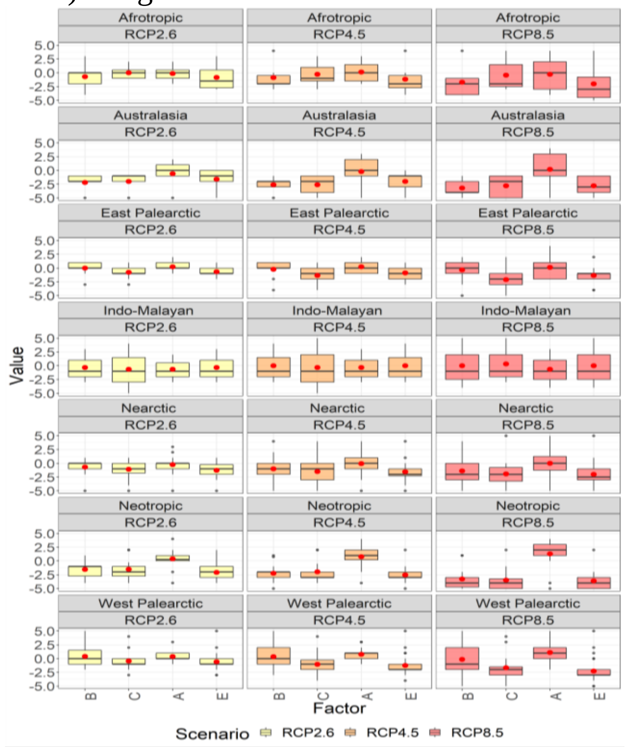

Region-2100

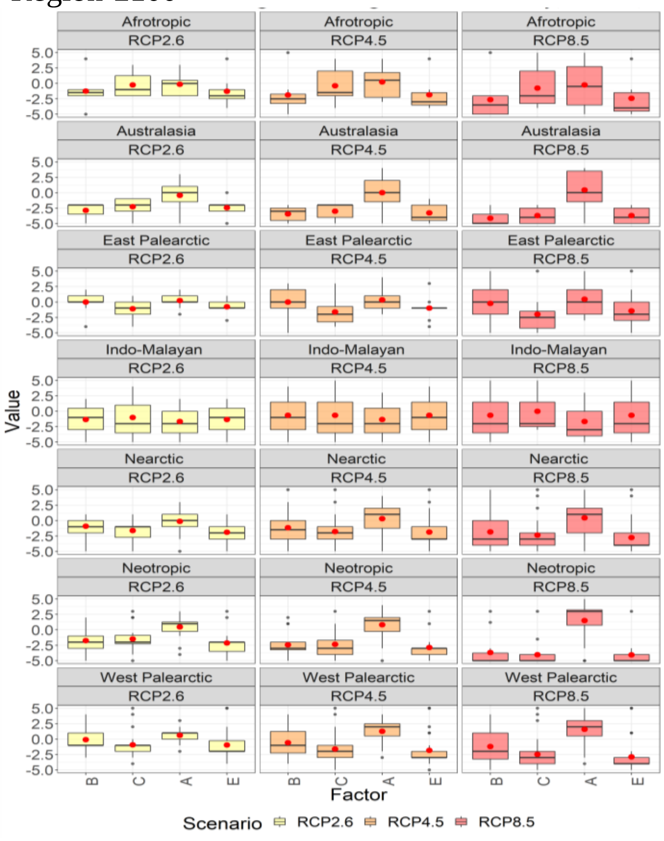

Region-2300

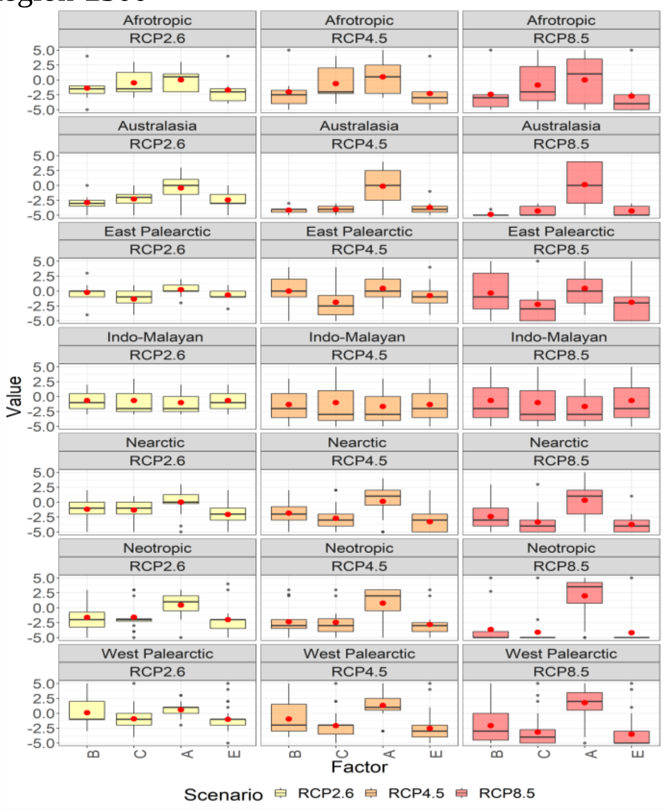

f) Biome-2050

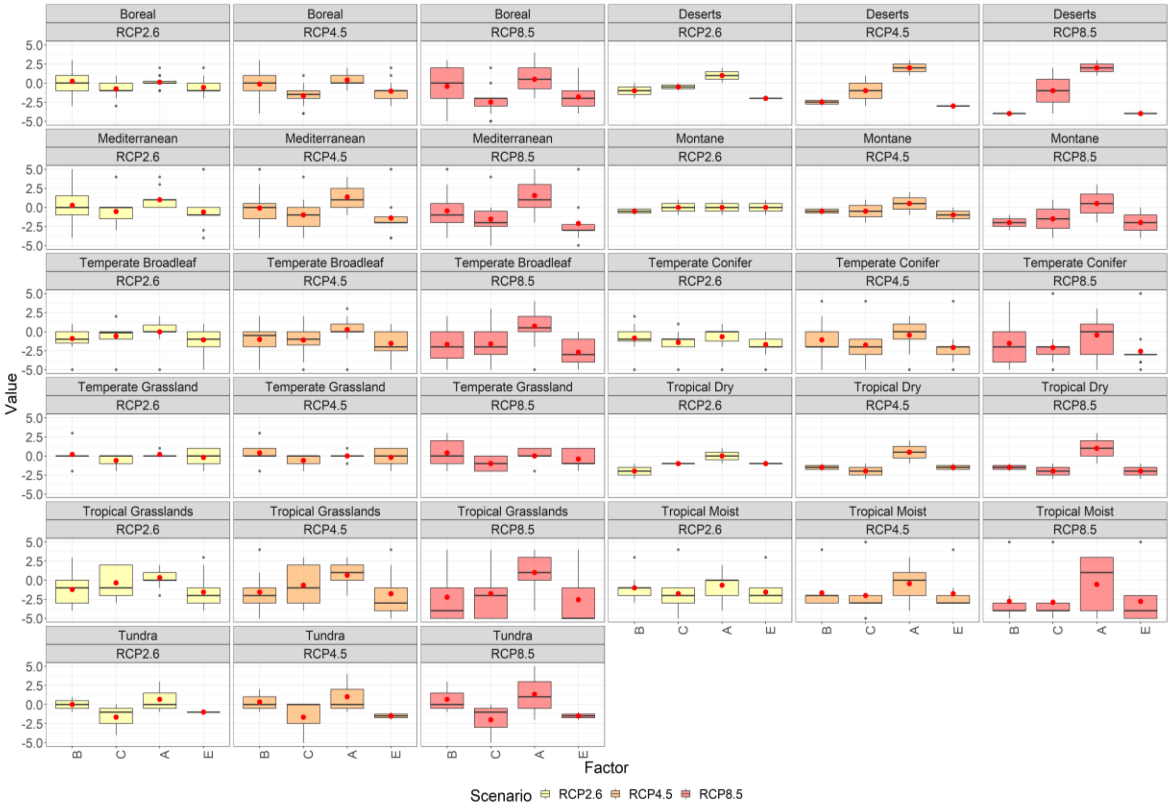

Biome-2100

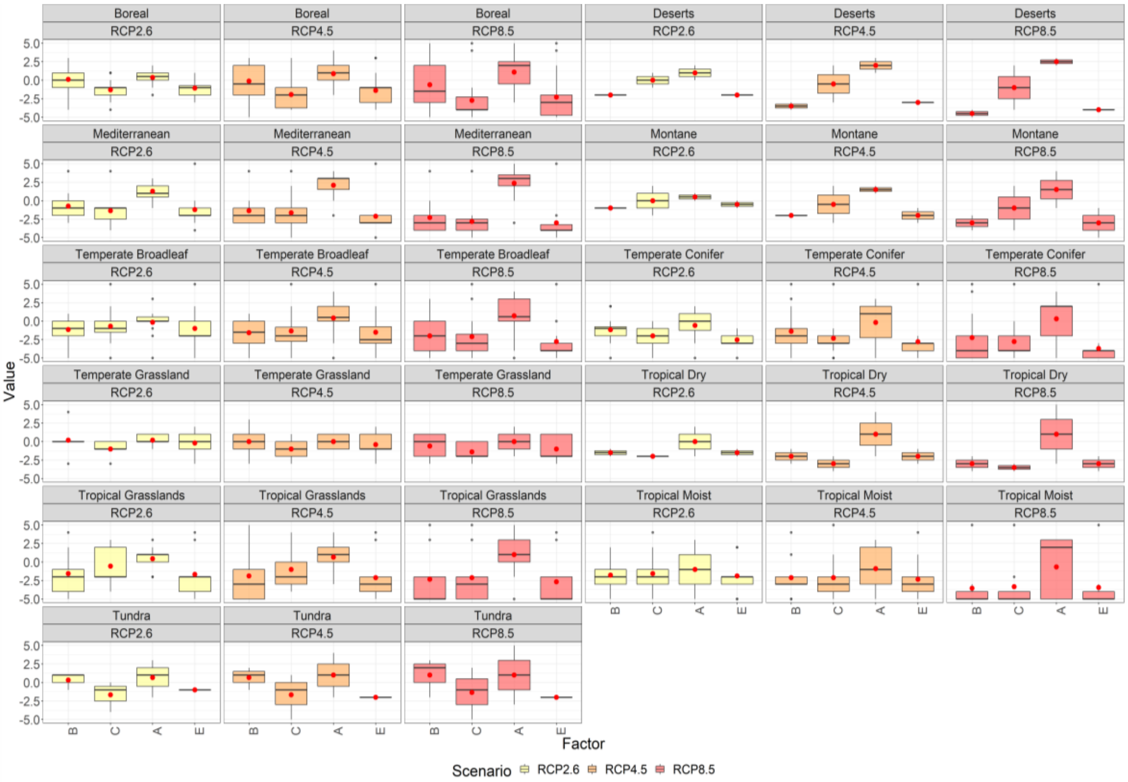

Biome- 2300

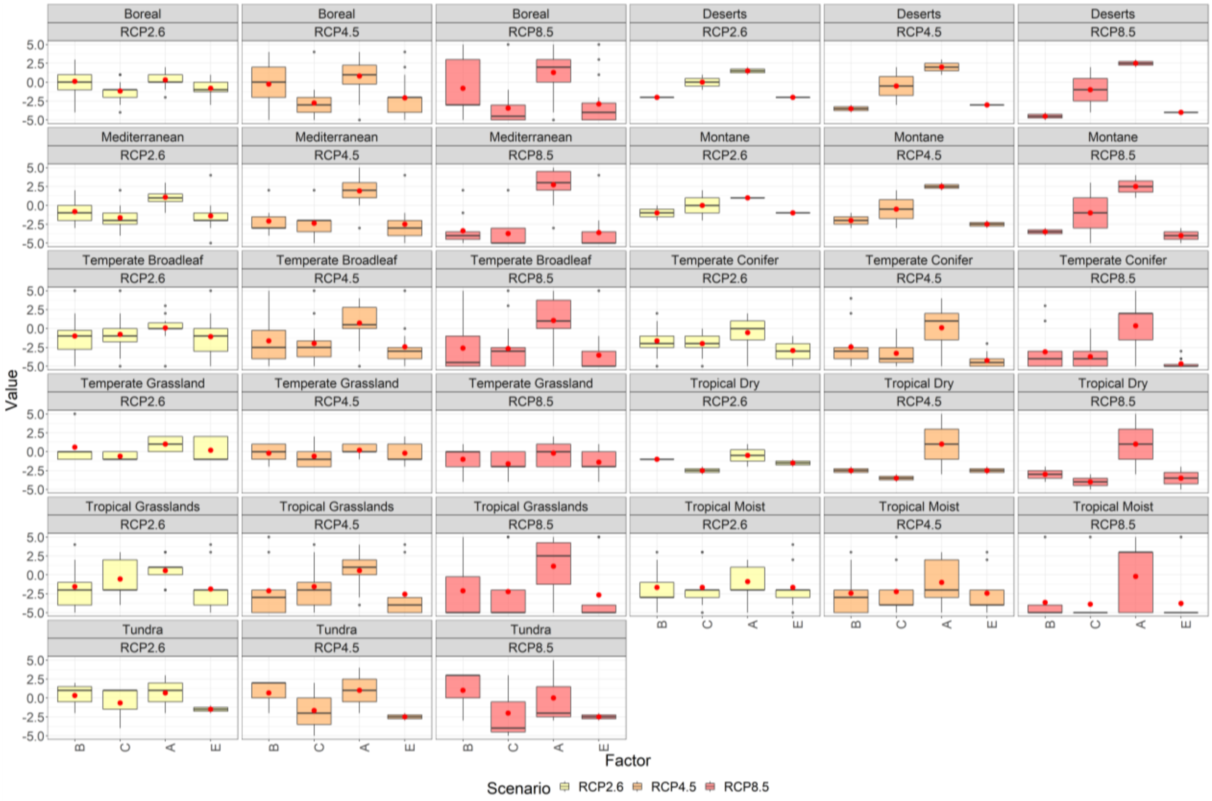

**Figure S17.** Net effect change of biodiversity (B), carbon stocks (C), albedo (A), ecosystem services (E) as a consequence of fire regime change for years 2050-2300 and under RCP 2.6-4.5-8.5. The boxplots represent the median (black line) and average values (red dots) of experts estimate of net change of different ecosystem values under three different RCP scenario for the year 2100, (-5 = strong net decrease, 0 = no net effect, 5= strong net increase).

#### Biodiversity

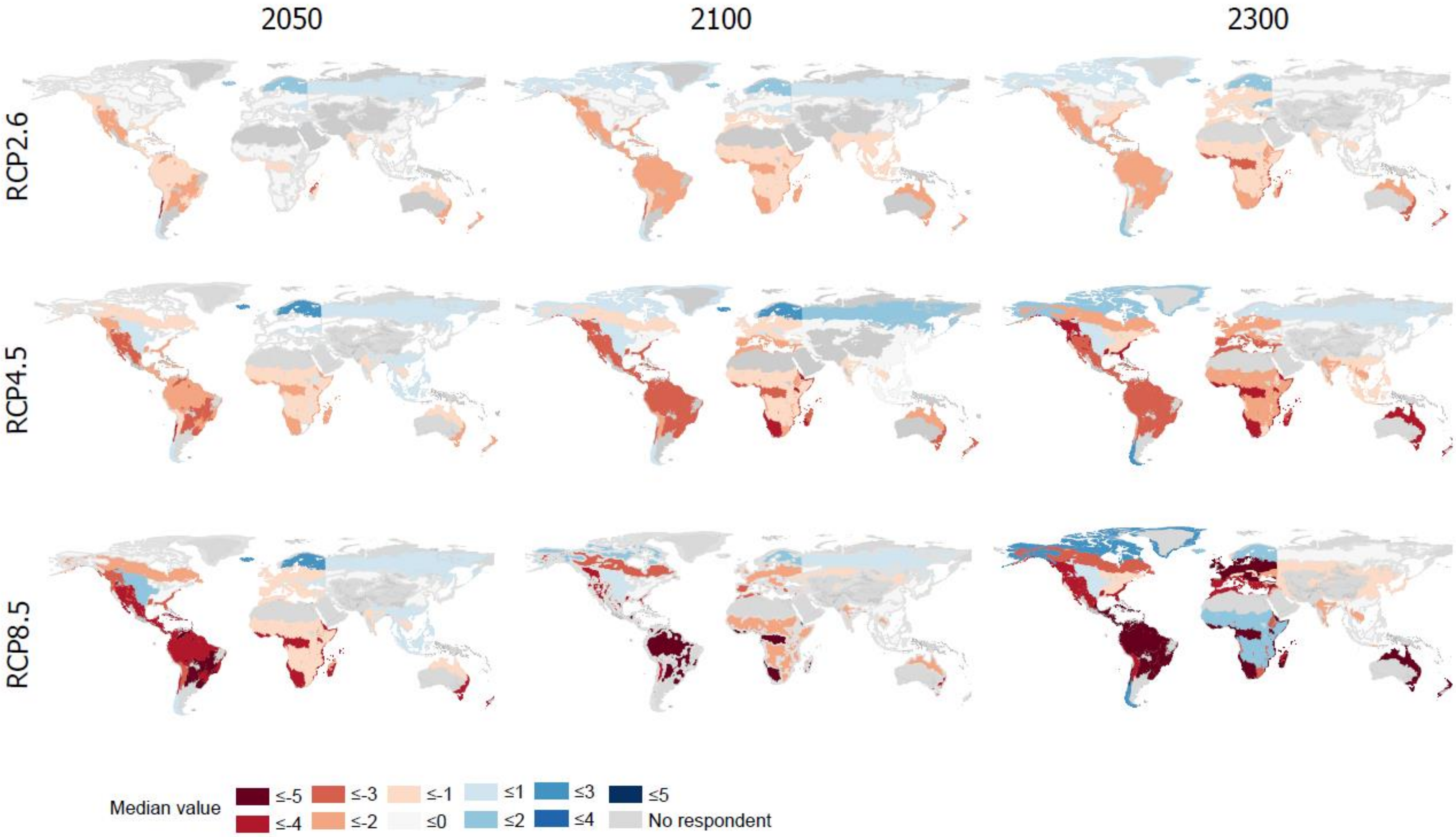

**Figure S18.** Median values of biodiversity change as a consequence of fire regime change until 2050-2300 and three RCP scenario

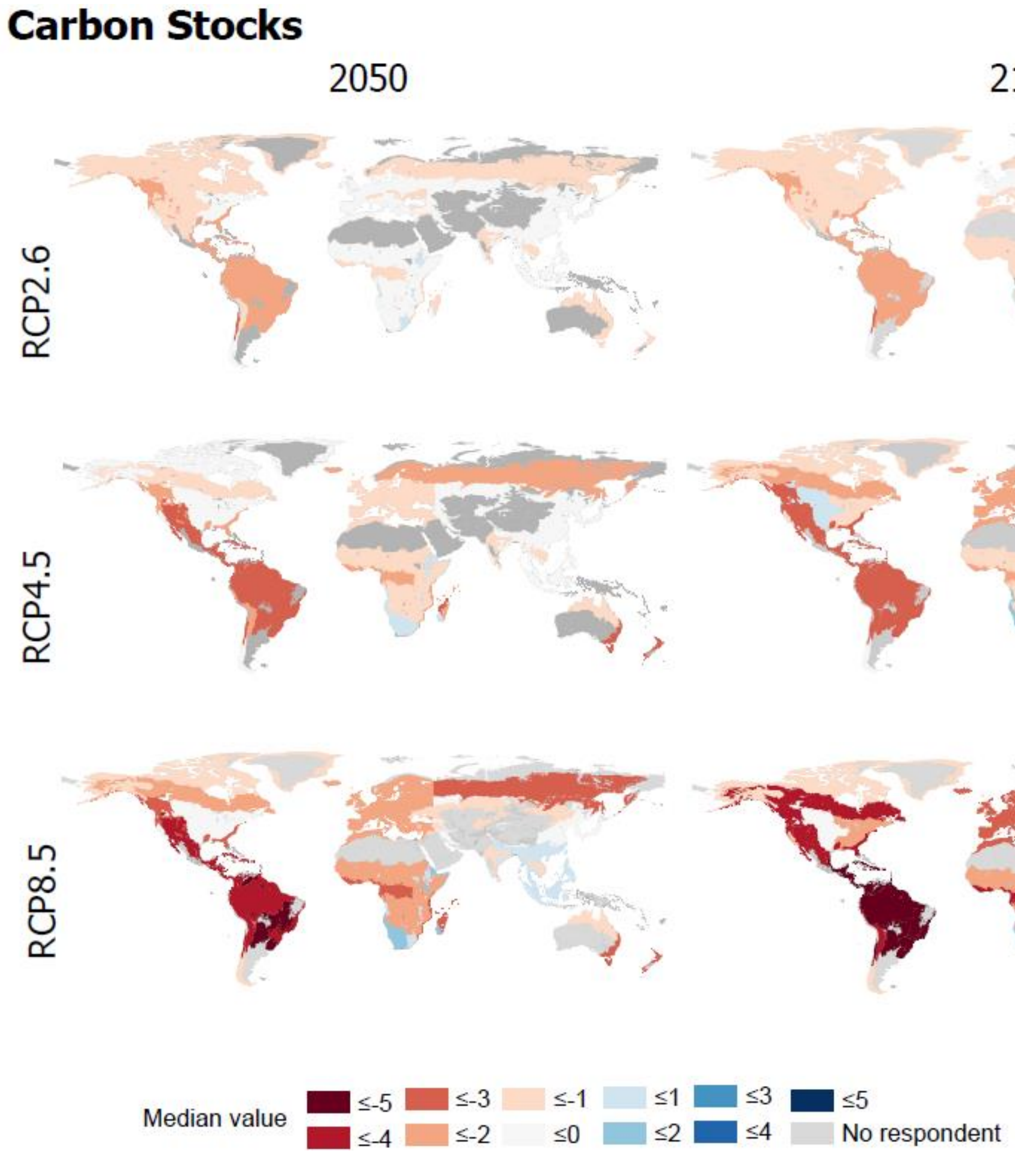

**Figure S19.** Median values of carbon stocks change as a consequence of fire regime change until 2050-2300 and three RCP scenarios

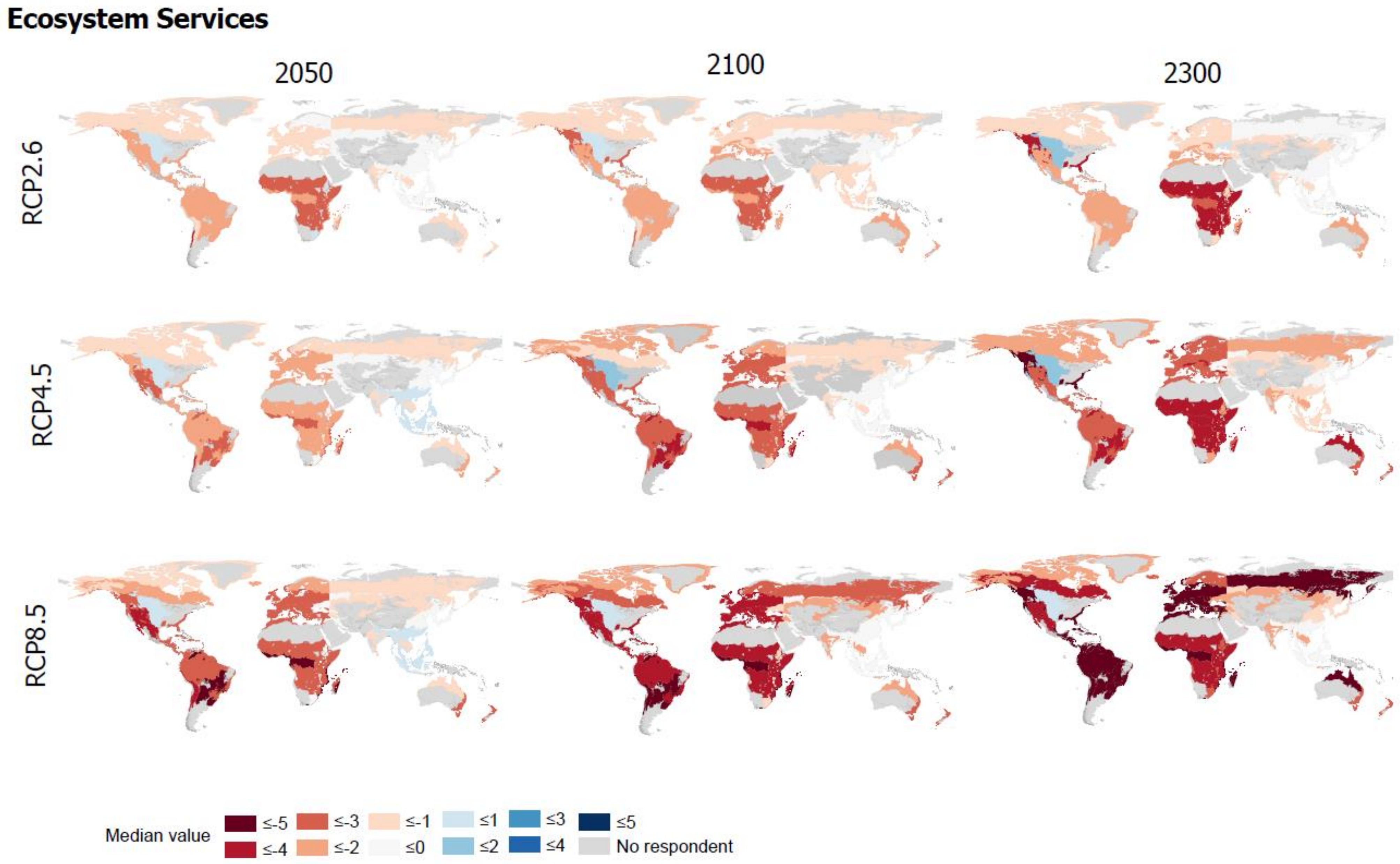

**Figure S20.** Median values of ecosystem service change as a consequence of fire regime change until 2050-2300 and three RCP scenario

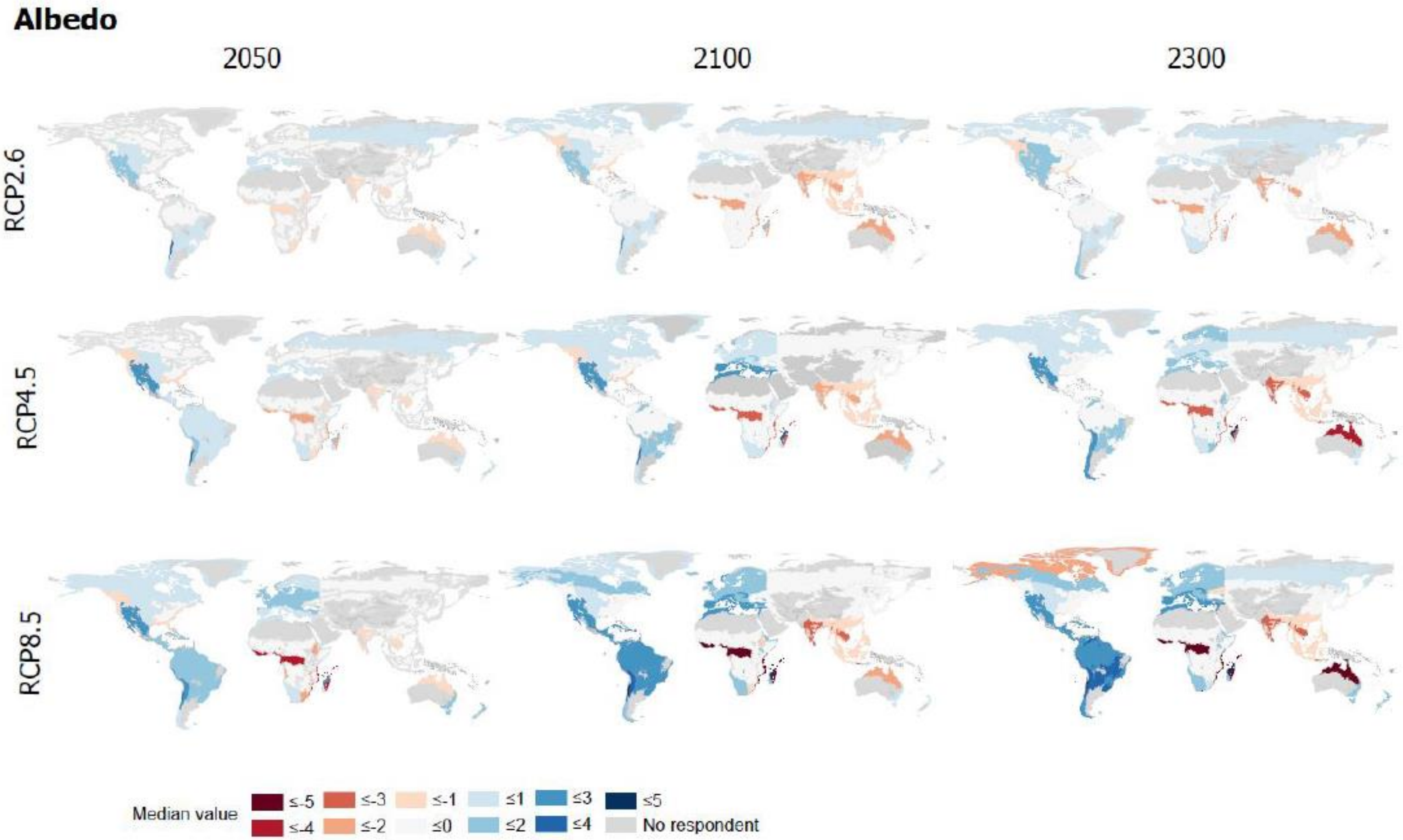

**Figure S21.** Median values of albedo change as a consequence of fire regime change until 2050-2300 and three RCP scenario

Drivers of fire regime change

a) Global

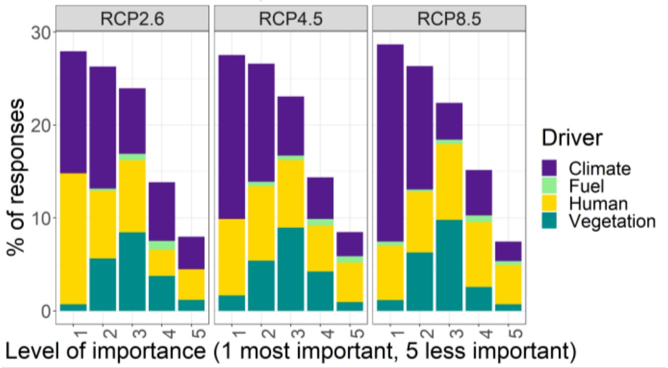

b) Region

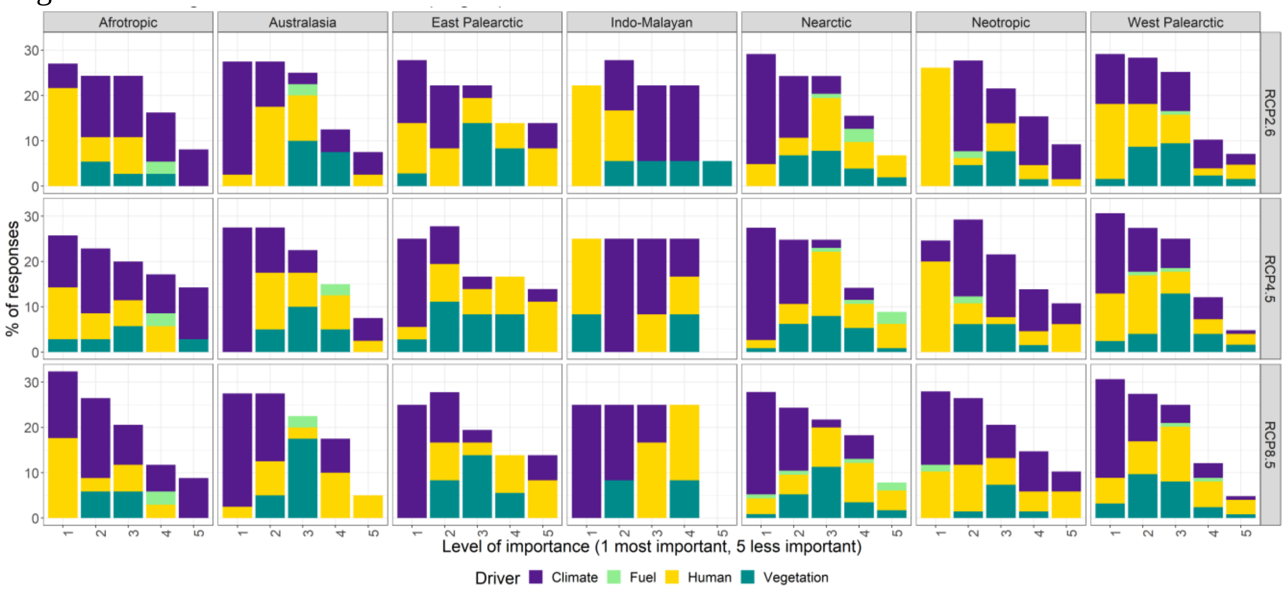

c) Biome

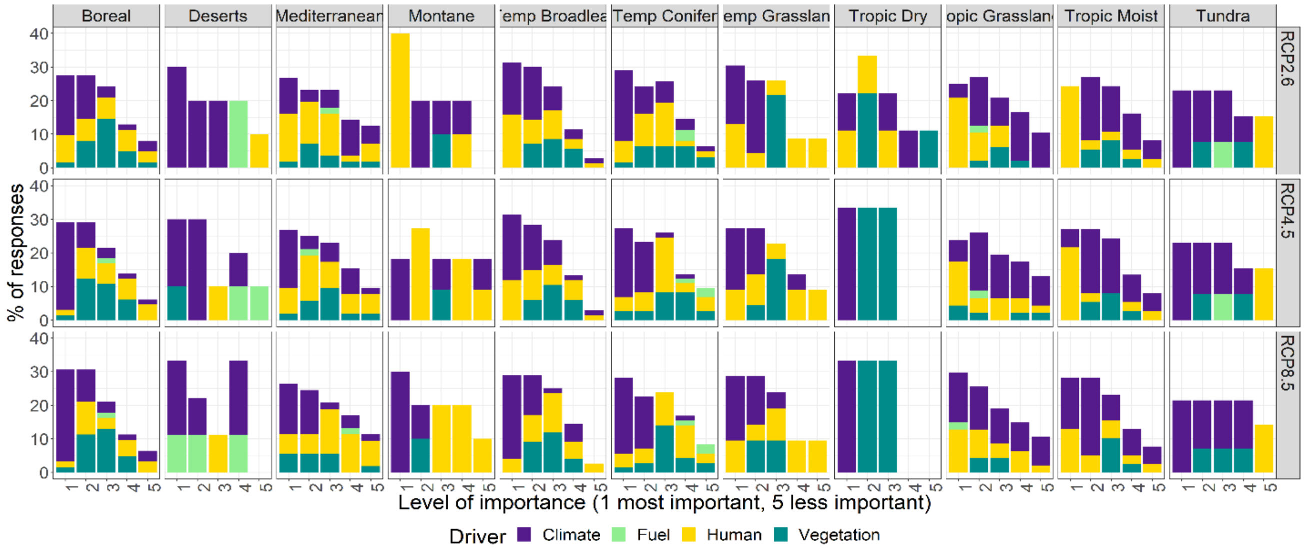

**Figure S22.**Identified drivers of fire regime change under three RCP scenarios by year 2100 in order of importance.

#### Human intervention effectiveness

##### a) Region

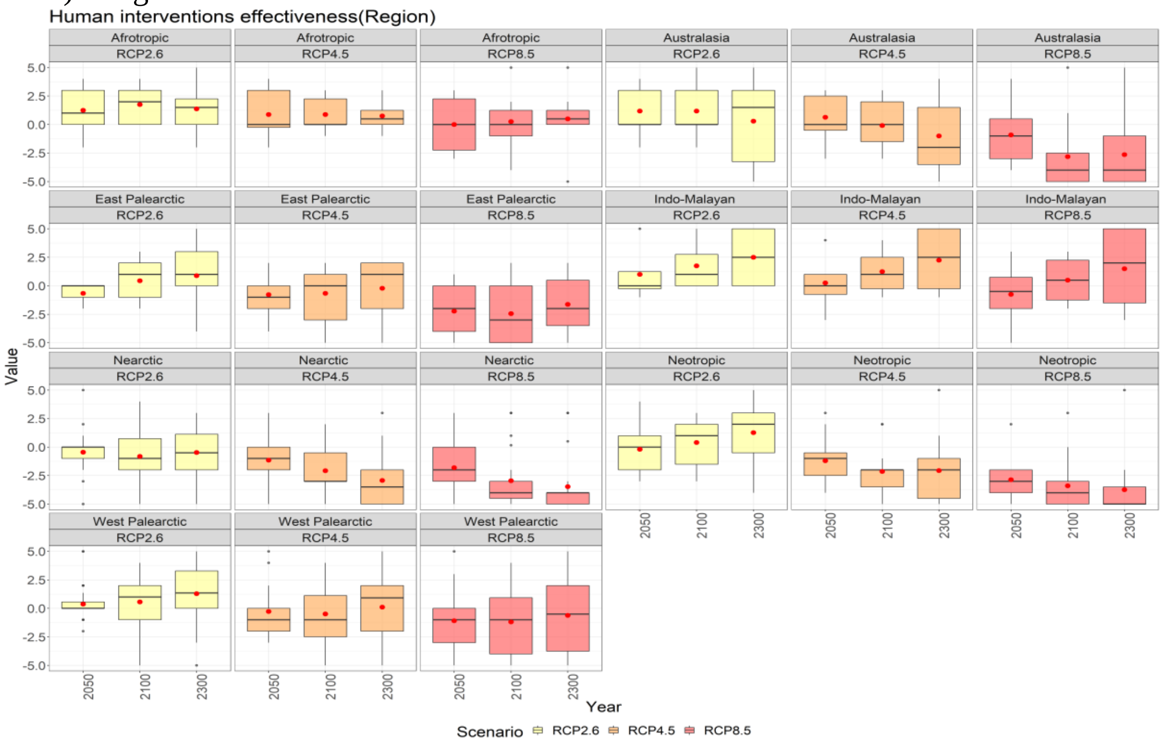

##### b) Biome

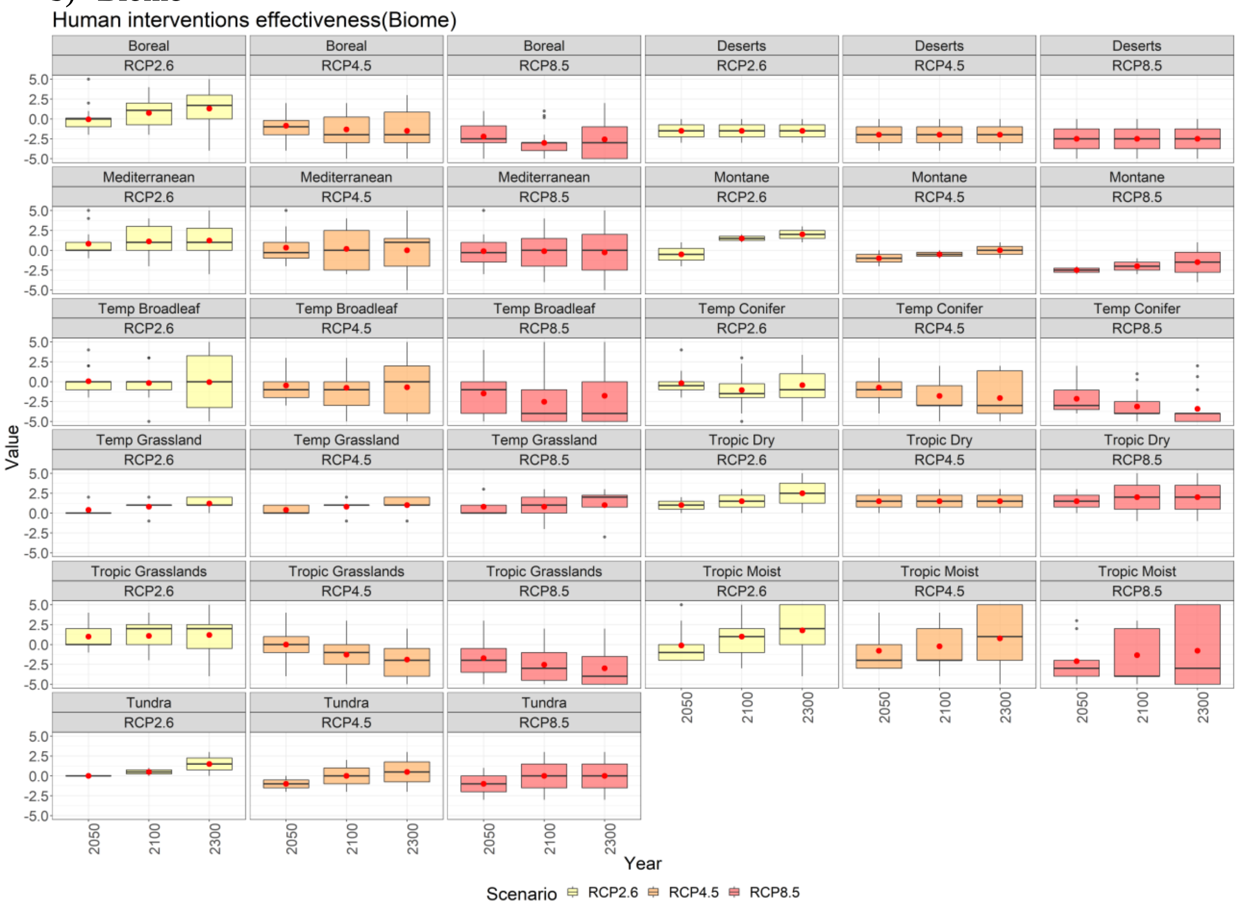

**Figure S23.** Effectiveness of human interventions (identified in table S1) in mitigating potential damage to societies and ecosystems for the three RCP scenarios for 2050-2300. where -5 = much less capacity to control fire regime for this scenario and period, 0 = same capacity as today, and 5 = much stronger capacity

**Table S1.** Summarized management and intervention action

| Action | Description and examples | Biodiversity | Carbon St | Albedo | Eco Service |
|---|---|---|---|---|---|
| **Fuel treatment (14%)** | • Prescribed burning: Indigenous burning-controlled burning (10.2% of the responses) | 3 (median value) | 2 | 0 | 3 |
| | • Fuel management: Fuel removal by harvesting /cleansing; Fuel breaks and assisted migration/planting (3.2%) | 2 | 1 | 0 | 2 |
| **Vegetation (23%)** | • Forest-native vegetation restoration –Reducing exotic invasive Species-Reforestation/ Stop deforestation, etc. (17.5%) | 3 | 3 | 3 | 3 |
| | • Selecting nonflammable species – Plant more resilient trees, etc. (4.3%) | -2 | 1 | 0 | 1 |
| **Landscape (7%)** | • E.g. Enhancing heterogeneous landscapes-Creating fire-resilient landscapes by reducing the spatial extent and continuity of forests and tall shrublands (5%) | 3 | 2 | 2 | 3 |
| | • Limiting wildland urban interface (2%) | 3 | 1.5 | 0 | 2 |

| | | | | | |
|---|---|---|---|---|---|
| **Improved Agriculture and sustainability**<br><br>**(5%)** | • E.g. Reduction in area of cropland in marginal areas- stop using fire in agriculture practices; Better prevention of agricultural fire expanding into the forested area, Promoting agroforestry systems, etc. (5%) | 3 | 3 | 2 | 3 |
| **Direct fire response**<br><br>**(17%)** | • Fire suppression (11%) | -1 | 1 | 0 | 1 |
| | • Fire management: E.g. Increased firefighting capability and deployment for forest fires (5%) | 2 | 1 | 0 | 2 |
| **Social - infrastructure intervention (23%)** | • Climate change mitigation (5%) | 4 | 5 | 3 | 4.5 |
| | • Policy: E.g. Promoting conservation policy- Incorporate indigenous knowledge and practices- Regulating economic activities – Preventing human ignition – Monitoring Long-term and real-time (early detection); water and waste management;, etc. (13%) | 3 | 3 | 1 | 3 |
| | • Education (4%) | 2 | 2 | 0 | 4 |
| **Non - Intervention (11%)** | • Non Intervention (11%) | 0 | -1 | 0 | -2 |

**Table S2.** Composition and characteristics of expert respondents

| Survey Section | Paleo perspective | Current fire regime state | Future fire projection | Intervention and management |
|---|---|---|---|---|
| **Response per region** | | | | |
| **Afrotropic** | 9 | 9 | 8 | 8 |
| **Australasia** | 15 | 11 | 10 | 11 |
| **East Palearctic** | 9 | 10 | 8 | 9 |
| **Indo-Malyan** | 4 | 3 | 4 | 3 |
| **Nearctic** | 31 | 24 | 24 | 22 |
| **Neotropic** | 16 | 16 | 16 | 16 |
| **West Palearctic** | 36 | 31 | 30 | 28 |
| **Average modeling/field self-rating[a]** | 2.36 | 2.36 | 2.38 | 2.32 |
| **Combined years of experience** | 1224 | 1176 | 1124 | 1003 |
| **Average years of experience** | 10 | 11 | 11 | 10 |
| **Ratio male: female** | 46:43 | 44:41 | 43:36 | 41:32 |

[a]1 was defined as exclusively field research and 5 as exclusively modeling research.

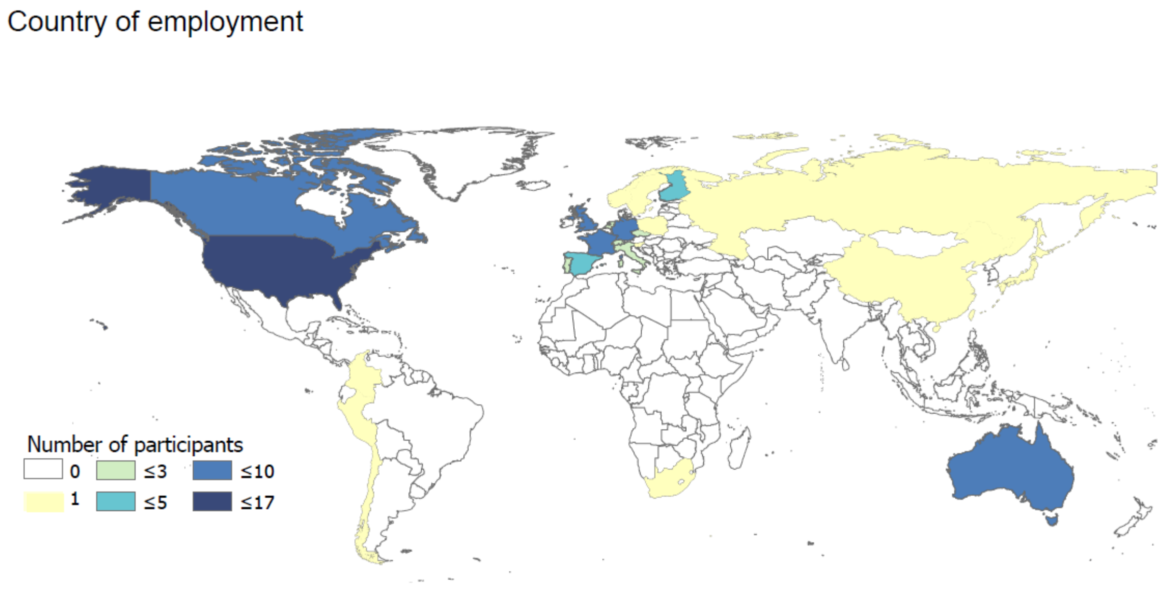

**Figure S24.** Respondents country of employment

**Table S3.** Mean self-reported confidence and expertise level for each section

<table>
<tr><th>Section</th><th>Mean expertise level</th><th>Mean confidence level</th><th>Estimates generate base</th></tr>
<tr><td>Paleo perspective</td><td>3.7</td><td>3.5</td><td>Published empirical data: 101<br>Published model estimates:31<br>Unpublished data:51<br>Professional opinion:68</td></tr>
<tr><td>Current fire regimes</td><td>2.8</td><td>2.7</td><td>Published empirical data:84<br>Published model estimates:32<br>Unpublished data:16<br>Professional opinion:69</td></tr>
<tr><td>Future projections</td><td>2.5</td><td>2.3</td><td>Published empirical data:50<br>Published model estimates:52<br>Unpublished data:18<br>Professional opinion:83</td></tr>
<tr><td>Management and intervention</td><td>2.4</td><td>2.3</td><td>Published empirical data:37<br>Published model estimates:22<br>Unpublished data:14<br>Professional opinion:71</td></tr>
<tr><td colspan="4">The five-point “Confidence level” scale is defined as follows:<br>1 = My answer is my best guess, but I am not confident in it; it could easily be far off the mark.<br>2 = My answer is an educated guess; it could be far off the mark, but I have some confidence in it.<br>3 = I am moderately confident in my answer; it is not precise, but it may be near the true value.<br>4 = I am confident in my answer; the true value is likely to be somewhat different from my answer, but it is unlikely to be dramatically different.<br>5 = Given current understanding, I would be surprised if my answer were far off from the true value.</td></tr>
<tr><td colspan="4">The five-point “Expertise level” scale is defined as follows:<br>1 = I have little familiarity with the literature, and I do not actively work on this area.<br>2 = I have some familiarity with the literature, and I’ve worked on related questions but haven't contributed to the literature on this issue; it is not an area of central expertise for me.<br>3 = I have worked on related issues and have contributed to the relevant literature but do not consider myself an expert on this issue.<br>4 = I am very familiar with relevant literature and have worked on related questions. This is an area of central expertise for me.<br>5 = I contribute actively to the literature directly concerned with this issue, and I consider myself one of the foremost experts on it.</td></tr>
</table>

#### Affiliations

[1]Brigham Young University, Department of Plant and Wildlife Sciences. Provo, Utah, USA

[2]Chrono-environnement, UMR 6249 CNRS, Université Bourgogne Franche-Comté, Besançon, France;

[3]MSHE, UAR 3124 CNRS, Université Bourgogne Franche-Comté, Besançon, France;

[4]Institute of Plant Sciences and Oeschger Centre for Climate Change Research, University of Bern, Bern, Switzerland

[5]Aix Marseille Univ, Avignon Univ, CNRS, IRD, IMBE, Aix-en-Provence, France

[6]Centre for Quaternary Research (CQR), Department of Geography, Royal Holloway University of London (RHUL), Egham, Surrey , UK

[7]Dept. of Ecology Philipps-Marburg University, Marburg, Germany.

[8]Past Landscape Dynamics Laboratory, Institute of Geography and Spatial Organization, Polish Academy of Sciences, Warsaw, Poland

[9]Département de Géographie,Université de Montréal

[10]Department of Physical Geography, Goethe University, Altenhöferallee 1, 60438 Frankfurt am Main, Germany

[11]STAR-UBB Institute Babeș-Bolyai University, Kogălniceanu 1, 400084, Cluj-Napoca, România

[12]Natural Resources Canada, Canadian Forest Service, Canada

[13]University of British Columbia, Department of Earth, Environmental and Geographic Sciences, Kelowna, Canada

[14]Department of Forest Sciences, University of Eastern Finland, Joensuu, Finland

[15]Department of Archaeology, Turku Institute for Advaned Studies (TIAS), University of Turku, 20014 Turku, Finland

[16]Geography, Planning, and Spatial Sciences, University of Tasmania, Hobart, Australia;

[17]School of Ecosystem and Forest Sciences, University of Melbourne, Richmond, Australia;

[18]ARC Centre of Excellence for Australian Biodiversity and Heritage

[19]ZRC SAZU, Institute of Archaeology, Ljubljana, Slovenia

[20]Institut de recherche sur les forêts, Université du Québec en Abitibi-Témiscamingue, Rouyn-Noranda, QC J9X 5E4, Canada

[21]Département des sciences biologiques, Université du Québec à Montréal, Montréal, QC H2X 1Y4, Canada

[22]Albrecht-von-Haller Institute, Department of Palynology and Climate Dynamics, University of Goettingen

[23]School of Environmental Sciences, University of Liverpool, Liverpool, UK

[24]The Institute of Evolutionary Science of Montpellier

[25]Department of Geography & Planning, School of Environmental Sciences, University of Liverpool, L69 7ZT Liverpool, UK

[26]Department of Environmental Sciences, University of Basel, Switzerland

[27]Department of Geography, University of Oregon, Eugene, Oregon, USA

[28]ARC Centre of Excellence for Australian Biodiversity and Heritage, School of Earth and Atmospheric Sciences, University of Wollonong, NSW, Australia

[29]University of Padova, Department of Land, Environment, Agriculture and Forestry (TESAF), Italy;

[30]ISEM, University of Montpellier, CNRS, IRD, EPHE, Montpellier, France

[31]EPHE-PSL University

[32]UMR LEHNA, Université Claude Bernard Lyon, Villeurbanne, France

33Department of Botany, Faculty of Science, Charles University, Prague, Czech Republic

[34]CEABN/InBIO – Centre for Applied Ecology / Research Network in Biodiversity and Evolutionary Biology, School of Agriculture, University of Lisbon, Tapada da Ajuda, 1349-017 Lisboa, Portugal

[35]ARC Centre of Excellence for Australian Biodiversity and Heritage, Global Ecology, College of Science and Engineering, Flinders University, Adelaide, SA, Australia

[36]Eco&Sols, Univ Montpellier, CIRAD, INRAE, Institut Agro, IRD, Montpellier, France

[37]CIRAD, UMR Eco&Sols, Montpellier, France

[38]Laurentian Forestry Centre, Canadian Forest Service, Natural Resources Canada, Quebec City, QC, G1V 4C7, Canada;

[39]Atlantic Forestry Centre, Canadian Forest Service, Natural Resources Canada, Fredericton, NB, E3B 5P7, Canada

[40]Syracuse University, Department of Earth and Environmental Sciences, Syracuse, NY, USA

[41]Swiss Federal Institute for Forest, Snow and Landscape Research WSL, Insubric Ecosystems Research Group, A Ramél 18, CH-6593 Cadenazzo, Switzerland

[42]School of Culture, History & Language, and ARC Centre of Excellence for Australian Biodiversity and Heritage, Australian National University, Canberra ACT-2601, Australia

[43]University of New South Wales, School of Biological, Earth and Environmental Sciences, Sydney, Australia

[44]Geoecology, Department of Environmental Sciences, University of Basel, Klingelbergstrasse 27, 4056 Basel, Switzerland;

[45]Center for Water Infrastructure and Sustainable Energy (WISE) Futures, Nelson Mandela African Institution of Science & Technology, P.O. Box 9124 Nelson Mandela, Tengeru, Arusha, Tanzania

[46]York Institute for Tropical Ecosystems, Department of Environment and Geography, University of York, Wentworth Way, York, YO10 5DD, United Kingdom

[47]School of Animal, Plant & Environmental Sciences, University of the Witwatersrand, Braamfontein, 2000, South Africa

[48]University of Massachusetts Amherst. Department of Geosciences. Amherst, Massachusetts, USA

[49]Departmento de Estratigrafía y Paleontología, Facultad de Ciencias, Universidad de Granada, Granada, Spain

[50]Polar Terrestrial Environmental Systems, Alfred Wegener Institute Helmholtz Center for Polar and Marine Research, Potsdam, Germany;

[51]Institute of Geography, Georg-August-University Göttingen, Goldschmidtstr. 5, 37077 Göttingen, Germany

[52]Mediterranean Ecogeomorphological and Hydrological Connectivity Research Team (http://medhycon.uib.cat), Department of Geography, Universitat de les Illes Balears, Carretera de Valldemossa km 7.5, 07122 Palma, Balearic Islands, Spain;

[53]Institute of Agro-Environmental and Water Economy Research –INAGEA (http://inagea.com), Universitat de les Illes Balears, Carretera de Valldemossa km 7.5, 07122, Palma, Balearic Islands, Spain

[54]University of Trás-os-Montes and Alto Douro, Centre for the Research and Technology of Agro-Environmental and Biological Sciences, Vila Real, Portugal

[55]ISEM, University of Montpellier, CNRS, EPHE, IRD, Montpellier, France

[56]Department of Biological Sciences, University of Bergen, PO Box 7803, 5020 Bergen, Norway

[57]Australian Bureau of Meteorology: Hobart, Tasmania, Tas, AU

[58]Center for Climate and Resilience Research (CR)2 & Institute of Ecology and Biodiversity (IEB), Chile

[59]Laurentian Forestry Centre, Canadian Forest Service, Natural Resources Canada, Quebec City, QC, G1V 4C7, Canada;

[60]Brigham Young University, Department of Public Health, Provo, UT, USA

[61]Departamento de Ciencias Biológicas, Universidad de los Andes, Bogotá-Colombia

[62]Weber State University, Department of Geography, Environment, and Sustainability, Ogden, UT, USA

[63]Max Planck Institute for the Science of Human History, Jena, Germany

[64]Marine, Earth, and Atmospheric Sciences, North Carolina State University, Raleigh, NC, 27695, USA

[65]Earth System Science Program, Faculty of Natural Sciences, Universidad del Rosario, Colombia

[66]School of the Environment, Geography and Geosciences, University of Portsmouth, Buckingham Building, Lion Terrace, Portsmouth, PO1 3HE, UK

[67]School of Geography and Sustainable Development, Irvine Building, University of St Andrews, St Andrews, Scotland, UK (ROW 57)

[68]Department of Forest and Conservation Sciences, University of British Columbia, Vancouver, British Columbia, Canada

[69]Earth Lab (CIRES), University of Colorado, Boulder, CO, USA

[70]Department of Gesciences, Osaka Metropolitan University, Osaka, Japan

[71]Department of Ecology and Evolutionary Biology, Yale University, New Haven, CT, USA

[72]Department of Earth and Environmental Sciences, The Graduate Center of CUNY, 365 Fifth Avenue, New York, NY 10016, USA

[73]Department of Geosciences and Geography, University of Helsinki, Helsinki, Finland

[74]Department of Biology and Centre for Forest Biology, University of Victoria, Victoria, BC, Canada

[75]Institut of Evolution Sciences, University of Montpellier CNRS IRD EPHE, Montpellier, France

[76]Univ. Rennes 1, CNRS, ECOBIO UMR 6553, Rennes, France

[77]Department of Geography, University of Wisconsin Oshkosh, Oshkosh WI, USA

[78]Environmental Archaeology Research Group, Institute of History, CSIC, Albasanz 26-28, 28037 Madrid, Spain

[79]Department of Geography and Environmental Sciences, Northumbria University Newcastle, Newcastle upon Tyne, United Kingdom

[80]Biology Department, Luther College, Decorah, IA 55102, USA

[81]Swedish University of Agricultural Sciences, Sweden

[82]Forestry Science Institute (ICIFOR), INIA, CSIC, Madrid, Spain/University Polithechnic of Madrid, Spain

[83]Department of Archaeology, Max Planck Institute for Geoanthropology, Jena, Germany

[84]Climate Change Ecology Research Unit, Faculty of Geographical and Geological Sciences, Adam Mickiewicz University, Poznań, Poland

[85]Dept. of Earth & Planetary Sciences, University of New Mexico,Albuquerque NM

[86]School of Geography, University of Nottingham, NG7 2RD, Nottingham, UK

[87]Department of Earth Sciences, Montana State University, Bozeman, Montana, 59717 USA

[88]Department of Earth, Environment and Life Sciences (DISTAV), University of Genoa, Italy

[89]Geosciences Barcelona (GEO3BCN) CSIC, c/ Lluis Sole i Sabaris s/n, 08028 Barcelona, Spain

[90]School of Biological, Earth and Environmental Sciences, UNSW Sydney, NSW Australia

[91]Institute of Plant Sciences and Oeschger Centre for Climate Change Research, University of Bern, Altenbergrain 21, CH-3013, Bern, Switzerland

[92]Grupo de Ecología y Restauración Forestal, Departamento de Ciencias de la Vida, Facultad de Ciencias, Universidad de Alcalá, Alcalá de Henares, Spain

[93]Department of Geography, University of Utah, Salt Lake City, Utah, 84112 USA

[94]School of Earth and Environmental Sciences, The University of Queensland, Brisbane, Queensland 4072, Australia

[95]Environmental Change Institute, School of Geography and the Environment, University of Oxford, OX13QY Oxford, United Kingdom

[96]Forest Research Centre, School of Agriculture, University of Lisbon

[97]Institute of Geology and Palaeoontology, University of Münster, Germany

[98]CNR - Institute of Environmental Geology and Geoengineering (IGAG), Lab. of Palynology and Palaeoecology, Piazza della Scienza 1, 20126 Milano, Italy

[99]Interdisciplinary Laboratory for Continental Environments, CNRS, University of Lorraine, Campus Bridoux, 8 rue du Général Delestraint, 57070 Metz, France

[100]ARC Centre of Excellence for Australian Biodiversity and Heritage, College of Arts, Society and Education, James Cook University, Cairns, Australia

[101]Institute of Geography, Augsburg University, Augsburg, Germany

[102]NESSC - Netherlands Earth System Science Centre

[103]PaleoData Lab, Institute of Archaeology and Ethnography SB RAS, Novosibirsk, Russia

[104]Biological Institute, Tomsk State University, Tomsk, Russia

[105]Darwin Centre for Bushfire Research, Charles Darwin University, Darwin, Northern Territory, Australia

[106]Department of Palynology and Climate Dynamics, Georg-August-University of Göttingen, Wilhelm-Weber-Str. 2a, 37073 Göttingen, Germany

[107]Global Environment and Natural Resources Institute (GENRI)

[108]Department of Geography and Geoinformation Science

George Mason University"

[109]Division of Ecology, Department of Biology, Hacettepe University, Beytepe 06800 Ankara, Turkey

[110]Departments of Biology and Environmental Studies, St Olaf College, Northfield, Minnesota, 55057 USA

[111]Department of Systematic and Evolutionary Botany, University of Zurich, Zollikerstrasse 107, CH-8008 Zurich, Switzerland.

[112]Herbario Vargas CUZ, Universidad Nacional de San Antonio Abad del Cusco, Av. de La Cultura 773, Cusco, Perú.

[113]Geography, Faculty of Environment, Science and Economy, University of Exeter, United Kingdom

[114]Department of Geosciences, Auburn University, Auburn, USA

[115]Faculty of Biological and Environmental Sciences, University of Helsinki, Finland

[116]Institute of Tibetan Plateau Research, Chinese Academy of Sciences, Beijing, China; Chongqing University, Chongqing, 400044, China

[117]Univ. Bordeaux, CNRS, Bordeaux INP, EPOC, UMR 5805, F-33600 Pessac, France

#### Questionnaire

### Global assessment of abrupt change in fire regimes

**A project from the International Paleofire Network** (previously the Global Paleofire Working Group of Future Earth's Past Global Changes—PAGES—project)

#### 1. Introduction

You have been identified as a fire expert by our analysis of the literature, and we invite you to participate in this expert assessment project. Our goal is to document scientific opinion on changes in fire regimes and their effects on ecosystems, climate, and societies. We are focusing on centennial to millennial changes in past, present, and future fire regimes by region and biome. Because responding to the 15 questions below will likely take between 5 to 10 hours, all participants will have an opportunity to be co-authors on the resulting manuscript, which we will submit to Nature Geoscience in the fall of 2020. Previous expert assessments have been highly successful at combining available information and identifying knowledge gaps (Abbott et al., 2016; Schuur et al., 2011, 2013). Additionally, these efforts have repeatedly led to longer-term collaborations and new research activities (Bamber & Aspinall, 2013; Morgan, 2014).

We recognize that climate- and human-caused fire feedbacks in complex Earth systems are not, and cannot be, precisely and definitively modeled. As such, we are only asking for your informed opinion, realizing that some of the included parameters are not well understood. Possible thresholds and tipping points in the relationship between climate, land cover, land use changes, ecosystem structure, and fire regime are of particular interest, because such nonlinearity is difficult to predict with models. Combining assessments from multiple scientists with applicable and diverse expertise will allow an integrative evaluation of the range of possible futures, providing a valuable complement to projections from numerical models. We hope sincerely that you will participate if you are able.

#### 2. Definitions of fire regime & state change

##### *Fire regime*

While the term "fire regime" has multiple meanings in different fields, for the purposes of this survey we define fire regime in terms of spatial and temporal fire behavior (e.g. Hély et al., 2019; Keeley, 2009; Whelan, 1995). Spatial components of fire regime include the type of fire (e.g. heat and height) and its extent. Temporal components of fire regime include fire frequency, cycle, and seasonal timing. These variables interact to determine the effect of fire on ecosystem functioning and structure. For reference, we provide a short definition of common **aspects of fire regime** as described in the literature:

**Frequency**: Number of fires per unit of time in a defined area.
**Interval**: Time between two fire events in a defined area. In paleofire reconstructions, the interval is the time between two detectable fire episodes, given limitations in temporal resolution and sensitivity of fire proxies.
**Seasonality**: Timing of fire in relation to seasonal cycles (e.g. growing, dry, and rainy).
**Extent**: Spatial size of burned area.
**Cycle**: Time in years for cumulated fire area to equal 100% of the area of interest.
**Type**: Vegetation layer most impacted by flames and heat (e.g. ground, surface, understory, crown).
**Intensity**: Amount of heat released over time per area.
**Severity**: Degree of alteration of vegetation and soil (e.g. mortality %, organic matter combustion %, depth burned).

##### *State Change*

For the purposes of this survey, we define "state change" broadly as a large and sustained departure from a system behavior. State changes can be triggered by smooth or abrupt external or internal drivers (e.g. climate change, disturbances), and can be reversible or permanent. A state change in fire regime can be a shift in central tendency (e.g. significant decrease in mean annual area burned), overall variance (e.g. significant increase in interannual variability of area burned), or frequency of events that exceed some ecologically-relevant threshold (e.g. significant change in the return interval of crown fires). The defining characteristic of **a state change is not the abruptness of the shift, but the magnitude of change in regard to functioning of the system** (Biggs et al., 2009; Scheffer et al., 2001, 2009). Some examples of state changes related to fire:

- In the same climatic conditions, either tropical forest or savanna may develop, depending on feedbacks between vegetation and fire (Staver et al., 2011). A state change in the fire regime can trigger transition from savanna to forest in the case of fire suppression, or forest to savanna in the case of more frequent or severe fire.

- Broadleaf and coniferous trees in the boreal forest are associated with different fire regimes, due to differences in inherent and seasonal flammability. Long-term changes in vegetation have caused multiple state changes in fire due to the interaction of climatological and ecological factors (e.g. Girardin et al., 2013; Higuera et al., 2009; Jasinski & Payette, 2005).
- The transition from temperate woodlands to grasslands is driven by fire regime (Pausas, 2015) , which has experienced multiple state changes due to intentional and unintentional human management (Valkó et al., 2016).

Visual examples of state change in fire regimes:

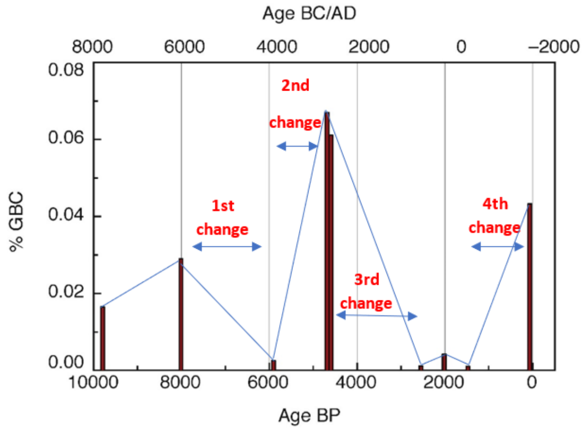

**Figure 1|** Adapted from (Anderson & Wahl, 2016). Graphite Black Carbon (GBC) percent vs. time at Lago Paixban, Guatemala.

---

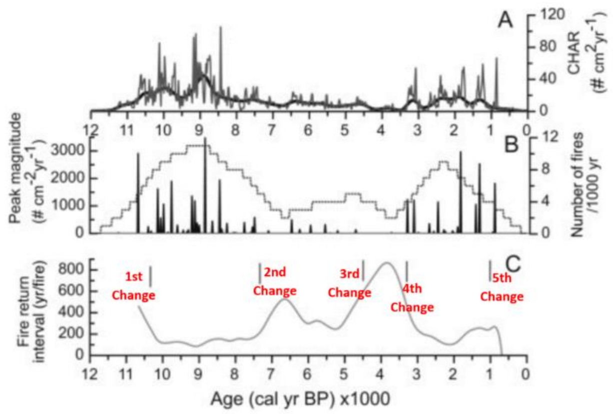

**Figure 2|** Changes in charcoal and fire frequency and magnitude from (Feurdean et al., 2013). "**A)** Interpolated macroscopic charcoal accumulation rate (CHAR; grey curve) and background CHAR (black curve); **B)** inferred number of fires/1000 years (grey dashed curve) and peak magnitude (vertical lines), and **C)** inferred fire return interval/1000 years. Vertical grey lines denote statistically significant zones."

### 3- Questionnaire instructions

You will be asked to provide estimates about past and current fire regime, possible response of fire regime to different climate scenarios, and then conclude with your opinion about fire management. For the purposes of this expert assessment, we are asking for regional estimates, and have partitioned the globe into nine continental regions and 14 eco-regions or biomes (Olson et al., 2001) that reflect bioclimatic, socioeconomic, and fire regime characteristics **(Figure 7)**. We recognize that this partitioning is imperfect and may group multiple conflicting fire regimes. Given the large spatial and temporal scales of interest, please consider the overall response of the region and biome.

For the fire projection section of the questionnaire (section 5.3), we ask for estimates for three warming scenarios (RCP2.6, RCP4.5 and RCP8.5) from the IPCC *AR5 Synthesis Report*, **Figures 3 - 6**). We will ask for estimates over short (Present-2050), medium (Present-2100), and long (Present-2300) time frames. Climate projections and estimates of system response become increasingly uncertain for distant time frames. However, because fire regime can take many decades or centuries to fully respond to disturbance, we have included the 2300-timestep to account for lags in this response. You may think of this third timestep conceptually, e.g. the eventual fire regime/ecosystem state if the described climate conditions persisted.

For each fire region you choose, you will have a chance to indicate your level of confidence and expertise concerning your answer, make comments on how you selected your estimates, and identify key sources of uncertainty concerning the future response of the system (e.g. what data or processes missing from current understanding would most improve our ability to predict system behavior). If there is not yet clear supporting evidence in the literature, but you have some basis for an estimate based on professional judgment, please make a note of that. These supporting questions allow us to compare responses from multiple experts and are just as valuable as your quantitative estimates.

***Confidence level***

The five-point "**Confidence level**" scale is defined as follows:

- **1 =** My answer is my best guess, but I am not confident in it; it could easily be far off the mark.
- **2 =** My answer is an educated guess; it could be far off the mark, but I have some confidence in it.
- **3 =** I am moderately confident in my answer; it is not precise, but it may be near the true value.
- **4 =** I am confident in my answer; the true value is likely to be somewhat different from my answer, but it is unlikely to be dramatically different.
- **5 =** Given current understanding, I would be surprised if my answer were far off from the true value.

***Expertise level***

The five-point "**Expertise level**" scale is defined as follows:

- **1 =** I have little familiarity with the literature, and I do not actively work on this area.
- **2 =** I have some familiarity with the literature, and I've worked on related questions but haven't contributed to the literature on this issue; it is not an area of central expertise for me.
- **3 =** I have worked on related issues and have contributed to the relevant literature but do not consider myself an expert on this issue.
- **4 =** I am very familiar with relevant literature and have worked on related questions. This is an area of central expertise for me.
- **5 =** I contribute actively to the literature directly concerned with this issue, and I consider myself one of the foremost experts on it.

### 4- Climate and disturbance scenarios

This section provides background information on the climate change scenarios and land use changes referenced in the "Fire projections" section of the questionnaire. There is a large body of literature on the scenarios (e.g. *IPCC*), which you are encouraged to draw on as needed for your region. Though the scenarios are largely defined by changes in temperature and precipitation, we encourage you to consider all climatic, ecological, and societal changes for the scenarios that could influence fire regime.

**Table.1** RCP scenarios. From (Moss et al., 2010)

| Scenario | Radiative forcing | $CO_2$-equiv (ppm) concentration | Pathway |
|---|---|---|---|
| RCP8.5 | >8.5Wm$^{-2}$ in 2100 | >1370 in 2100 | Rising |
| RCP4.5 | ~4.5Wm$^{-2}$ at stabilization after 2100 | ~650 at stabilization after 2100 | Stabilization without overshoot |
| RCP2.6 | Peak at ~3Wm$^{-2}$ before 2100 and then declines | 490 before 2100 and then declines | Peak and decline |

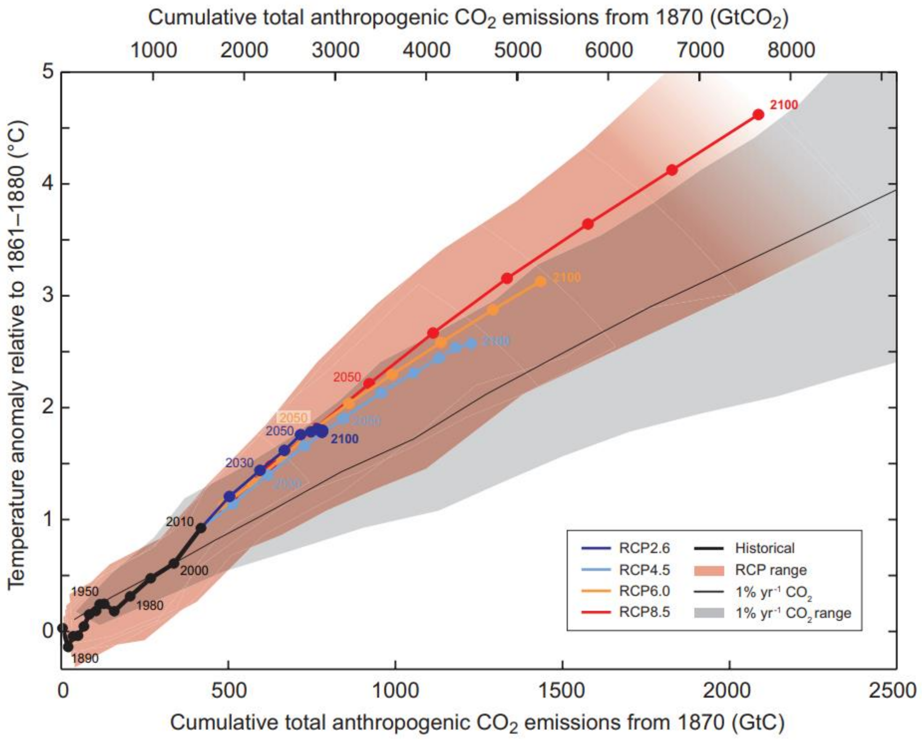

**Figure 3|** Global mean surface temperature change compare to pre-industrial time (1861-1880) using different RCP scenarios. Each scenario is showed with a colored line and decadal mean(dots). From IPCC WG1AR5_Summary for policy makers, figure SPM10, page 28.

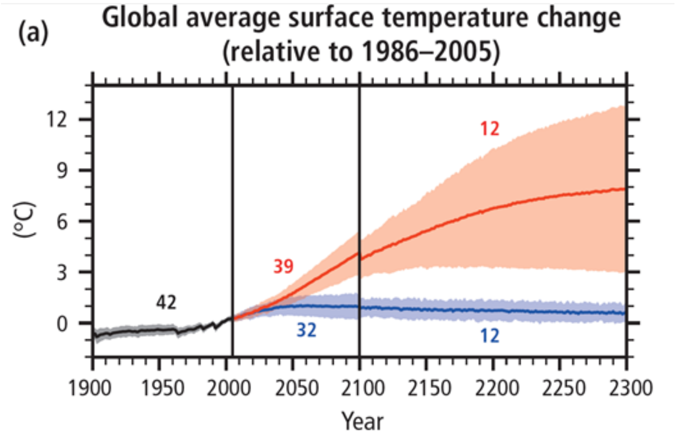

(b)

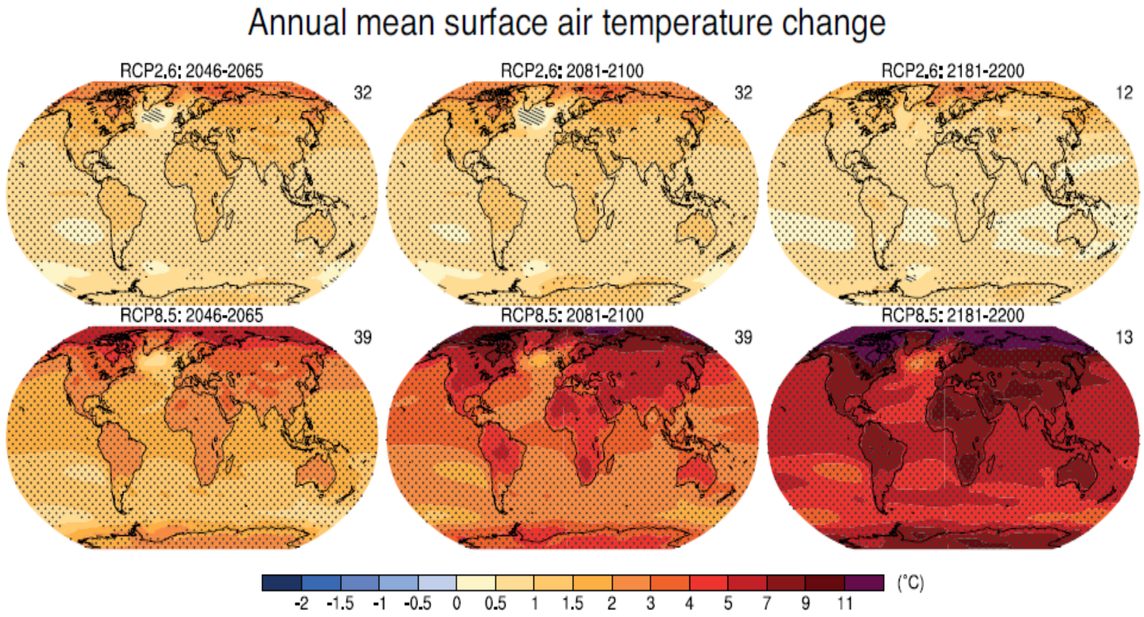

**Figure 4|** Time series of global annual change in mean annual surface temperature for the 1900–2300 period (relative to 1986–2005) from Coupled Model Intercomparison Project Phase 5 (CMIP5). **(a)** Time series of projections and a measure of uncertainty (shading) are shown for scenarios RCP2.6 (blue) and RCP8.5 (red). The number of CMIP5 models used to calculate the multi-model mean is indicated. Projections are shown for the multi-model mean (solid lines) and the 5 to 95% range across the distribution of individual models (shading). Discontinuities at 2100 are due to different numbers of models performing the extension runs beyond the 21st century and have no physical meaning. Modified from IPCC AR5 Synthesis report, figure 2.1, page 59 **(b)** Projected change in average surface temperature for 2081–2100 and 2181-2200 relative to 1986-2005 under the RCP2.6 (top) and RCP8.5 (bottom) scenarios based on multi-model mean projections. Stippling (i.e. dots) shows regions where the projected change is large compared to natural internal variability, and where at least 90% of models agree on the sign of change. Hatching (i.e. diagonal lines) shows regions where the projected change is less than one standard deviation of the natural internal variability. Figures modified from IPCC WG1AR5 chapter 12 ,figure 12.11, page 1063. For detail on the scenarios and additional visualizations, visit: https://ar5-syr.ipcc.ch/topic_futurechanges.php

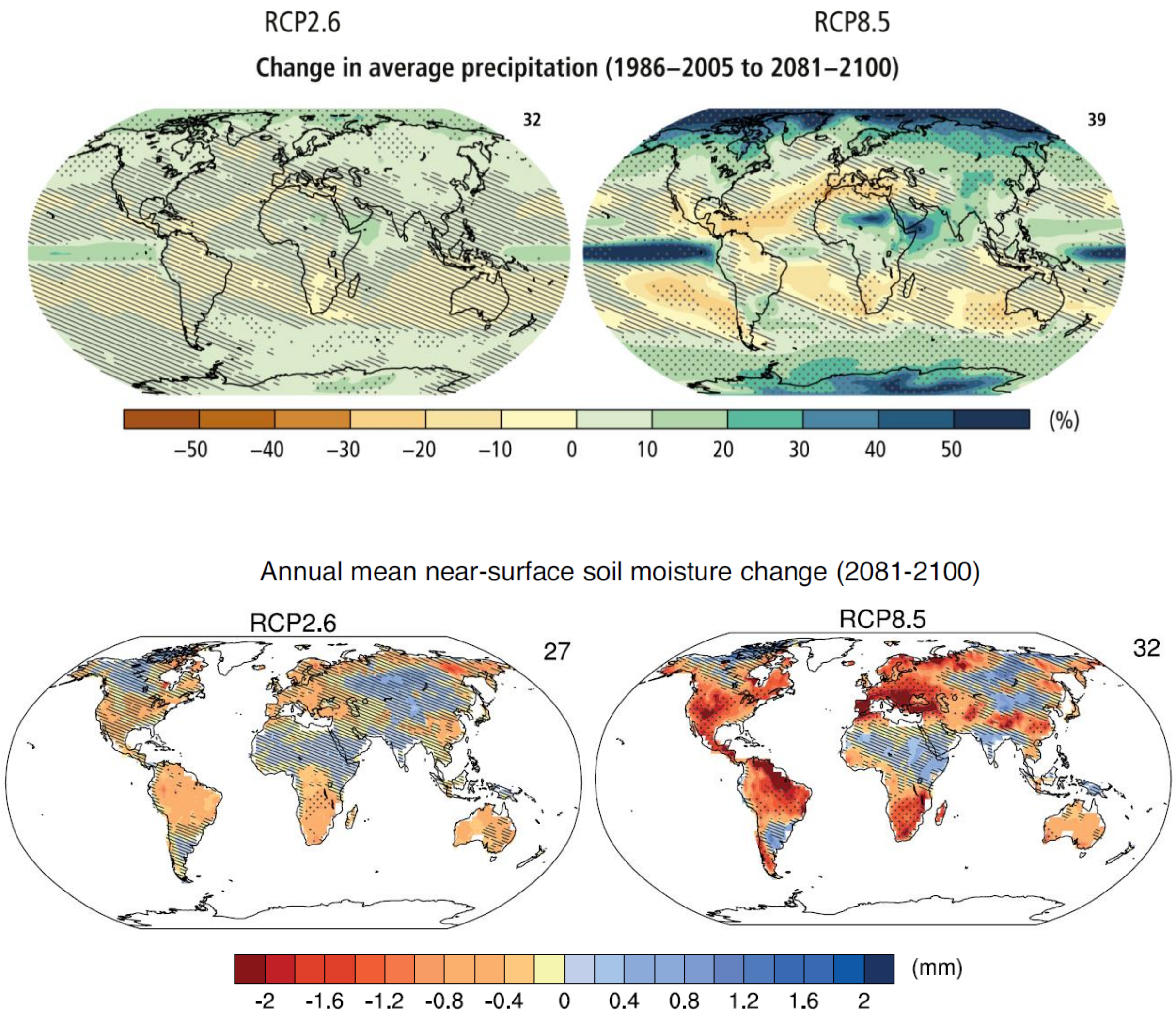

**Figure 5|** Projected change in **(a)** annual mean precipitation Modified from IPCC AR5 Synthesis report, figure 2.2, page 61 and **(b)** annual mean soil moisture in the top 10 cm for 2081-2100 (relative to 1986–2005) for RCP2.6 and RCP8.5. Modified from the IPCC WG1AR5 chapter 12 ,figure 12.23, page 1080. For detailed projections, including seasonal temperature and precipitation trends, relative humidity, temperature return intervals, and more visit: http://www.climatechange2013.org/images/report/WG1AR5_Chapter12_FINAL.pdf

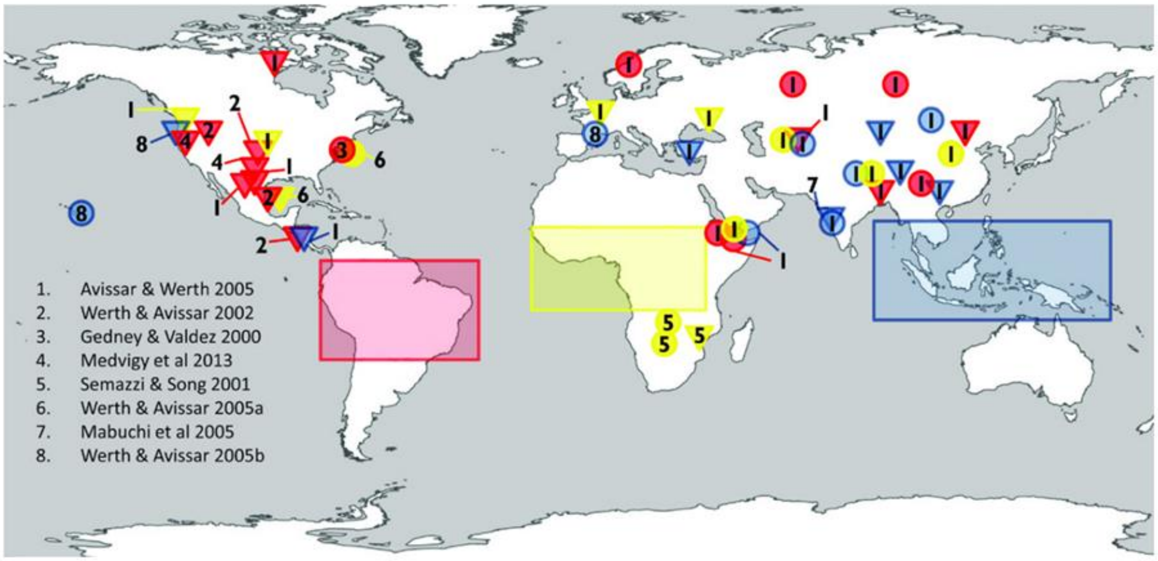

**Figure 6|** An example of links between land-use/direct human disturbance and ecological conditions that could influence fire regime. Extra-tropical effects on precipitation due to deforestation in each of the three major tropical regions. The circles indicate increasing and the triangles indicate decreasing of precipitation as a result of deforestation of the three areas shown with boxes in the figure, including: Red(Amazonia); Yellow(Africa); Blue (Southeast Asia) reviewed by (Lawrence & Vandecar, 2015). From IPCC Climate change and Land_Chapter 2, figure 2.23, page 185.

### 5- Questionnaire

| Respondent background information |
|---|
| First and Last Name |
| Gender |
| Primary research discipline |
| Secondary research discipline |
| Rate yourself on a scale of 1 to 5 where 1 is exclusively field research and 5 is exclusively modeling research. |
| Country of origin |
| Country of employment |
| Years of experience in fire research |

#### 5.1 Selecting region

Please fill out the questionnaire for each region and biome combination (hereafter "fire region") for which you are qualified. If you are qualified to answer for multiple fire regions, please copy the questionnaire from this point on and paste it at the bottom of the document as many times as needed.

| |
|---|
| **Region name** (9 options listed in Fig. 7, e.g. Afrotropic, Australasia, Est Palearctic, Neotropic, etc.) |
| **Biome name** (15 options listed in Fig. 7, e.g. Tundra, Mangroves, Temperate conifer forests, etc.) |
| **Number of years' experience in this area** |
| **Regional expertise rating** (on a scale of 1 to 5, see *Expertise level*) |

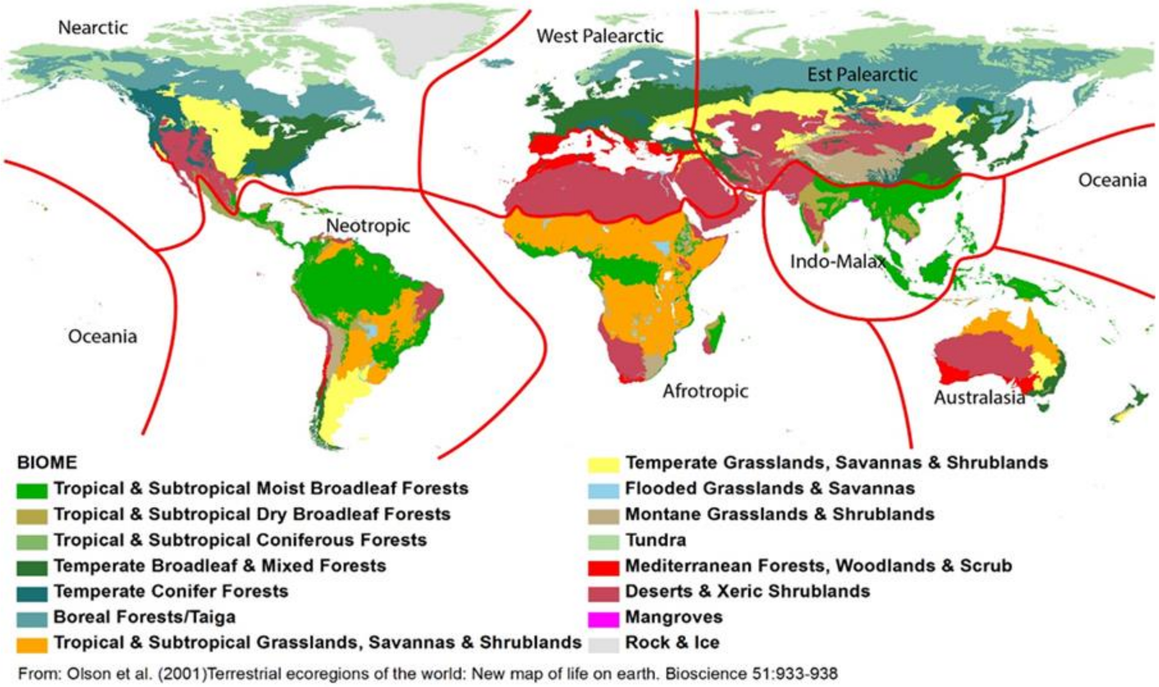

**Figure 7**| Regional partition of the globe by continents and biomes from Olson et al. (2001), "Terrestrial ecoregions of the world: New map of life on earth."

#### 5.2 Paleo perspective

**1.** How many fire regime state changes have occurred in your selected fire region over the last 12,000 years (e.g. during the Holocene)? (See ***State Change*** for background)

| Number of fire regime state changes |
|---|

**2.** When did the three largest of these state changes occur?

| Fire regime state | Year before present | Name or description of the event |
|---|---|---|
| Change 1 | | |
| Change 2 | | |
| Change 3 | | |

**3.** What were the primary drivers of these state changes?
**Note:** List up to three drivers in order of relative importance.

| Fire regime state | State change 1 | State change 2 | State change 3 |
|---|---|---|---|
| Driver 1 | | | |
| Driver 2 | | | |
| Driver 3 | | | |

**4.** Which aspects of the fire regime in this region did post-industrial society influence most strongly?
**Note:** Please identify up to three aspects (see introduction) and how they were influenced (provide as much detail as you wish).

| |
|---|
| Aspect 1 |
| Aspect 2 |
| Aspect 3 |

**5.** How influential is past fire behavior on current fire regimes?

| Percentage (0% = no skill or utility at predicting future behavior, 100% equals deterministic relationship) |
|---|

**Please indicate your expertise and confidence related to section 5.2 Paleo perspective**

| **Average Expertise level (1lowest-5 highest)** | | **Average Confidence level (1lowest-5 highest)** |
|---|---|---|
| **How did you generate these estimates (mark with "x" all that apply)?** | | **What are the largest sources of uncertainty in these estimates?** |
| a) published empirical data:<br>b) published model estimates:<br>c) unpublished data:<br>d) professional opinion:<br>other (please specify): | | |
| **Additional comments:** | | |

#### 5.3 Current fire regime state

**Note:** These questions seek to characterize recent fire behavior in your region. They establish a baseline for the next section. For many of the questions that follow, you will be asked to provide “Lower,” “Central,” and “Upper” estimates, constituting a qualitative 90% confidence interval (see diagram below).

There is a 90% probability the real value is within this range.

**Lower** =
There is a 95% chance that the real value will be greater than this estimate

**Central** =
This is my best estimate

**Upper** =
There is a 95% chance that the real value will be lower than this estimate

**6.** How long has the current fire regime persisted in your selected region?
**Note**: For this question, if there had been a state change in fire regime for your region in the 1950s or 60s, you might answer , “60”, “70”, and “85” in the 3 columns.

| Duration of current fire regime state (years) | Lower | Central | Upper |
|---|---|---|---|

**7.** What is the mean area burned yearly in your region for the period identified in Question 6?

| Extent burned (% of region burned per year) | Lower | Central | Upper |
|---|---|---|---|

**8.** What is the mean fire return interval in your region for the period identified in Question 6?

| Fire Interval (years) | Lower | Central | Upper |
|---|---|---|---|

**9.** What is the current mean fire severity in your region?
**Note**: While there are multiple definitions of fire severity, please respond with the mean percentage of surface organic matter combusted. If you only know the qualitative severity, you can convert to percentage combustion using the scale below (Miesel et al., 2015).

**Lightly burned** = surface organic matter combustion less than 15%
**Moderately burned** = surface organic matter combustion between 15% to 60%
**Severely burned** = surface organic matter combustion more than 60%

| Surface organic matter combustion (%) | Lower | Central | Upper |
|---|---|---|---|

**Please indicate your expertise and confidence related to section 5.3 Current fire state**

| **Average Expertise level (1lowest-5 highest)** | | **Average Confidence level (1lowest-5 highest)** |
|---|---|---|
| **How did you generate these estimates (mark with “x” all that apply)?** | | **What are the largest sources of uncertainty in these estimates?** |
| a) published empirical data:<br>b) published model estimates:<br>c) unpublished data:<br>d) professional opinion:<br>other (please specify): | | |
| **Additional comments:** | | |

#### 5.4 Fire projections

**10.** What is the likelihood of a state change in fire regime for the following global warming scenarios and time steps?

**Note:** Please provide a lower, central, and upper estimate in percent (0 = no possibility of state change, 100 = assured state change) for each time step. Estimates for all three time-steps should be compared with the current fire regime you defined in the last section. In addition to climatic factors, please consider all possible changes for each scenario and time step, including human actions, ecosystem responses, etc.

| Your estimate in % | | | | | | | | | |
|---|---|---|---|---|---|---|---|---|---|
| Scenario | RCP2.6 | | | RCP4.5 | | | RCP8.5 | | |
| Year / CI | 2050 | 2100 | 2300 | 2050 | 2100 | 2300 | 2050 | 2100 | 2300 |
| Lower | | | | | | | | | |
| Central | | | | | | | | | |
| Upper | | | | | | | | | |

**11.** If a change occurs in the fire regime, in what direction would fire extent (area), frequency, and severity likely trend/change for the following global warming scenarios and time steps?

**Note:** Indicate the direction and magnitude for each fire regime dimension by selecting a value between -5 and 5, where -5 = strong decrease, 0 = no change, and 5= strong increase.

| Your estimate between -5 to 5 | | | | | | | | | |
|---|---|---|---|---|---|---|---|---|---|
| Scenario | RCP2.6 | | | RCP4.5 | | | RCP8.5 | | |
| Year | 2050 | 2100 | 2300 | 2050 | 2100 | 2300 | 2050 | 2100 | 2300 |
| Area | | | | | | | | | |
| Frequency | | | | | | | | | |
| Severity | | | | | | | | | |

**12.** What would be the net effect of the fire regime changes you estimated in questions 10 and 11 on biodiversity, carbon stocks, albedo, and ecosystem services (-5 = strong net decrease, 0 = no net effect, 5= strong net increase)?

**Note:** This question focuses on the possible ecological and societal consequences of changes in fire regime. We are interested if the fire regime change will make things better or worse regarding biodiversity (habitat extent, diversity, and quality), climate feedbacks (soil and vegetation carbon stocks and surface albedo), and ecosystem services (other benefits for societies living in this region).

| Your estimate between -5 to 5 | | | | | | | | | |
|---|---|---|---|---|---|---|---|---|---|
| Scenario | RCP2.6 | | | RCP4.5 | | | RCP8.5 | | |
| Year | 2050 | 2100 | 2300 | 2050 | 2100 | 2300 | 2050 | 2100 | 2300 |
| Biodiversity | | | | | | | | | |
| Carbon stocks | | | | | | | | | |
| Albedo | | | | | | | | | |
| Ecosystem services | | | | | | | | | |

**13.** What are the most important drivers of fire regime for the climate scenarios?
**Note:** This question aims to identify how medium and long-term drivers (e.g. climate, vegetation, human activity) interact to determine fire regime over different timescales. For the purposes of this question, we ask you to consider change by the year 2100. List up to 5 fire regime drivers in order of relative importance.

| Fire regime drivers in order of relative importance | | | |
|---|---|---|---|
| Scenario | RCP2.6 | RCP4.5 | RCP8.5 |
| (Most important) 1- | | | |
| 2- | | | |
| 3- | | | |
| 4- | | | |
| (Less important) 5- | | | |

**Please indicate your expertise and confidence related to section 5.4 Fire Projections**

| Average Expertise level (1lowest-5 highest) | | Average Confidence level (1lowest-5 highest) |
|---|---|---|
| **How did you generate these estimates (mark with "x" all that apply)?** | | **What are the largest sources of uncertainty in these estimates?** |
| a) published empirical data:<br>b) published model estimates:<br>c) unpublished data:<br>d) professional opinion:<br>other (please specify): | | |
| **Additional comments:** | | |

#### 5.5 Intervention and management

**14.** What human actions regarding fire would be most effective in preserving or enhancing the following values over the next 20-50 years?
**Note:** Please note that "Non-intervention" is also an action. Indicate up to 5 actions. For each action indicate the degree of effect on a scale of -5 to 5, where -5 = most damaging or counterproductive, 0 = no effect, and 5 = most preserving or enhancing. (**Ecosystem services** here refer to all provisioning, regulating, social and cultural services that forests provide for humans.)

| Human actions regarding fire | Your estimate between -5 to 5 | | | |
|---|---|---|---|---|
| **Human action** | **Biodiversity** | **Carbon Stocks** | **Albedo** | **Other ecosystem services** |

**15.** How effective could human interventions be in mitigating potential damage to societies and ecosystems for the following scenarios and time periods?
**Note:** This question seeks to assess humans' ability to manage fire under increasing climatological and ecological forcing of fire regime. For example, in the future will environmental changes systematically surpass human's technical and social abilities to control fire regime, or does human capacity grow as fast or faster as environmental drivers? Indicate your response on a scale of -5 to 5 compared to current conditions, where -5 = much less capacity to control fire regime for this scenario and period, 0 = same capacity as today, and 5 = much stronger capacity.

| **Your estimate between -5 to 5** | | | | | | | | | |
|---|---|---|---|---|---|---|---|---|---|
| **Scenario** | **RCP2.6** | | | **RCP4.5** | | | **RCP8.5** | | |
| **Year** | **2050** | **2100** | **2300** | **2050** | **2100** | **2300** | **2050** | **2100** | **2300** |

**Please indicate your expertise and confidence related to section 5.5 Intervention and management**

| **Average Expertise level (1lowest-5 highest)** | | **Average Confidence level (1lowest-5 highest)** |
|---|---|---|
| **How did you generate these estimates (mark with "x" all that apply)?** | | **What are the largest sources of uncertainty in these estimates?** |
| a) published empirical data:<br>b) published model estimates:<br>c) unpublished data:<br>d) professional opinion:<br>other (please specify): | | |
| **Additional comments:** | | |

| What are the three most important scientific citations/studies about fire in your selected region. |
|---|
| Reference 1 |
| Reference 2 |
| Reference 3 |
